# Improving split reporters of protein-protein interactions through orthology-based protein engineering

**DOI:** 10.1101/2023.08.31.555680

**Authors:** Louise-Marie Rakotoarison, Alison G. Tebo, Dorothea Böken, Arnaud Gautier

## Abstract

Protein-protein interactions (PPI) can be detected through selective complementation of split fluorescent reporters made of two complementary fragments that reassemble into a functional fluorescent reporter when in close proximity. We previously introduced splitFAST, a chemogenetic PPI reporter with rapid and reversible complementation. Here, we present the engineering of RspA-splitFAST, an improved reporter displaying higher brightness, lower self-complementation and higher dynamic range for optimal monitoring of PPI using an original protein engineering strategy that exploits proteins with orthology relationships. Our study allowed the identification of a system with improved properties and enabled a better understanding of the molecular features controlling the complementation properties. Because of the rapidity and reversibility of its complementation, its low self-complementation, high dynamic range, and improved brightness, RspA-splitFAST is well suited to study PPI with high spatial and temporal resolution, opening great prospects to decipher the role of PPI in various biological contexts.

## INTRODUCTION

Molecular reporters allowing the visualization of protein-protein interactions (PPI) have become an essential tool in biological research and drug discovery. They can enable the study of the function and dynamics of PPI in a given biological process or the identification of specific molecules able to activate or inhibit PPI. Selective optical detection of PPI can be achieved through complementation of split reporters generated by bisection of protein-based fluorescent or bioluminescent reporters into two complementary fragments able to reassemble into a functional reporter when in close proximity [1, 2]. Our lab recently developed a PPI reporter [3] through bisection of FAST (Fluorescence-Activating and absorption-Shifting Tag), a small chemogenetic reporter that allows monitoring of gene expression and protein localization in live cells and organisms [4, 5]. Evolved from the bacterial photoreceptor photoactive yellow protein (PYP) from *Halorhodospira halophila*, FAST is a small protein tag that binds and stabilizes the fluorescent state of hydroxybenzylidene rhodanine chromophores (HBR) (Li et al., 2017; Plamont et al., 2016; Tebo et al., 2018). Dark when free in solution or cells, these fluorogenic chromophores (so-called fluorogens) allow the imaging of FAST-tagged proteins with very high contrast without the need for washing. FAST and its variants have proved to be a useful tool that is compatible with multiple microscopy modalities and model organisms [4, 6-8]. In particular, they excel in applications in oxygen-poor environments [9-11] or in situations where the lack of delay in the formation of a fluorescent complex allows the detection of rapid biological events, such as cell cycles during early zebrafish development [7]. Bisection of FAST allowed the generation of splitFAST, a bimolecular fluorescence complementation system for detecting PPI [3]. As fluorogen binding is non-covalent, the complementation of splitFAST is rapid and reversible allowing the visualization of dynamic PPI. The original splitFAST suffered however from lower brightness than the parent protein:fluorogen complex and some detectable background signal due to self-complementation.

Here, we described the use of an original orthology-based protein engineering approach to generate an improved splitFAST with low self-complementation and improved brightness. Given the tripartite nature of splitFAST, optimization using the strategies employed to expand and enhance genetically encoded fluorescent probes was challenging. The difficulty came from the challenge of predicting the functional effect of an amino acid change on the brightness and/or the self-complementation of the system. Moreover, the necessity of possibly changing amino acids in both fragments rendered the exploration of combinatorial libraries by directed evolution extremely arduous. In order to sample broad sequence space and functionalities, we took advantage of the natural sequence homology and diversity found in members of a protein family. Members in a protein family can be separated into orthologs, when they are derived from speciation, or paralogs when they result from gene duplication. Orthologous proteins are particularly attractive for protein engineering as they are assumed to have the same function and usually the same specificity in close organisms, while displaying various degrees of sequence homology. We thus hypothesized that the sequence diversity of proteins with orthology relationships could generate various properties while keeping the overall function. Here, we describe the use of orthologous proteins to develop an improved splitFAST system for the detection of PPI with high spatial and temporal resolution.

## RESULTS

### Engineering new FASTs using orthologs of PYP

Prototypical FAST differs from *Halorhodospira halophila* (Hha)-PYP by the mutation C69G and the sequence of the loop 94-101 that gates the chromophore binding pocket, which is WMIPTSRG in FAST rather than YTFDYQMT in Hha-PYP [4]. FAST was isolated by yeast cell surface screening from a large library of *Halorhodospira halophila* (Hha)- PYP-C69G variants randomized by saturation mutagenesis in the loop 94-101. Given that Hha-PYP is the prototypical member of the PYP family of proteins, of which there are now over 140 members [12], we explored the use of PYP orthologs to generate new proteins with FAST behavior. Classified into seven classes based on percent identity, shared insertions or deletions, and conserved residues, PYP orthologs can share between 25 to 78 % amino acid sequence identity and display distinct functions. The class to which Hha-PYP belongs has 17 other members all derived from proteobacterial strains. These proteins share 70-78% amino acid sequence identity with Hha-PYP with no insertions or deletions. We hypothesized that the sequence WMIPTSRG found in the loop 94-101 could confer FAST-like behavior to PYP orthologs. To test this hypothesis, we selected six orthologs of PYP exhibiting between 70-78% amino acid sequence identity (**Fig. 1b**). We replaced the chromophore-binding cysteine C69 by a glycine, and the 94-101 loop with the sequence WMIPTSRG (present in FAST) in orthologs from *Halomonas boliviensis* LC1 (HboL), *Halomonas sp.* GFAJ-1 (HspG), *Rheinheimera sp.* A13L (RspA*), Idiomarina loihiensis* (Ilo), *Thiorhodospira sibirica* ATCC 700588 (TsiA) and *Rhodothalassium salexigens* (Rsa). We heterologously expressed the resulting constructs in *E. coli*, purified them and screened them for HMBR binding. Of the six proteins that were tested, all expressed acceptably well for screening, were able to bind HMBR, and were thus characterized in further detail.

**Figure 1.**
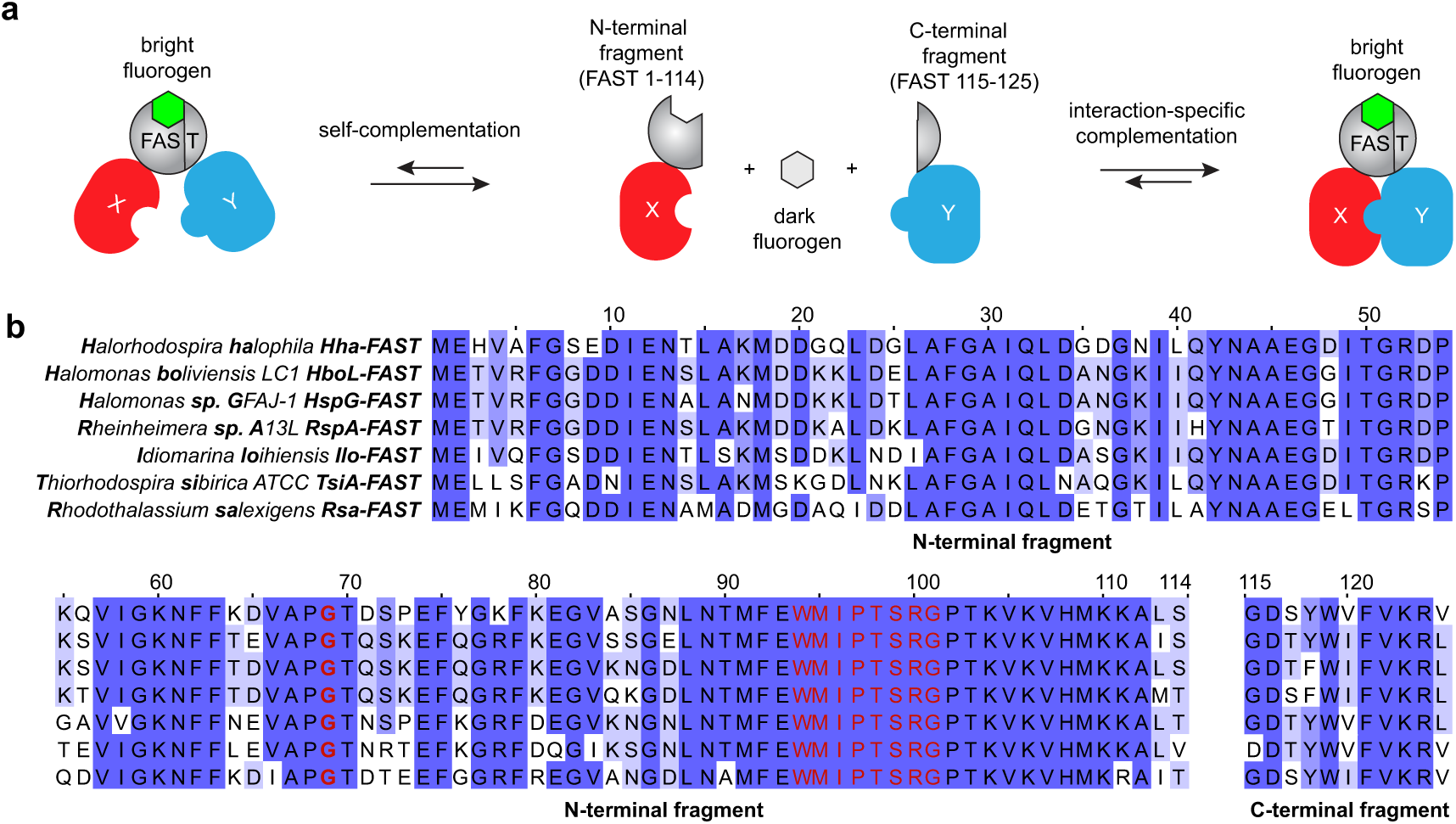
Improvement of split-FAST through orthology-based engineering. **a** Bisection of FAST between residues 114 and 115 allowed the creation of a split fluorescent reporter with rapid and reversible complementation. When in close proximity, the two fragments reconstitute into a functional bipartite FAST protein able to bind and stabilize the fluorescent state of fluorogenic dyes (so-called fluorogens). An optimal system should present efficient interaction-dependent complementation and minimal self-assembly to allow the acute visualization of PPI. **b** Sequences of the seven ortholog-based split reporters generated in this study. In red are indicated the residues inserted for generating FAST behavior.

The affinity for the fluorogen and the quantum yield of the six proteins were measured with the three fluorogens, HMBR, HBR-3,5DM, HBR-3,5DOM, which emit between 540 nm and 600 nm when bound to FAST [13, 14]. All of the six proteins formed fluorescent complexes with the tested fluorogens (**Table S1**). RspA-FAST and TsiA-FAST displayed improved affinity for the fluorogens, including for HBR-3,5DOM (*K*_D_ = 74 and 60 nM), which typically binds FAST variants with ∼ 1 μM affinity [7, 13, 14]. Furthermore, several complexes displayed higher fluorescence quantum yields than FAST: in particular, HspG-FAST and Ilo-FAST with HMBR and RspA-FAST and Ilo-FAST with HBR-3,5DOM (**Table S1**). Thus, despite large sequence variability, the mutated orthologs of PYP are types of FASTs. Because of their conserved functions and the orthology relationship between their parental PYP proteins, the six generated proteins are called hereafter FAST orthologs. To test whether the FAST orthologs were also functional in live cell imaging, we expressed them in HEK293T cells with mTurquoise2 as a transfection marker. All 18 ortholog-fluorogen pairs were viable fluorescence imaging tags, displaying distribution similar to mTurquoise2 and no visible aggregates (**Fig. S1**).

### PYP orthologs as a way to improve splitFAST

Improvement of complementation assays is a non-trivial protein engineering task due to the challenge of optimizing two protein chains that must efficiently re-fold together. In the case of Hha-splitFAST, optimization must also consider fluorogen binding. Given the difficulty of improving these systems directly, we reasoned that PYP orthologs would allow us to sample a high, yet tolerated, protein sequence diversity. Furthermore, as the mutated PYP orthologs perform as FASTs, we hypothesized that they would also function as splitFASTs. To evaluate whether the FAST orthologs could be used to generate improved split fluorescent reporters, we split each of them into two complementary fragments in their last loop between Ser114 and Gly115 (**Fig. 1b**), as was previously done to generate Hha-splitFAST [3].

To evaluate the complementation of the orthologous splitFASTs, we developed a flow cytometry assay based on the rapamycin-induced association of the FK506-binding protein (FKBP) and the FKBP-rapamycin-binding (FRB) domain of mechanistic target of rapamycin (mTOR) [15, 16]. Self-and interaction-dependent complementation could be evaluated by measuring the fluorescence response of the splitFAST orthologs without and with rapamycin (**Fig. 2a**). For each orthologous protein, we fused the C-terminal fragment 115-125 to the C-terminus of FKBP and the N-terminal fragment 1-114 to the C-terminus of FRB. FKBP and FRB fusions were co-expressed in HEK293T cells using two bicistronic vectors that allowed also the expression of iRFP670 and mTurquoise2, respectively, as transfection reporters. In order to compare the different systems, we quantified cell fluorescence by flow cytometry in the absence and presence of rapamycin at various HMBR concentrations (**Fig. S2**).

**Figure 2.**
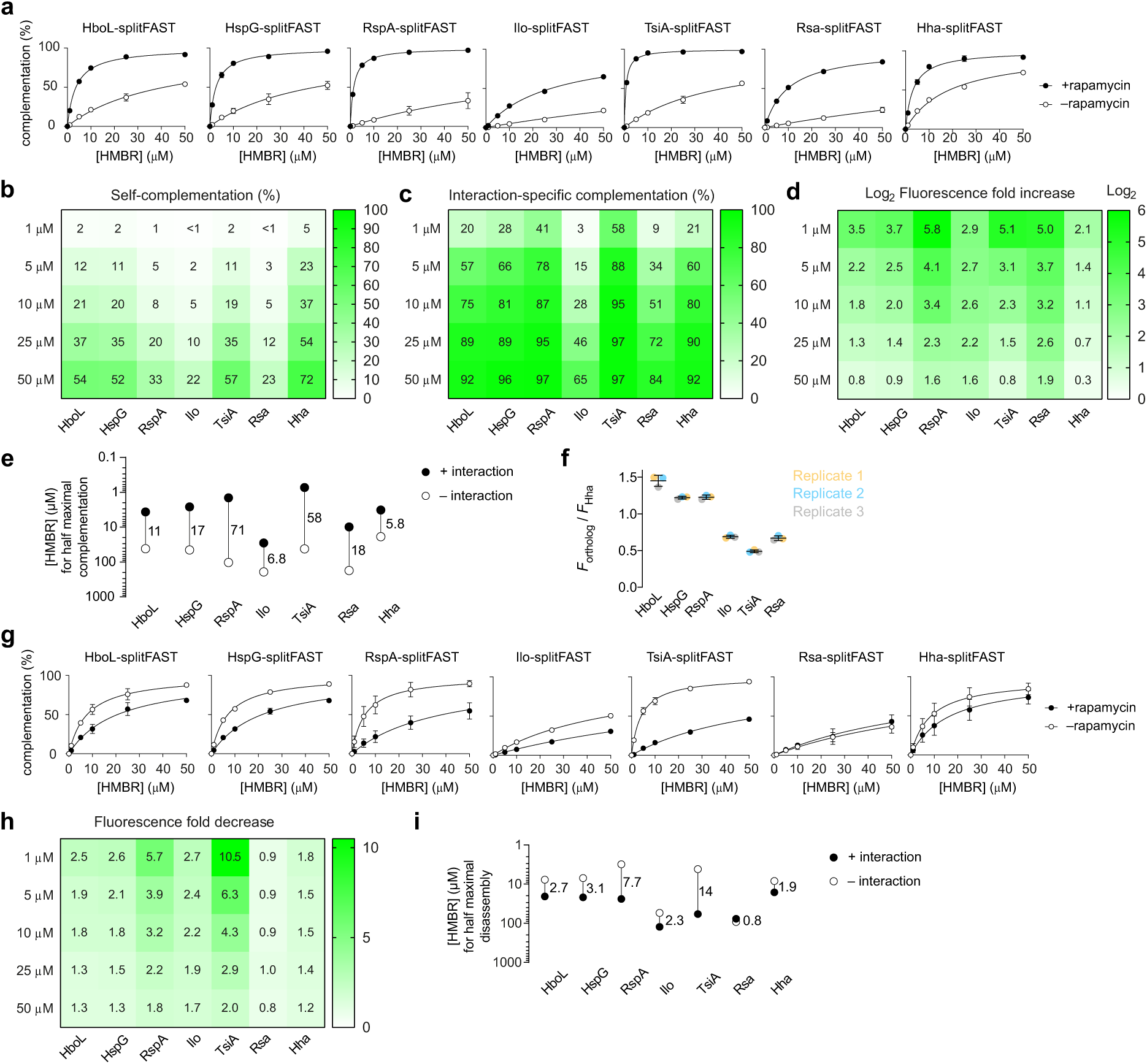
Flow cytometry analysis of the ortholog-based split reporters. **a** Normalized average fluorescence of about 50,000 HEK293T cells co-expressing the FK506-binding protein (FKBP) fused to the C-terminal fragment of the split reporters and the FKBP-rapamycin-binding domain of mammalian target of rapamycin (FRB) fused to the N-terminal fragment of the split reporters treated without or with 500 nM of rapamycin, and with 1, 5, 10, 25 or 50 μM of HMBR (see **Fig. S2**). Data represent the mean ± standard deviation of two independent experiments. These data allowed to determine for each split reporter **b** the percentage of self-complementation, **c** the percentage of interaction-specific complementation and **d** the fluorescence fold increase (dynamic range) at various fluorogen concentrations. **e** Concentration of fluorogen for half maximal complementation in presence (EC_50,+interaction_) or absence (EC_50,–interaction_) of rapamycin (see also **Table S2**). The ratio EC_50,–interaction_ / EC_50,+interaction_ is indicated for each ortholog-based split reporter. **f** Relative brightness of the ortholog-based split reporters at 50 μM of HMBR in presence of rapamycin from three independent experiments. Each biological replicate is color-coded. **g** Normalized average fluorescence of about 50,000 HEK293T cells co-expressing the homodimerizing FK506-binding protein F36M mutant (FKBP_F36M_) fused to the N-terminal and C-terminal fragments of the split reporters treated without or with 500 nM of rapamycin, and with 1, 5, 10, 25 or 50 μM of HMBR. Data represent the mean ± standard deviation of two independent experiments. These data allowed to determine for each split reporter **h** the fluorescence fold decrease at various fluorogen concentrations. **i** Concentration of fluorogen for half maximal complementation in presence (EC_50,–interaction_) or absence (EC_50,+interaction_) of rapamycin (see also **Table S4**). The ratio EC_50,+interaction_ / EC_50,–interaction_ is indicated for each ortholog-based split reporter.

For all orthologous split proteins, we first observed that the higher the fluorogen concentration, the lower the dynamic range observed (i.e. the fluorescence increase upon rapamycin addition). This progressive decrease in dynamic range with increasing fluorogen concentrations is due to an increase in the self-assembly of the system (**Fig. 2a-d**), which is consistent with the fluorogen contributing to the stabilization of the interaction of the split protein fragments. Interestingly, we observed that, at a given fluorogen concentration, the dynamic range of all the ortholog-based split reporters was improved relative to the original Hha-splitFAST (**Fig. 2d**), due mainly to lower self-assembly (**Fig. 2b**). The system with the highest dynamic range and lowest self-assembly was the one generated from RspA-FAST: for example, at 5 µM of HMBR, addition of rapamycin induced a 17-fold increase of fluorescence, while only a 3-fold increase was observed for Hha-splitFAST. In order to compare the systems using a parameter independent of the fluorogen concentration used, we determined for each system the effective concentration of fluorogen for half maximum complementation in the absence (EC_50,– interaction_) and in the presence of interaction (EC_50,+ interaction_) (**Fig. 2e** and **Table S2**). The EC_50,– interaction_ value reflects the ability of the system to self-complement in the absence of interaction: the higher the value, the lower the self-complementation. The EC_50,+ interaction_ value on the other hand reflects the efficacy of the interaction-dependent complementation: the lower the value, the more efficient the interaction-dependent complementation. This EC_50,+ interaction_ value also allows an estimation of the apparent binding affinity of the fluorogen for the complemented split system in cells. The EC_50,– interaction_ / EC_50,+interaction_ ratio provides a direct way to quantify and compare the dynamic range of the reporters. RspA-splitFAST appeared to be the system with the highest dynamic range with a ratio value of 71 compared to 5.8 for Hha-splitFAST (**Fig. 2e**). This analysis allowed us to show that the gain in dynamic range observed for RspA-splitFAST was due to both a reduced self-complementation and a more efficient interaction-dependent complementation.

To better evaluate the relative brightness of the splitFAST orthologs, we used the transfection reporters, mTurquoise2 and iRFP670 to select cells with approximately equivalent expression levels. We then compared the fluorescence in the HMBR channel, measured at 50 µM to maximize fluorogen binding, and normalized all measurements to Hha-splitFAST (**Fig. 2f**). We observed that Ilo-splitFAST, TsiA-splitFAST, and Rsa-splitFAST displayed lower brightness (between one-half and three-quarters of the fluorescence of Hha-splitFAST), while HboL-splitFAST, HspG-splitFAST, and RspA-splitFAST displayed improved brightness.

We previously demonstrated that the affinity of the two split fragments could be tuned by truncating the C-terminal fragment, thus increasing the dynamic range by reducing the self-assembly of the system [3]. However, this truncation reduced the overall brightness of the reporter, presumably due to an overall loss of affinity for the fluorogen. We therefore compared RspA-splitFAST with Hha-splitFASTΔC1, the truncated version of splitFAST missing the last residue in its C-fragment (**Fig. S3- S4** and **Table S3**). Although RspA-splitFAST displayed slightly higher self-assembly than Hha-splitFASTΔC1, it showed much higher interaction-dependent complementation efficiency, resulting overall in a higher or comparable dynamic range. Our titration experiments showed that labeling of RspA-splitFAST was optimal at 5-10 μM HMBR, while suboptimal for Hha-splitFASTΔC1 due to a lower fluorogen binding affinity (as evidenced by a much higher EC_50,+ interaction_). We evaluated the performance of RspA-splitFAST with two other fluorogens, HBR-3,5DM and HBR-3,5DOM, which emit at 560 nm and 600 nm, respectively. As with HMBR, we observed that RspA-splitFAST outperformed Hha-splitFAST and Hha-splitFASTΔC1 in terms of dynamic range and/or brightness (**Fig. S3 and Fig. S4**). These results demonstrate that RspA-splitFAST retains the ability to bind multiple fluorogens and that its improved dynamic range is independent of the fluorogen used. Overall these experiments suggested that bisection of RspA-FAST allowed the generation of a split reporter with high dynamic range without compromising on the brightness, outperforming thus both Hha-splitFAST and Hha-splitFASTΔC1.

One of the benefits of the original Hha-splitFAST system was its reversibility, which is not found in other complementation-based fluorescence reporters. We thus examined whether the reversibility was maintained in the novel splitFAST variants. For each ortholog variant, we fused the two split fragments to the C-terminus of FKBP-F36M, a variant of FKBP known to spontaneously assemble into a weak homodimer, which can be disrupted by addition of rapamycin [17]. Cells co-expressing the two fusions were treated with various concentrations of HMBR, and treated with or without rapamycin (**Fig. 2g-i**). Addition of rapamycin led to a loss of fluorescence for all orthologs but Rsa-splitFAST, in agreement with the dissociation of the split reporter upon homodimer disruption. Rsa-splitFAST appeared to poorly complement in this context likely explaining the poor response. Among the tested systems, TsiA-splitFAST and RspA-splitFAST showed the largest variations of fluorescence upon dissociation. To quantitatively compare the systems, we evaluated the ability of the systems to dissociate using the EC_50,+interaction_ / EC_50,–interaction_ ratio. TsiA-splitFAST and RspA-splitFAST displayed a ratio of 14 and 7.7 respectively, compared to 1.9 for Hha-splitFAST, demonstrating their improved ability to dissociate (**Fig 2.i** and **Table S4**). These results agree with the reduction of self-complementation we observed in the association assay. We observed that the dynamic range for dissociation was lower than for association. We attribute this difference to the low affinity of the FKBP-F36M homodimer, which can result in an incomplete complementation before the addition of rapamycin, and therefore to a lower change in fluorescence upon dissociation. For RspA-splitFAST, we observed very similar results when using HBR-3,5DM and HBR-3,5DOM (**Fig. S5** and **Table S5**), in agreement with the results of the association assay. Comparison of RspA-splitFAST with Hha-splitFASTΔC1 in this dissociation assay also confirmed the superior performance of RspA-splitFAST for the detection of protein complex disassembly (**Fig. S5** and **Table S5**). As splitFAST is a three-component system composed of a chromophore and two complementary fragments of FAST, the concentrations of the different components influence the extent of self-complementation and thus the dynamic range. Our analysis showed that increasing the fluorogen concentration increases self-complementation and reduces the dynamic range. To complement this analysis, we evaluated the impact of the protein expression level on the self-complementation and dynamic range. We focused our analysis on RspA-splitFAST, Hha-splitFAST and Hha-splitFASTΔC1 (**Fig. S6**). We used the transfection reporters to select cells with “low” and “high” expression levels. We observed that, for almost all fluorogen concentrations and protein variant combinations, the dynamic range was higher at low expression levels, and that, accordingly, low expression levels led also to less self-complementation.

We further compared RspA-splitFAST, Hha-splitFAST and Hha-splitFASTΔC1 by assessing whether these systems perturb the efficacy of the known PPI modulators. Specifically, we have relied on the ability of rapamycin to induce an interaction between FRB and FKBP and inhibit the FKBP-F36M homodimerization. We determined the EC_50_ of the FRB-FKBP interaction to be ∼45 nM for all three split systems, which is consistent with the measured dissociation constant, *K_D_*, of ∼12 nM for the ternary complex [15] (**Fig. S7a**). In the case of the rapamycin-induced inhibition of FKBP-F36M homodimerization, we determined an effective concentration of rapamycin for half maximum inhibition of around 100 nM for the split reporters (**Fig. S7b**), in agreement with previously reported values [17]. In both cases, the RspA-splitFAST system displayed the highest dynamic range.

### Exploring sequence space through systematic fragment crossing

Considering the strong homology between the sequences of the N-and C-terminal fragments found in the seven FAST orthologs, we decided to pair each N-terminal fragment with each C-terminal fragment in order to explore a wider sequence space. We studied the self-complementation and interaction specific-complementation of the 49 generated split systems in HEK293T cells by flow cytometry using the rapamycin-inducible interaction of FRB and FKBP as previously. Because of the high number of systems to analyze in parallel, the set of conditions have been reduced to only three concentrations of HMBR (0, 10 and 50 µM). This set of conditions allowed us to evaluate for each system the self-and interaction-specific complementation (and thus the dynamic range) at 10 µM of fluorogen (**Fig. 3a-c**). The mean intensity at 50 µM of fluorogen in the presence of rapamycin allowed us to evaluate and compare the relative brightness of the different systems (**Fig. 3d**).

**Figure 3.**
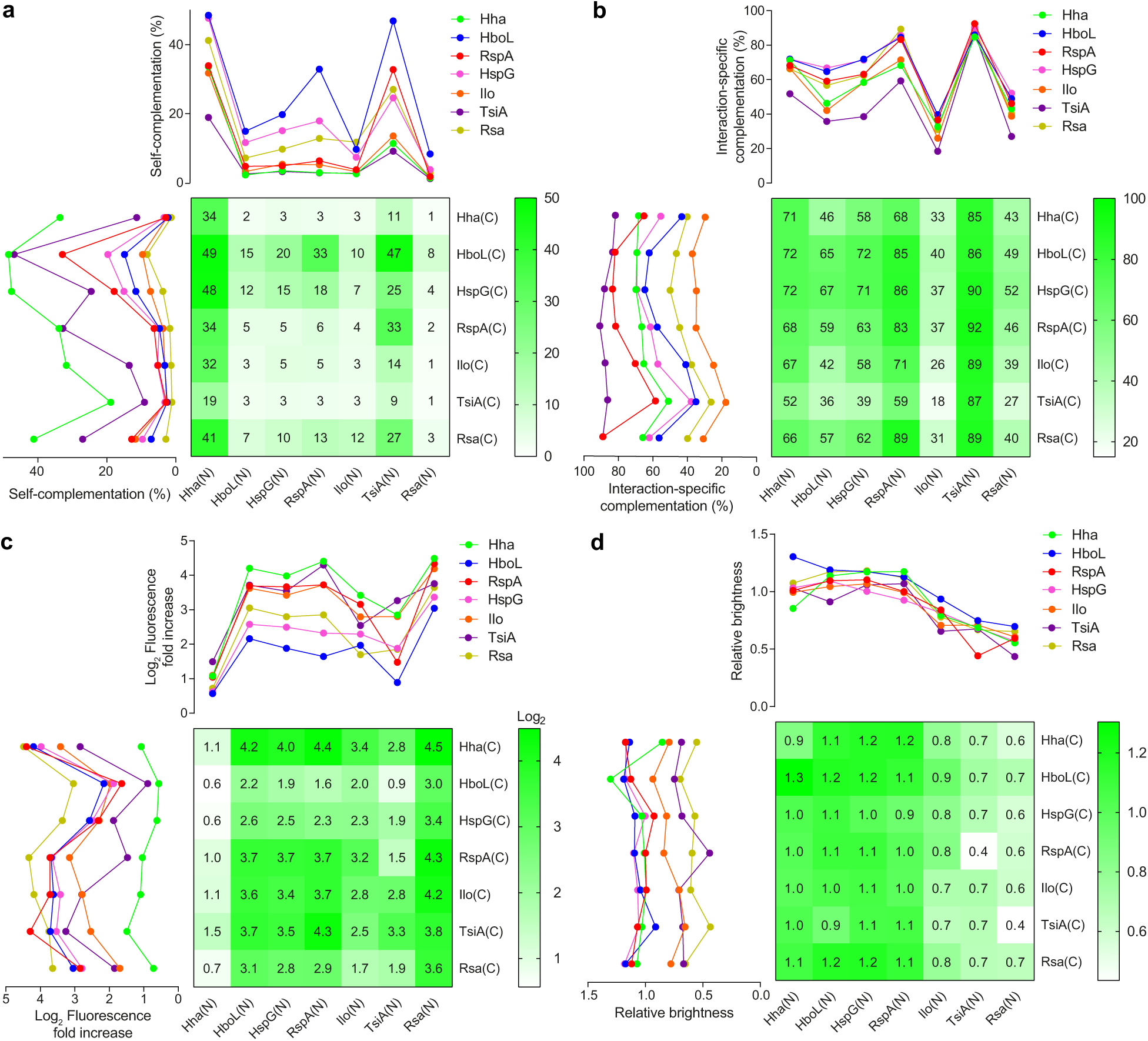
Systematic fragment crossing. HEK293T cells co-expressing the FK506-binding protein (FKBP) fused to the C-terminal fragment of the split reporters and the FKBP-rapamycin-binding domain of mammalian target of rapamycin (FRB) fused to the N-terminal fragment of the split reporters were treated with 500 nM or without rapamycin and with 0, 10 and 50 μM of HMBR. The fluorescence of about 50,000 HEK293T cells was analyzed by flow cytometry for each condition, allowing to estimate for each system **a** the self-complementation, **b** the interaction-specific complementation, **c** the fluorescence fold increase, and **d** the relative brightness. Data represent the mean of two independent experiments.

Our analysis allowed us to extract meaningful information about the role of the N-terminal and C-terminal fragments on the self-and interaction-dependent complementation and the fluorogen binding affinity. With respect to self-complementation, we observed that some N-terminal fragments, such as those of Hha-FAST and, to a lesser extent, TsiA-FAST, systematically led to higher self-complementation, irrespective of the C-terminal fragment used, while others, such as those of HboL-FAST, HspG-FAST, RspA-FAST, Ilo-FAST and Rsa-FAST led systematically to lower self-complementation (**Fig. 3a**). We also observed that the C-terminal fragments of HboL-FAST, HspG-FAST and, to a lesser extent Rsa-FAST led also systematically to higher self-complementation than those of Hha-FAST, RspA-FAST, Ilo-FAST and TsiA-FAST, no matter the N-terminal fragment (**Fig. 3a**).

Analysis of the efficacy of the interaction-dependent complementation showed that the N-terminal fragments of TsiA-FAST, RspA-FAST and Hha-FAST systematically led to high interaction-dependent complementation, while those of Ilo-FAST and Rsa-FAST led to modest or low interaction-dependent complementation (**Fig. 3b**). The highest interaction-specific complementation was observed with the N-terminal fragment of TsiA-FAST, followed by RspA-FAST, and then Hha-FAST. The nature of the C-terminal fragment seems to have less impact on the interaction-dependent complementation: beside the C-terminal fragment of TsiA-FAST that led to lower interaction-dependent complementation in average, all the others showed comparable efficacy.

Analysis of the relative brightness of the different systems showed two clear groups: the systems with the N-terminal fragment of Hha-FAST, HboL-FAST, HspG-FAST and RspA-FAST were brighter, while those with the N-terminal fragments of Ilo-FAST, TsiA-FAST and Rsa-FAST were dimmer (**Fig. 3d**). For a given N-terminal fragment, the brightness was comparable regardless of which C-terminal fragment was used, suggesting that the brightness depends mainly on the sequence of the N-terminal fragment.

Finally, analysis of the dynamic range of the different systems showed that the N-terminal fragment of Hha-FAST led systematically to a lower dynamic range, followed by the N-terminal fragment of TsiA-FAST and Ilo-FAST, in agreement with their higher self-complementation (**Fig. 3c**). The highest dynamic range were obtained with the N-terminal fragments of HboL-FAST, HspG-FAST, RspA-FAST and Rsa-FAST (**Fig. 3c**).

Overall, our analysis showed that RspA-splitFAST was the best compromise for minimizing the self-complementation, while optimizing the efficacy of interaction-dependent complementation, the dynamic range and the brightness. This analysis also pointed out the important role of the N-terminal fragment on the self-complementation, the dynamic range and the brightness of the systems. In particular, our results suggest that the lower self-complementation and higher dynamic range of RspA-splitFAST compared to Hha-splitFAST could be mainly due to the changes in their N-terminal fragments. This was supported by the fact that, not only did the N-terminal fragment of RspA-FAST and Hha-FAST systematically lead, respectively, to lower and higher self-complementation (or higher and lower dynamic range) no matter the paired C-terminal fragment, but also because crossing the fragments of Hha-splitFAST and RspA-splitFAST generated two new systems with opposite properties: RspA(N)::Hha(C) with low self-complementation and high dynamic range (as in RspA-splitFAST), and Hha(N)::RspA(C) with high self-complementation and low dynamic range (as in Hha-splitFAST). To better understand the role of the N-and C-terminal fragments on the complementation, we characterized the association and dissociation of the crossed systems RspA(N)::Hha(C) and Hha(N)::RspA(C) at various fluorogen concentrations as previously done for characterizing the six orthologous split systems. The full analysis of the association experiments confirmed our initial observation: RspA(N)::Hha(C) showed very low self-complementation, even lower than RspA-splitFAST in absence of interaction, while Hha(N)::RspA(C) showed high self-complementation, higher than Hha-splitFAST (**Fig. 4a-e**). These results suggest that the sequence of the N-terminal fragment controls the extent of the self-complementation. Very similar conclusions could be drawn from the analysis of the dissociation experiments (**Fig. 4f-h**).

**Figure 4.**
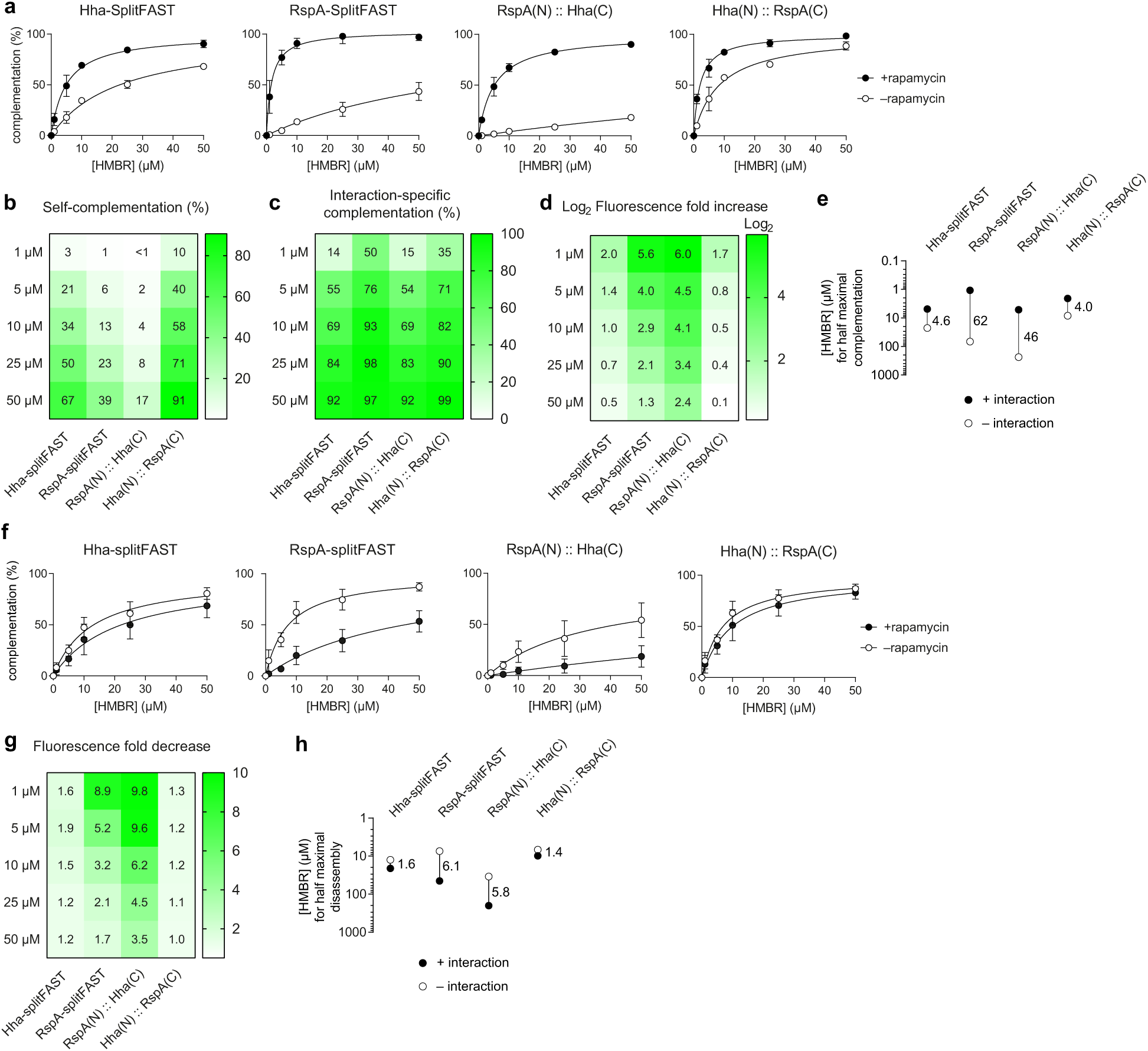
Flow cytometry analysis of the chimera resulting from the fragment crossing of Hha-splitFAST and Rspa-splitFAST. **a** Normalized average fluorescence of about 50,000 HEK293T cells co-expressing the FK506-binding protein (FKBP) fused to the C-terminal fragment of the split reporters and the FKBP-rapamycin-binding domain of mammalian target of rapamycin (FRB) fused to the N-terminal fragment of the split reporters treated without or with 500 nM of rapamycin, and with 1, 5, 10, 25 or 50 μM of HMBR (see **Fig. S2**). Data represent the mean ± standard deviation of three independent experiments. These data allowed to determine for each split reporter **b** the percentage of self-complementation, **c** the percentage of interaction-specific complementation and **d** the fluorescence fold increase (dynamic range) (at various fluorogen concentrations. **e** Concentration of fluorogen for half maximal complementation in presence (EC_50,+interaction_) or absence (EC_50,–interaction_) of rapamycin (see also **Table S6**). The ratio EC_50,– interaction_ / EC_50,+interaction_ is indicated for each ortholog-based split reporter. **f** Normalized average fluorescence of about 50,000 HEK293T cells co-expressing the homodimerizing FK506-binding protein F36M mutant (FKBP_F36M_) fused to the N-terminal and C-terminal fragments of the split reporters treated without or with 500 nM of rapamycin, and with 1, 5, 10, 25 or 50 μM of HMBR. Data represent the mean ± standard deviation of three independent experiments. These data allowed to determine for each split reporter **g** the fluorescence fold decrease at various fluorogen concentrations. **h** Concentration of fluorogen for half maximal complementation in presence (EC_50,–interaction_) or absence (EC_50,+interaction_) of rapamycin (see also **Table S7**). The ratio EC_50,+interaction_ / EC_50,–interaction_ is indicated for each ortholog-based split reporter.

The results previously shown on **Fig. 2f** suggested that the improved dynamic range of RspA-splitFAST resulted not only from a lower self-complementation but also from a more efficient interaction-dependent complementation. This later parameter is directly linked to the binding affinity of the fluorogen for the complemented protein. Interestingly, we observed that replacing the C-terminal fragment of Hha-splitFAST by that of RspA-splitFAST increased the efficiency of interaction-dependent complementation, while replacing the C-terminal fragment of RspA-splitFAST by that of Hha-splitFAST decreased it (**Fig. 4a-e**).

Next, we evaluated the complementation kinetics of RspA-splitFAST, Hha-splitFAST and their crossed chimera by time-lapse confocal microscopy (**Fig. 5a-c** and **Movies S1-S4**). To do so, we first followed the rapamycin-induced heterodimerization of FRB and FKBP in presence of 5 μM HMBR. We observed that the four systems display comparable complementation kinetics (**Fig. 5a,b** and **Movies S1-S4**). Analysis of the fluorescence fold increase upon rapamycin addition in single cells allowed us to characterize the dynamic range of the systems in fluorescence microscopy. RspA-splitFAST and RspA(N)::Hha(C) displayed high dynamic ranges, while Hha-splitFAST and Hha(N)::RspA(C) displayed lower dynamic ranges (**Fig. 5c**), in agreement with the results obtained by flow cytometry.

**Figure 5.**
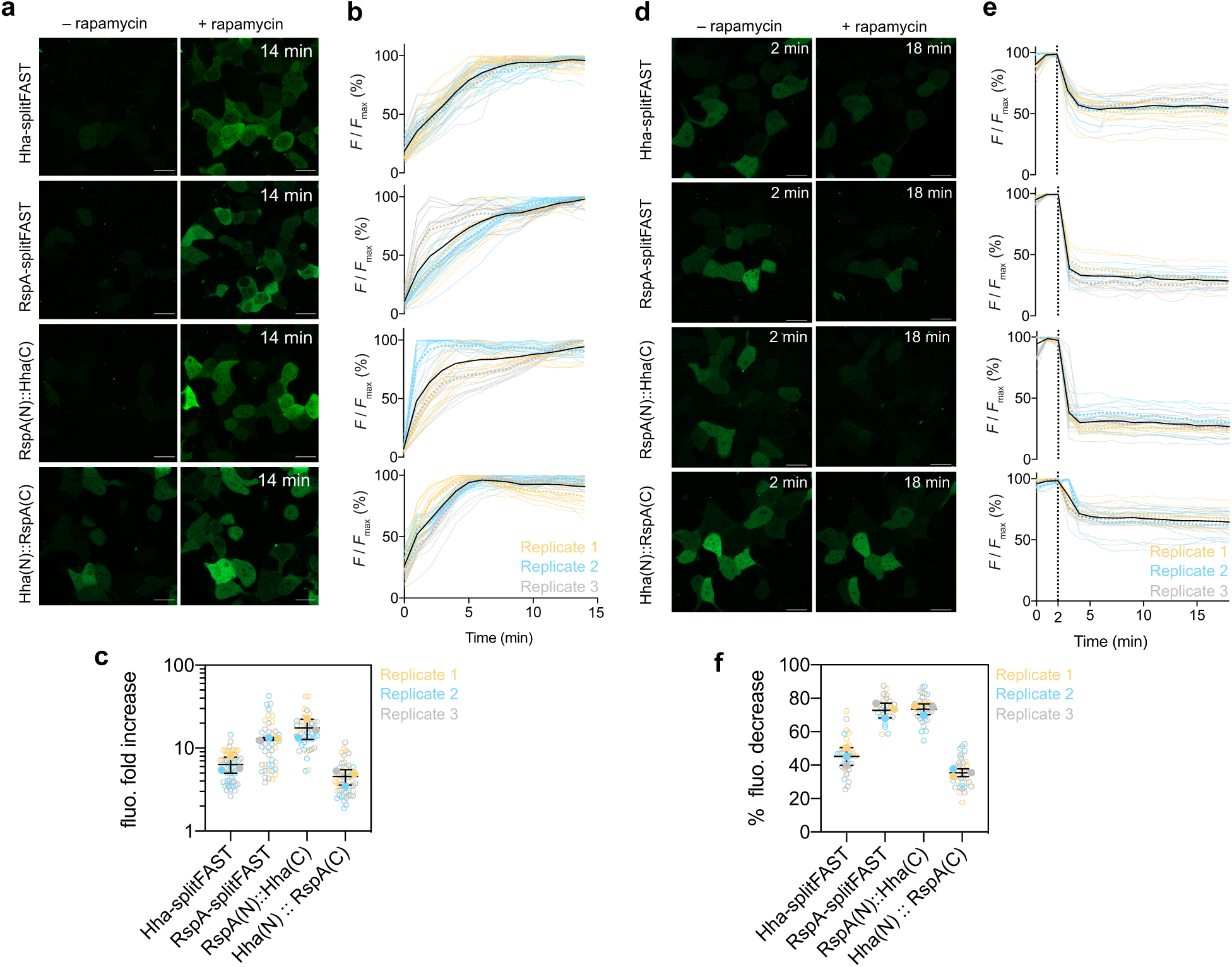
Kinetics of complementation and disassembly. **a-c** HEK293T cells co-expressing the FK506- binding protein (FKBP) fused to the C-terminal fragment of the split reporters and the FKBP-rapamycin-binding domain of mammalian target of rapamycin (FRB) fused to the N-terminal fragment of the split reporters were treated with 5 μM of HMBR. Cells were imaged by time-lapse confocal microscopy after addition of 100 nM of rapamycin. Experiments were repeated three times with similar results. **a** Representative micrographs before and after addition of rapamycin (see also **Movies S1-S4**). **b** Temporal evolution of the fluorescence signal after addition of rapamycin n = 57 (Hha), 47 (RspA), 44 (RspA(N)::Hha(C)), 53 (Hha(N)::RspA (C)) cells from three independent experiments. Each cell is color-coded according to the biological replicate it came from. The dotted lines represent the mean value of each biological replicate, while the black line represents the mean of the three biological replicates. **c** Fluorescence fold increase upon addition of rapamycin. Each cell is color-coded according to the biological replicate it came from. The solid circles correspond to the mean of each biological replicate. The black line represents the mean ± SD of the three biological replicates. **d-f** HEK293T cells co-expressing the homodimerizing FK506-binding protein F36M mutant (FKBP_F36M_) fused to the N-terminal and C-terminal fragments of the split reporters were treated with 5 μM of HMBR. Cells were imaged by time-lapse confocal microscopy after addition of 2.5 μM of rapamycin. Experiments were repeated three times with similar results. **d** Representative micrographs before and after addition of rapamycin. **e** Temporal evolution of the fluorescence signal after addition of rapamycin of n = 33 (Hha), 25 (RspA), 30 (Hha(N)::RspA (C)), 35 cells (RspA(N)::Hha(C)) from three independent experiments. Each cell is color-coded according to the biological replicate it came from. The dotted lines represent the mean value of each biological replicate, while the black line represents the mean of the three biological replicates. **f** Percentage of fluorescence loss upon addition of rapamycin. Each cell is color-coded according to the biological replicate it came from. The solid circles correspond to the mean of each biological replicate. The black line represents the mean ± SD of the three biological replicates. Scale bars 10 μm.

Next, we evaluated the kinetics and amplitude of dissociation of the different systems by monitoring the rapamycin-induced dissociation of the FKBP-F36M homodimer through time-lapse confocal microscopy (**Fig. 5d-f**). We observed very similar kinetics of dissociation for the four systems. In agreement with the self-complementation properties of each systems, we observed improved dynamic range for the dissociation of RspA-splitFAST, and RspA(N)::Hha(C), and reduced dynamic range for the dissociation of Hha(N)::RspA(C).

Overall, this set of experiments further demonstrated that RspA-splitFAST outperformed Hha-splitFAST to image dynamic and reversible PPIs.

### Thermodynamic characterization of RspA-splitFAST and Hha-splitFAST

To get further insights on the parameters responsible for the improvement of RspA-splitFAST, we determined the thermodynamic parameters characterizing the self-complementation of the ternary assembly for RspA-splitFAST and Hha-splitFAST. The complemented systems are composed of three parts, the fluorogen (F) and the N-and C-terminal fragments (hereafter called N and C). Assuming that F and C have negligible affinity, the ternary assembly NCF can result only from two paths (**Fig. S8a**). First, C can bind to N to form NC, and then F binds NC to form NCF. The two steps are characterized respectively by the thermodynamic binding constant *K*_d,C_ and *K*_d,F/C_. A second path involves first F binding to N to form a complex NF, and then in the second step C binding to NF to form the ternary assembly NCF. These two steps are characterized respectively by the thermodynamic binding constant *K*_d,F_ and *K*_d,C/F_. As the binding of C to N increases the binding affinity of F, *K*_d,F/C_ and *K*_d,F_ are linked by a cooperativity constant α, such that *K*_d,F/C_ = *K*_d,F_ / α (with α > 1). As the two paths form a thermodynamic box, reciprocally, the binding of F to N increases the affinity of C in the same extent, such that *K*_d,C/F_ = *K*_d,C_ / α. The full theoretical model is described in **supporting text 1**.

We used fluorescence titration experiments to determine the different thermodynamic constants. First, we characterized the *K*_d,F/C_ constant, corresponding to the affinity constant of the fluorogen for the assembled NC complex. To do so, we recombinantly expressed FRB-Hha/RspA(N) and FKBP-Hha/RspA(C) fusions in *E. coli* and purified the proteins by affinity chromatography. Then, we preassembled the NC complex through addition of rapamycin, and determined the fraction of fluorescent ternary complex at different concentrations of the fluorogen HMBR. This fluorescence titration experiment allowed us to extract *K*_d,F/C_ = 150 ± 10 nM and 1.1 ± 0.1 μM for RspA-splitFAST and Hha-splitFAST, respectively (**Fig. S8b,d**). The higher affinity of HMBR for RspA-splitFAST versus Hha-splitFAST agreed with the trend observed with the full-length proteins (see **Table S1**), and with our experiments in cells showing that the interaction-dependent complementation was more efficient with RspA-splitFAST than with Hha-splitFAST.

To determine *K*_d,F_ and *K*_d,C_, we then performed fluorescence titration experiments using recombinant N-terminal fragments and synthetic C-terminal fragments. Fluorescence titrations were performed at a constant concentration of the N-terminal fragment varying both the concentration of the C-terminal fragment and the concentration of the fluorogen. A global fit of the data using the expression of the fraction of the ternary complex NCF in function of [C] and [F] (see supporting text for full explanation), knowing the *K*_d,F/C_ values, allowed us to estimate *K*_d,C_ = 3.1 ± 0.3 μM and *K*_d,F_ = 2.0 ± 0.2 μM in the case of Hha-splitFAST(**Fig. S8c,d**). In the case of RspA-splitFAST, the global fit gave *K*_d,C_ = 220 ± 10 μM and *K*_d,F_ = 21 ± 2 μM (**Fig. S8c,d**). Because of the multiparametric nature of the global fit, the extracted thermodynamic constants are estimates; however, these constants suggest that the lower self-association of RspA-splitFAST is due to a reduction of the affinity of its N-terminal fragment with both the C-terminal fragment and the fluorogen. The fact that once complemented RspA-splitFAST displays a higher fluorogen binding affinity than Hha-splitFAST is due to a large increase in the cooperativity constant (**Fig. S8d**).

### Applications of RspA-splitFAST for the study of protein-protein interactions

To demonstrate the ability of RspA-splitFAST to monitor dynamic PPI in live cells, RspA-splitFAST was first used to visualize in real-time the recruitment of a cytosolic protein at the plasma membrane through an inducible PPI (**Fig. 6a,b**). We expressed in HEK293T cells FKBP-RspA(C) in the cytosol and targeted FRB-RspA(N) to the plasma membrane using the targeting sequence of Lyn protein kinase (Lyn11-FRB-RspA(N)). After treatment with HMBR, the recruitment of FKBP-RspA(C) to the plasma membrane was induced by addition of rapamycin. RspA-splitFAST rapidly and efficiently forms a fluorescent complex at the plasma membrane as measured using time-lapse confocal microscopy. Thus, RspA-splitFAST can monitor PPI at the plasma membrane in real-time.

**Figure 6.**
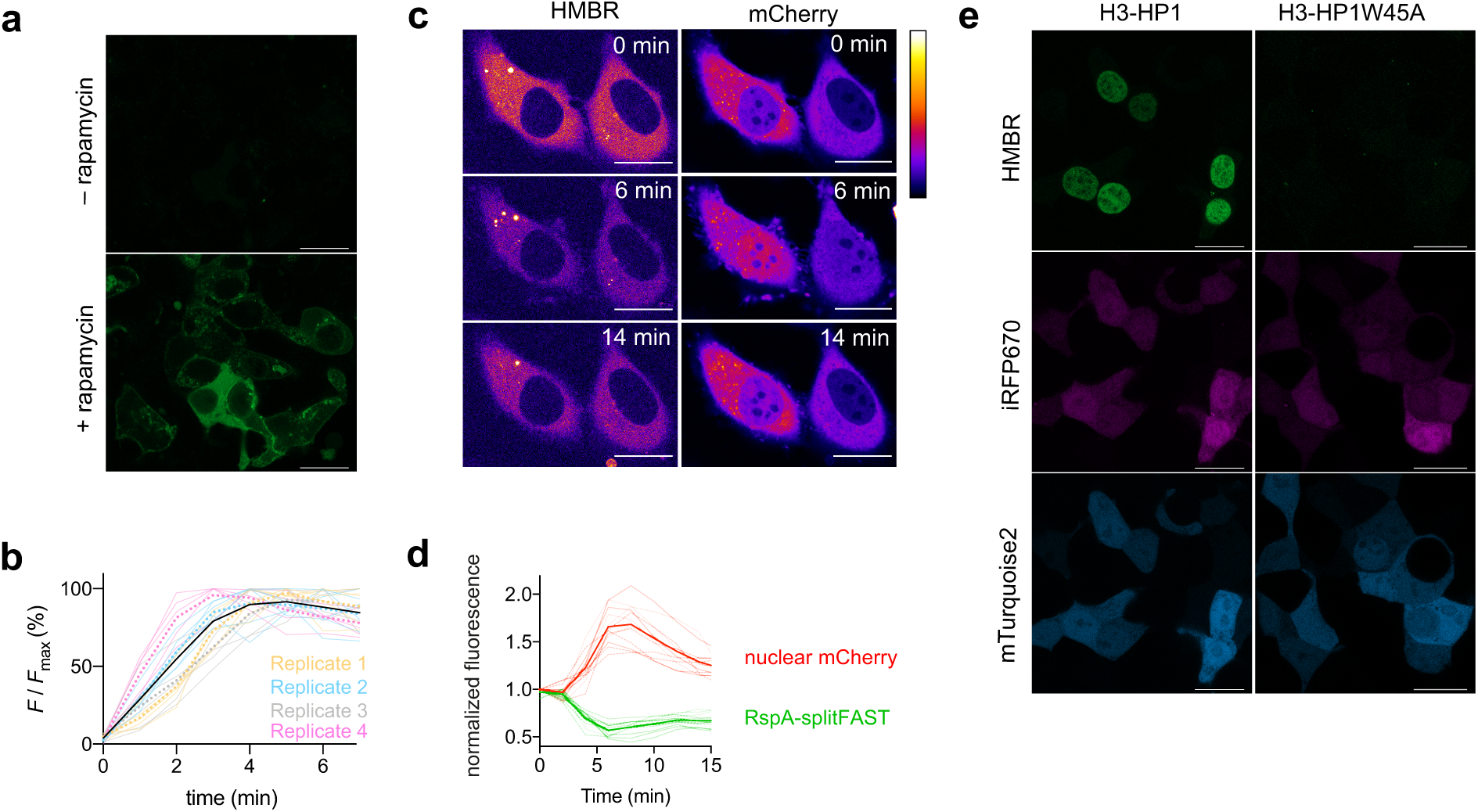
RspA-splitFAST enables the detection of PPI in various contexts. **a,b** HEK293T cells co-expressing Lyn11-FRB-RspA(N) and FKBP-RspA(C) were labeled with 5 μM HMBR and imaged before and after the addition of 100 nM rapamycin. **a** Representative images before and after rapamycin addition. **b** Temporal evolution of the fluorescence intensity after rapamycin addition in HMBR-treated cells co-expressing Lyn11-FRB-RspA(N) and FKBP-RspA(C) of n = 21 cells from 4 independent experiments. Cells from the same biological replicate are displayed with the same color. The dot lines represent the mean value of each biological replicate, while the black line represents the mean of the 4 biological replicates. **c,d** Use of splitFAST for imaging the evolution of MEK1/ERK2 interaction upon epidermal growth factor (EGF) stimulation. **c** Representative images of HMBR-labeled HeLa cells co-expressing MEK1-RspA(N) and mCherry-ERK2-RspA(C) after stimulation with EGF (see also **Movie S5**). **d** Temporal evolution of RspA-splitFAST fluorescence (green) and nuclear mCherry fluorescence (red) intensities of n = 12 cells from 3 independent experiments. Cells from the same biological replicate are displayed with the same line pattern. The thick solid lines represent the mean value of the 3 biological replicates. **e** Monitoring of the ^methylated^H3 – HP1 interaction. RspA(N)-H3 (H3: Histone 3) and HP1-RspA(C) (HP1: chromodomain of the heterochromatin protein 1) were co-expressed in HEK293T cells using two bicistronic plasmids allowing the expression of mTurquoise2 and IRFP2. Selectivity was validated using HP1W45A. Cells were treated with 5 μM of HMBR. Experiments were repeated three times with similar results. **a,c,e** Scale bars 20 μm.

To further demonstrate the suitability of RspA-splitFAST for the monitoring of dynamic PPIs, we imaged the interaction between the MAP kinase kinase, MEK1, and the downstream extracellular signal-regulated protein kinase, ERK2, one of the central interactions in the Raf/MEK/ERK signaling pathway (**Fig. 6c,d** and **Movie S5**). Upon stimulation with *e.g.,* epidermal growth factor, MEK1 phosphorylates ERK2, inducing its dissociation and subsequent translocation to the nucleus [18, 19], where it regulates the activity of transcription factors. Dephosphorylation of ERK2 by nuclear phosphatases cause its return to the cytoplasm [20-22]. MEK1 was fused to the N-terminal fragment of RspA-splitFAST and ERK2 to the C-terminal fragment. An additional mCherry fluorescent protein was fused to the ERK2 fusion to visualize its change of localization upon activation of the MAPK signaling pathway. The two proteins were co-expressed in HeLa cells, and cells were treated with HMBR. Prior to stimulation, we observed specific, cytosolic RspA-splitFAST signal, consistent with the role of MEK1 in sequestering ERK2 in the cytosol. Using confocal microscopy, we observed the partial loss of RspA-splitFAST fluorescence after application of epidermal growth factor. This was correlated to an accumulation of nuclear mCherry fluorescence, which is consistent with the dissociation of the MEK1- ERK2 cytosolic complex and translocation of ERK2 to the nucleus. After a few minutes, we observed nuclear-to-cytosolic shuttling of mCherry-ERK2-RspA(C) and an increase in the cytosolic RspA-splitFAST signal, in agreement with desensitized mCherry-ERK2-RspA(C) returning to the cytosol and reassembling with MEK1-RspA(N). This experiment allowed us to demonstrate the ability of RspA-splitFAST to monitor dynamic PPI in signaling pathways in real-time.

To further expand the use of RspA-splitFAST, we next studied its use for the detection of H3K9 methylation-dependent interactions (**Fig. 6e**). Histone H3 lysine 9 (H3K9) methylation is a conserved mark of transcriptional silencing that is involved in the formation of heterochromatin [23]. Methylated H3K9 (H3K9m) is specifically recognized by the Heterochromatin protein 1 (HP1). HP1 is a transcription repressor composed of two conserved domains: the chromodomain at the N-terminus, which binds H3K9m, and the chromo shadow domain at the C-terminus, which is implicated in PPI [24, 25]. HP1 recognized the cationic and hydrophobic methyl ammonium of H3K9m via cation-π interactions with its aromatic residues Y24, W45 and Y48 and by van der Waals interactions with its residues E23 and E52. In order to detect the H3K9m-HP1 interaction, we fused the N-terminal fragment of RspA-splitFAST to the N-terminus of H3 (RspA(N)- H3) and its C-terminal fragment to the C-terminus of HP1 (HP1-RspA(C)). The two proteins RspA(N)-H3 and HP1-RspA(C) were co-expressed in HEK293T cells using two bicistronic vectors that allowed also the expression of mTurquoise2 and iRF670, respectively, as transfection reporters. To evaluate the specificity of the complementation, we constructed a negative control by inserting the mutation W45A in HP1, previously shown to prevent interaction with methylated H3K9 [26]. Evaluation of the fluorescence complementation efficiency of RspA-splitFAST by flow cytometry and fluorescence microscopy allowed us to show the specific interaction of RspA(N)-H3 and HP1-RspA(C). The mutation W45A in HP1 led to a loss of fluorescence complementation, in agreement with the key role of W45 in the recognition of H3K9m. This experiment allowed us to show the ability of RspA-splitFAST to detect the effect of a single mutation on the efficiency of a PPI.

## DISCUSSION

Detection of PPI with high spatiotemporal resolution is facilitated by split complementation systems that can capture the often-transient nature of these interactions with high contrast. We previously developed a promising system, splitFAST (renamed Hha-splitFAST in this study), which displays rapid and reversible complementation. Hha-splitFAST functions as a ternary complex between two complementary protein fragments and a small fluorogenic chromophore, which renders engineering of this system challenging. It is based on the chemogenetic reporter, FAST, which was engineered from *Halorhodospira halophila* (Hha)-PYP. PYPs are part of a large protein family and many orthologs have been reported, although relatively few have been characterized. Here we explored the natural sequence diversity of PYP orthologs to overcome some of the limitations in combinatorial screening in order to quickly access improved split systems.

We selected six orthologs of PYP that display between 70 and 78% sequence homology with Hha-PYP. We replaced the sequence of their loop 94-101 with that of canonical FAST (identified through directed evolution), and evaluated their fluorogen binding ability *in vitro.* Not only did all generated proteins bind the three FAST fluorogens tested, but many formed tighter and/or brighter complexes than canonical FAST. All systems were shown to be fully functional in live cell fluorescence imaging. Because of their conserved functions and of the orthology relationship of their parent PYP proteins, the six generated proteins were called FAST orthologs.

Next, we used these FAST orthologs to generate split systems using the same split site used originally to generate splitFAST. To evaluate the complementation properties of the ortholog-based split systems, we used a flow cytometry assay to be able to simultaneously trigger a PPI as well as test the effect of fluorogen concentration on the self-assembly of the split construct. We found that all six orthologs tested were able to function as complementation reporters. Several orthologs showed greater dynamic range and higher brightness than the original Hha-splitFAST. In particular, the RspA-splitFAST system originating from *Rheinheimera sp.* A13L (RspA) PYP was brighter and displayed lower self-association but higher efficacy of interaction-dependent complementation, resulting overall in a high increase in dynamic range. We had previously observed that the affinity of the two protein fragments could be modulated by truncation of the small fragment, resulting in a system, named Hha-splitFASTΔC1, that showed higher dynamic range, although much lower overall brightness. RspA-splitFAST showed higher dynamic range and higher efficacy of interaction-dependent complementation than both Hha-splitFAST and Hha-splitFASTΔC1 in all of our assays.

Given the diversity of behavior observed among the seven orthologous split systems, we systematically crossed their N-terminal and C-terminal fragments to explore a wider sequence space and evaluate their complementation properties. Analysis of the chimeric split systems allowed us to confirm that RspA-splitFAST was the most improved system, and allowed an analysis of the role of the N-and C-terminal fragments on the efficacy of the self-and interaction-dependent complementation, as well as on the fluorogen binding affinity and brightness. In particular, our analysis suggested that the reduction of the self-complementation of RspA-splitFAST was mainly due to the sequence and properties of its N-terminal fragment, which was furthermore shown to display a lower affinity both for the C-terminal fragment and for the fluorogen in vitro.

The improved brightness, lower self-association and higher dynamic range of RspA-splitFAST allowed us to study dynamic interactions in live cells. Its lower self-complementation minimizes background signal from non-interacting proteins, and permits more efficient dissociation upon disassembly of a given protein complex.

In conclusion, the use of an original orthology-based protein engineering allowed us to optimize the properties of splitFAST, and generate RspA-splitFAST, a split fluorescent reporter of PPI displaying improved brightness and dynamic range. Because of the rapidity and reversibility of its complementation, and its low self-complementation, high dynamic range and improved brightness, RspA-splitFAST is well suited for monitoring dynamic interactions with high spatiotemporal resolution and opens exciting prospects to decipher the role of PPI in complex interaction networks, to create new cellular biosensors and to develop innovative screening assays for the discovery of PPI modulators.

## ACKNOWLEDGMENTS

We thank the imaging facility of the Institut de Biologie Paris Seine of Sorbonne University. This work has been supported by the European Research Council (ERC-2016- CoG-724705 FLUOSWITCH), and the Institut Universitaire de France.

## AUTHOR CONTRIBUTIONS

L.M.R., A.G.T. and A.G. designed the experiments. L.M.R., A.G.T. and D. B. performed the experiments. L.M.R., A.G.T. and A.G. analyzed the experiments. L.M.R., A.G.T. and A.G. wrote the paper.

## MATERIALS AND METHODS

### General

Synthetic oligonucleotides used for cloning were purchased from Sigma-Aldrich or Integrated DNA Technology. PCR reactions were performed with Q5 polymerase (New England Biolabs) in the buffer provided. PCR products were purified using QIAquick PCR purification kit (Qiagen). Restriction endonucleases, T4 ligase, Taq ligase, and Taq exonuclease were purchased from New England Biolabs and used with accompanying buffers and according to the manufacturer’s protocols. Isothermal assemblies (Gibson assembly) were performed using homemade mix prepared according to previously described protocols [27]. Restriction enzyme cloning were performed with BglII and BspEI restriction enzymes (New England Biolabs) in the buffer provided. The products of restriction enzyme digests were purified by preparative gel electrophoresis followed by QIAquick Gel Extraction Kit (Qiagen). Ligation were performed with T4 DNA ligase (New England Biolabs) in the buffer provided. Small-scale isolation of plasmid DNA was done using QIAprep miniprep kit (Qiagen) from 2mL of overnight culture. Large-scale isolation of plasmid DNA was done using the QIAprep maxiprep kit (Qiagen) from 150 mL of overnight culture. All plasmid sequences were confirmed by Sanger sequencing with appropriate sequencing primers (GATC-Biotech). Rapamycin was purchased from Sigma-Aldrich and dissolved in dimethyl sulfoxide to a concentration of 3 mM. Human recombinant EGF was purchased from Sigma-Aldrich and dissolved in sterilized water to a concentration of 500 μg/mL to be conserved. Peptides corresponding to the C-terminal of RspA-splitFAST and Hha-splitFAST were purchased from Clinisciences at 98% purity and are acetylated and amidated at the N and C termini. The preparation of HMBR, HBR- 3,5DM and HBR-3,5DOM was previously described[4, 13], and are commercially available from The Twinkle Factory under the name ^TF^Lime, ^TF^Amber and ^TF^Coral. The construction of the plasmids used in this study is described in **Supplementary Information**. A list of primers is given in **Table S8**, a list of plasmids is given in **Table S9** and sequences are provided in **Table S10**.

### Mammalian cell culture

HEK293T cells (ATCC CRL-3216) were cultured in Dulbecco’s modified Eagle’s medium (DMEM) supplemented with phenol red, Glutamax I, and 10% (vol/vol) fetal calf serum, at 37 °C in a 5% CO2 atmosphere. HeLa cells (ATCC CRM-CCL2) were cultured in Minimal Essential Media (MEM) supplemented with phenol red, Glutamax I, 1 mM of sodium pyruvate, 1% (vol/vol) of non-essential amino-acids and 10% (vol/vol) fetal calf serum (FCS), at 37 °C in a 5% CO2 atmosphere. For imaging, cells were seeded in μDish IBIDI (Biovalley) coated with poly-L-lysine. Cells were transiently transfected using Genejuice (Merck) according to the manufacturer’s protocol for 24 h prior to imaging. Live cells were washed with DPBS (Dulbecco’s Phosphate-Buffered Saline), and treated with DMEM media (without serum and phenol red) supplemented with the compounds at the indicated concentration. For the MEK1-ERK2 experiment, cells were serum-starved 15 hours before activation by EGF.

### Flow cytometry

Flow cytometry experiments were performed on MACSQuant® Analyser equipped with three LASER (405 nm, 488 nm and 635 nm) and eight filters and channels. To prepare samples, HEK 293 cells were first grown in cell culture flasks, then transiently co-transfected 24 h after seeding using Genejuice (Merck) according to the manufacturer’s protocol. After 24 h, transfected cells were trypsinized and centrifuged in PBS-BSA (1 mg/mL) and resuspend in PBS-BSA with the appropriate amounts of fluorogens (final volume of the cell suspension 200 μL). Simple positive controls with cells expressing only mTurquoise2, iRFP670 and FAST were used to adjust PMT voltage and compensation matrix. For each experiment, 50,000 cells positively expressing mTurquoise2 (Ex 434 / Em 450 ± 25) and iRFP670 (Ex 638 nm / Em 660 ± 10 nm) were analyzed with the following parameters: Ex 488nm, Em 525 ± 20 nm for cells labeled with HMBR and HBR-3,5DM; Ex 488nm, Em 585 ± 20 nm for cells labeled with HBR-3,5DOM. Data were analyzed using FlowJo v10.7.1. The analysis of the rapamycin-induced FRB-FKBP association was performed as followed: the evolution of the mean cell fluorescence in function of the concentration of fluorogen in presence and in absence of rapamycin was fitted with a one site binding model equation with GraphPad Prism v9.3.0, sharing the maximal value for the experiments in presence and in absence of rapamycin. This allows to determine the concentration of fluorogen for half maximal complementation, as well as the theoretical maximum fluorescence value. This latter value is then used to compute the percentage of complementation for each concentration of fluorogen. Similar analysis was performed to analyze the rapamycin-induced dissociation of FKBP-F36M homodimer. To determine the relative brightness of the different systems at 50 μM of fluorogen, we analyzed the fluorescence of cells with similar expression levels, using the transfection reporters, mTurquoise2 and iRFP670 to select cells with approximately equivalent expression levels.

### Fluorescence microscopy

Micrographs were acquired on a Zeiss LSM 980 Laser Scanning Microscope equipped with a plan apochromat 63x /1.4 NA oil immersion objective. ZEN software was used to collect the data. The images were analyzed using Icy (2.4.0.0) and Fiji (Image J). To track fluorescence signal in the nucleus and the cytoplasm, their contour was determined by masking the signal in the mTurquoise2 channel with the plug-in HK-means with the following parameters: intensity class equals 100, min object size (px) equals 5000, max object size (px) equals 15000-2000. The signal intensity of the ROI was tracked over the time for each channel by using plug-in Active Contours. The background signal was subtracted. Data were processed using GraphPad Prism v9.3.0.

### Protein production and purification

Plasmids were transformed in Rosetta (DE3) pLys *E. coli* (New England Biolabs). Cells were grown at 37°C in lysogen broth (LB) medium supplemented with 50 μg/ml kanamycin and 34 μg/mL chloramphenicol to OD600nm 0.6. Expression was induced at 37°C during 4 h by adding isopropyl-β-D-1- thiogalactopyranoside (IPTG) to a final concentration of 1 mM. Cells were harvested by centrifugation (4,300 × g for 20min at 4°C) and frozen. The cell pellet was resuspended in lysis buffer (PBS supplemented with 2.5 mM MgCl_2_, 1 mM of protease inhibitor PhenylMethaneSulfonyl Fluoride PMSF, 0.025 mg/mL of DNAse, pH 7.4) and sonicated (5 min, 20 % of amplitude) on ice. The lysate was incubated for 2-3 hours on ice to allow DNA digestion by DNAse. Cellular fragments were removed by centrifugation (9,000 × g for 1 h at 4°C). The supernatant was incubated overnight at 4°C by gentle agitation with pre-washed Ni-NTA agarose beads in PBS buffer complemented with 20 mM of imidazole. Beads were washed with 10 volumes of PBS complemented with 20 mM of imidazole and with 5 volumes of PBS complemented with 40mM of imidazole. His-tagged proteins were eluted with 5 volumes of PBS complemented with 0.5 M of imidazole. The buffer was exchanged to PBS (0.05 M phosphate buffer, 0.150 M NaCl) using PD-10 desalting columns (GE Healthcare).

### Determination of the thermodynamic dissociation constants

Thermodynamic dissociation constants were determined by performing titration experiments using a Spark 10M plate reader (Tecan) and fitting data in GraphPad Prism v9.3.0 using the binding models described in **Supplementary Text 1**. To determine *K*_d,F/C_, an equimolar solution of 50 nM of purified recombinant FRB-N and FKBP-C proteins were titrated with various concentrations of HMBR (0, 40, 80, 160, 320, 480, 640, 960, 1280, 2560 and 5000 nM) in presence of 500 nM rapamycin. To determine *K*_d,F_ and *K*_d,C_, 0.1 μM of recombinant N-terminal fragment was titrated with various concentrations of HMBR (0, 1, 2.5, 5, 7.5, 10, 15 μM) at different concentrations of the synthetic C-terminal fragment (0.5, 1, 2, 3, 4, 5, 6, 7, 8 and 9 μM).

