## Supplementary information for "Improving split reporters of protein-protein interactions through orthology-based protein engineering"

**This PDF file includes:**

Legends of Movies S1-S5

Supplementary Text 1

Fig. S1 to S8

Tables S1 – S10

Supplementary references

**Legends of Supplementary Movies**

**Movie S1. Association of Hha-splitFAST.** HEK293T cells co-expressing the FK506-binding protein (FKBP) fused to the C-terminal fragment of Hha-FAST and the FKBP-rapamycin-binding domain of mammalian target of rapamycin (FRB) fused to the N-terminal fragment of Hha-FAST were treated with 5 μM of HMBR. Cells were imaged by time-lapse confocal microscopy after addition of 100 nM of rapamycin. Experiment was repeated three times with similar results (see also **Fig. 5**). Scale bars 10 μm.

**Movie S2. Association of RspA-splitFAST.** HEK293T cells co-expressing the FK506-binding protein (FKBP) fused to the C-terminal fragment of RspA-FAST and the FKBP-rapamycin-binding domain of mammalian target of rapamycin (FRB) fused to the N-terminal fragment of RspA-FAST were treated with 5 μM of HMBR. Cells were imaged by time-lapse confocal microscopy after addition of 100 nM of rapamycin. Experiment was repeated three times with similar results (see also **Fig. 5**). Scale bars 10 μm.

**Movie S3. Association of the RspA(N):Hha(C) chimera.** HEK293T cells co-expressing the FK506-binding protein (FKBP) fused to the C-terminal fragment of Hha-FAST and the FKBP-rapamycin-binding domain of mammalian target of rapamycin (FRB) fused to the N-terminal fragment of RspA-FAST were treated with 5 μM of HMBR. Cells were imaged by time-lapse confocal microscopy after addition of 100 nM of rapamycin. Experiment was repeated three times with similar results (see also **Fig. 5**). Scale bars 10 μm.

**Movie S4. Association of the Hha(N):RspA(C) chimera**. HEK293T cells co-expressing the FK506-binding protein (FKBP) fused to the C-terminal fragment of Hha-FAST and the FKBP-rapamycin-binding domain of mammalian target of rapamycin (FRB) fused to the N-terminal fragment of RspA-FAST were treated with 5 μM of HMBR. Cells were imaged by time-lapse confocal microscopy after addition of 100 nM of rapamycin. Experiment was repeated three times with similar results (see also **Fig. 5**). Scale bars 10 μm.

**Movie S5. Imaging of MEK1/ERK2 interaction upon epidermal growth factor (EGF) stimulation.** HMBR-labeled HeLa cells co-expressing MEK1-RspA(N) and mCherry-ERK2-RspA(C) were stimulated with EGF and imaged by time-lapse confocal microscopy (See also **Fig. S6c,d**). Scale bars 20 μm.

**Supplementary text 1. Theoretical model for the self-association of splitFAST.**

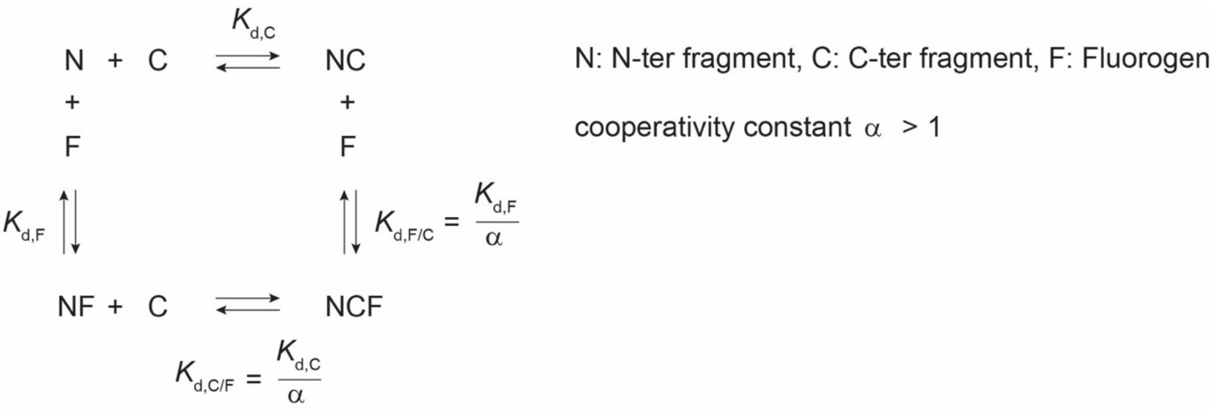


The scheme above shows the ternary equilibrium in which the N-terminal fragment N is able to bind the C-terminal fragment C and the fluorogen F and form a ternary complex NCF. Here it is assumed that C and F cannot interact together to form CF. $K_{d,C}$ and $K_{d,F}$ are the dissociation constant of the complexes NC and NF:

$$K_{d,C}=\frac{[N][C]}{[NC]} \quad (1)$$

$$K_{d,F}=\frac{[N][F]}{[NF]} \quad (2)$$

and $K_{d,F/C}$ and $K_{d,C/F}$ are the dissociation constants for the fluorogen F and the C-terminal fragment C when the N-terminal fragment N is already bound to the C-terminal fragment C and the fluorogen F respectively

$$K_{d,F/C}=\frac{[NC][F]}{[NCF]} \quad (3)$$

$$K_{d,C/F}=\frac{[NF][C]}{[NCF]} \quad (4)$$

One can assume that, when the C-terminal fragment binds the N-terminal fragment, the binding affinity of the fluorogen F raises. One can thus introduce a cooperativity constant $\alpha > 1$ for the binding of the fluorogen F when the C-terminal fragment is bound to the N-terminal fragment:

$$K_{d,F/C}=\frac{K_{d,F}}{\alpha} \quad (6)$$

According to equations (1-4)

$$K_{d,F}\,K_{d,C/F} = K_{d,C}\,K_{d,F/C} \quad (5)$$

Thus, it can be concluded that

$$K_{d,C/F} = \frac{K_{d,C}}{\alpha} \quad (7)$$

Therefore, the influence between the two ligands is reciprocal: if the binding of the C-terminal fragment C raises the binding affinity of the fluorogen F, the binding of the fluorogen F raises the binding affinity of the C-ter fragment C to the same extent. Thus, increasing the concentration of the fluorogen F increases self-association and decreases the dynamic range.

The fraction of the fluorescent ternary complex is given by:

$$F_{\text{ternary complex}} = \frac{[NCF]}{[N] + [NC] + [NF] + [NCF]} \quad (8)$$

$$F_{\text{ternary complex}} = \frac{\alpha \frac{[C][F]}{K_{d,C}K_{d,F}}}{1 + \frac{[C]}{K_{d,C}} + \frac{[F]}{K_{d,F}} + \alpha \frac{[C][F]}{K_{d,C}K_{d,F}}} \quad (9)$$

When $[F]_0$ and $[C]_0 >> [N]_0$, we can assume that at equilibrium $[F] \approx [F]_0$ and $[C] \approx [C]_0$, thus the equation (9) becomes:

$$F_{\text{ternary complex}} = \frac{\alpha \frac{[C]_0[F]_0}{K_{d,C}K_{d,F}}}{1 + \frac{[C]_0}{K_{d,C}} + \frac{[F]_0}{K_{d,F}} + \alpha \frac{[C]_0[F]_0}{K_{d,C}K_{d,F}}} \quad (10)$$

By using the relation between $K_{d,F/C}$, $K_{d,F}$, and α given by the equation (5), the equation of the fraction of the fluorescent ternary complex becomes:

$$F_{\text{ternary complex}} = \frac{\frac{[C]_0[F]_0}{K_{d,C}K_{d,F/C}}}{1 + \frac{[C]_0}{K_{d,C}} + \frac{[F]_0}{K_{d,F}} + \frac{[C]_0[F]_0}{K_{d,C}K_{d,F/C}}} \quad (11)$$

It is possible to determine the different thermodynamic constants by determining the fraction of complex at different concentrations of C and F. A global fit enables to obtain the different parameters.

To facilitate the global fitting, it is possible to independently determine $K_{d,F/C}$ using two interacting proteins fused to N and C to pre-assemble the NC complex. Titration experiments varying the concentration of F allows then to determine $K_{d,F/C}$. Indeed the fraction of fluorescent complex is given by the expression

$$F_{\text{fluorescent complex}} = \frac{[\text{NCF}]}{[\text{NC}] + [\text{NCF}]} \quad (12)$$

$$F_{\text{fluorescent complex}} = \frac{1}{1 + \frac{K_{d,F/C}}{[\text{F}]}} \quad (13)$$

When $[F]_0 >> [N]_0$ and $[C]_0$, we can assume that at equilibrium $[F] \approx [F]_0$, thus the equation (13) becomes:

$$F_{\text{fluorescent complex}} = \frac{1}{1 + \frac{K_{d,F/C}}{[\text{F}]_0}} \quad (14)$$

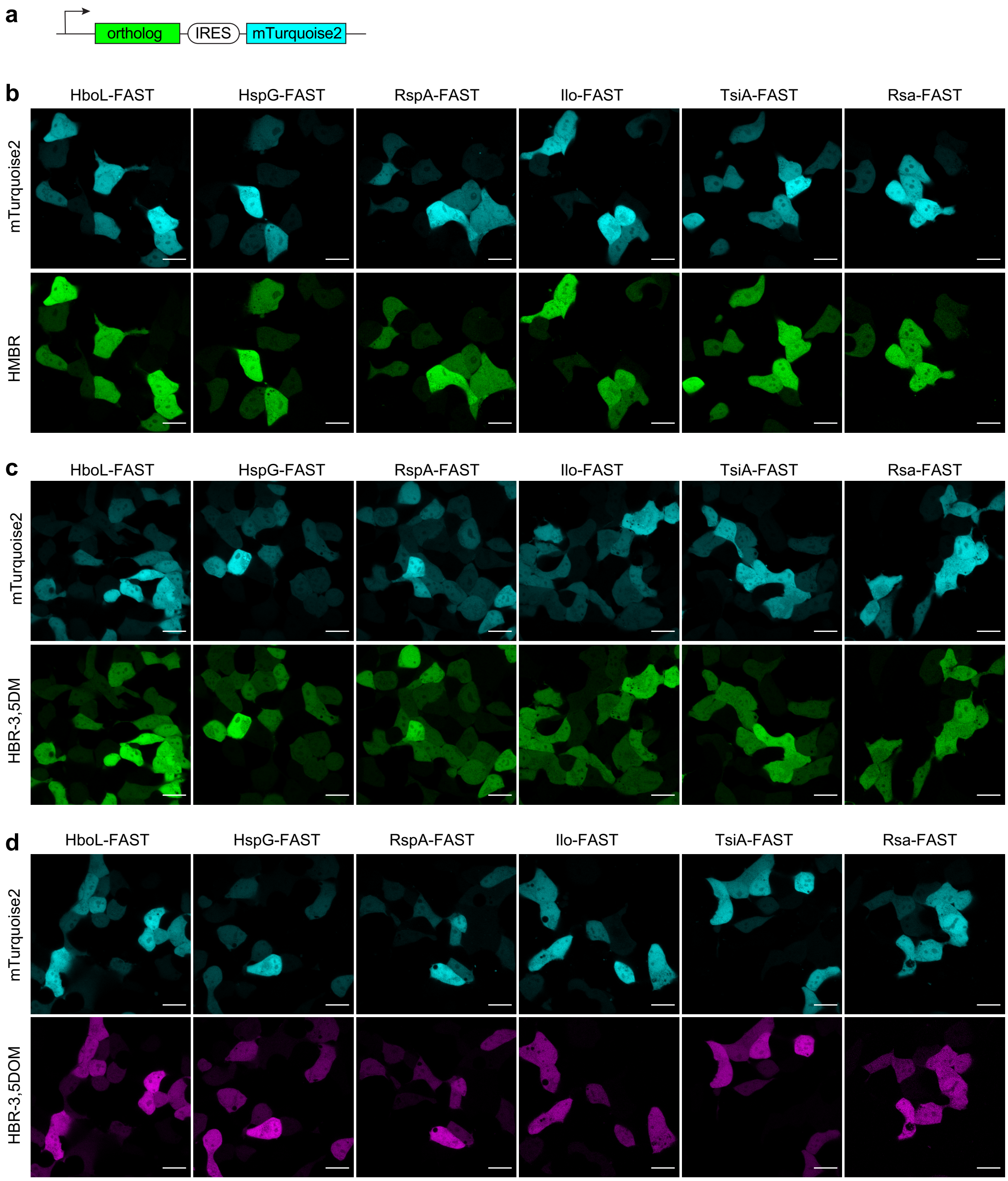


**Figure S1. Evaluation of FAST orthologs in mammalian cells.** HEK293T cells transfected with a bicistronic plasmid allowing the expression of FAST ortholog and mTurquoise2 as transfection reporter (**a**) were treated with 5 μM of HMBR (**b**), HBR-3,5DM (**c**) or HBR-3,5DOM (**d**) and imaged by confocal microscopy. Each sample were imaged with the same imaging settings for direct comparison. Microscopy settings: Cyan: ex/em 405/410-800 nm; Green: ex/em 488/493-800 nm; Magenta: ex/em 543/578-800 nm. All channels were sequentially acquired. Scale bars 10 μm.

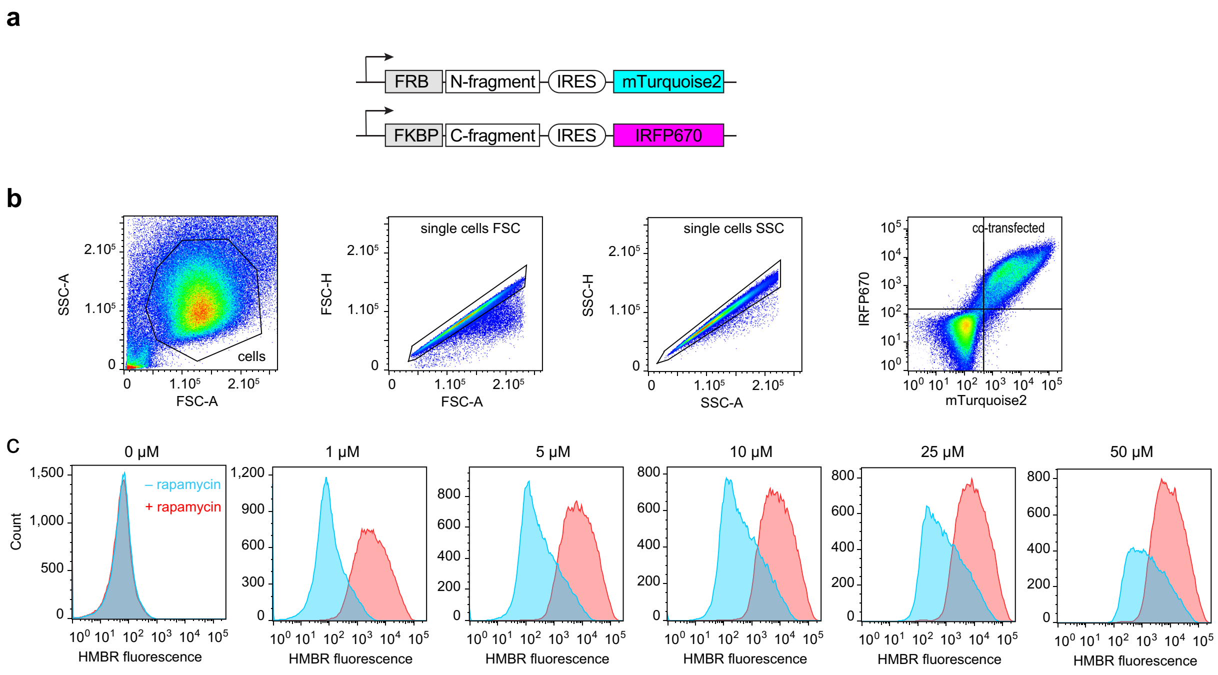


**Figure S2. Flow cytometry workflow. a** Design of the split reporter. The FK506-binding protein (FKBP) fused to the C-terminal fragment of the split reporters and the FKBP-rapamycin-binding domain of mammalian target of rapamycin (FRB) fused to the N-terminal fragment of the split reporters. Biscistronic vectors allowed the co-expression of mTurquoise2 and IRP670 as transfection reporters. **b** Gating strategy. Forward versus side scatter (FSC-A vs SSC-A) density plot was used to identify cells based on size and granularity. Secondly, forward scatter height (FSC-H) vs. forward scatter area (FSC-A) density plot and then side scatter height (SSC-H) vs side scatter area (SSC- A) plot were used to select single cells. Co-transfected cells were then selected by using the signal of IRFP670 and mTurquoise2 transfection reporters. **c** Representative fluorescence analysis. HEK293T cells co-expressing the FK506-binding protein (FKBP) fused to the C-terminal fragment of RspA-split-FAST and the FKBP-rapamycin-binding domain of mammalian target of rapamycin (FRB) fused to the N-terminal fragment of RspA-split-FAST were treated without (blue) or with (red) 500 nM of rapamycin, and with 1, 5, 10, 25 or 50 µM of HMBR and then analyzed by flow cytometry.

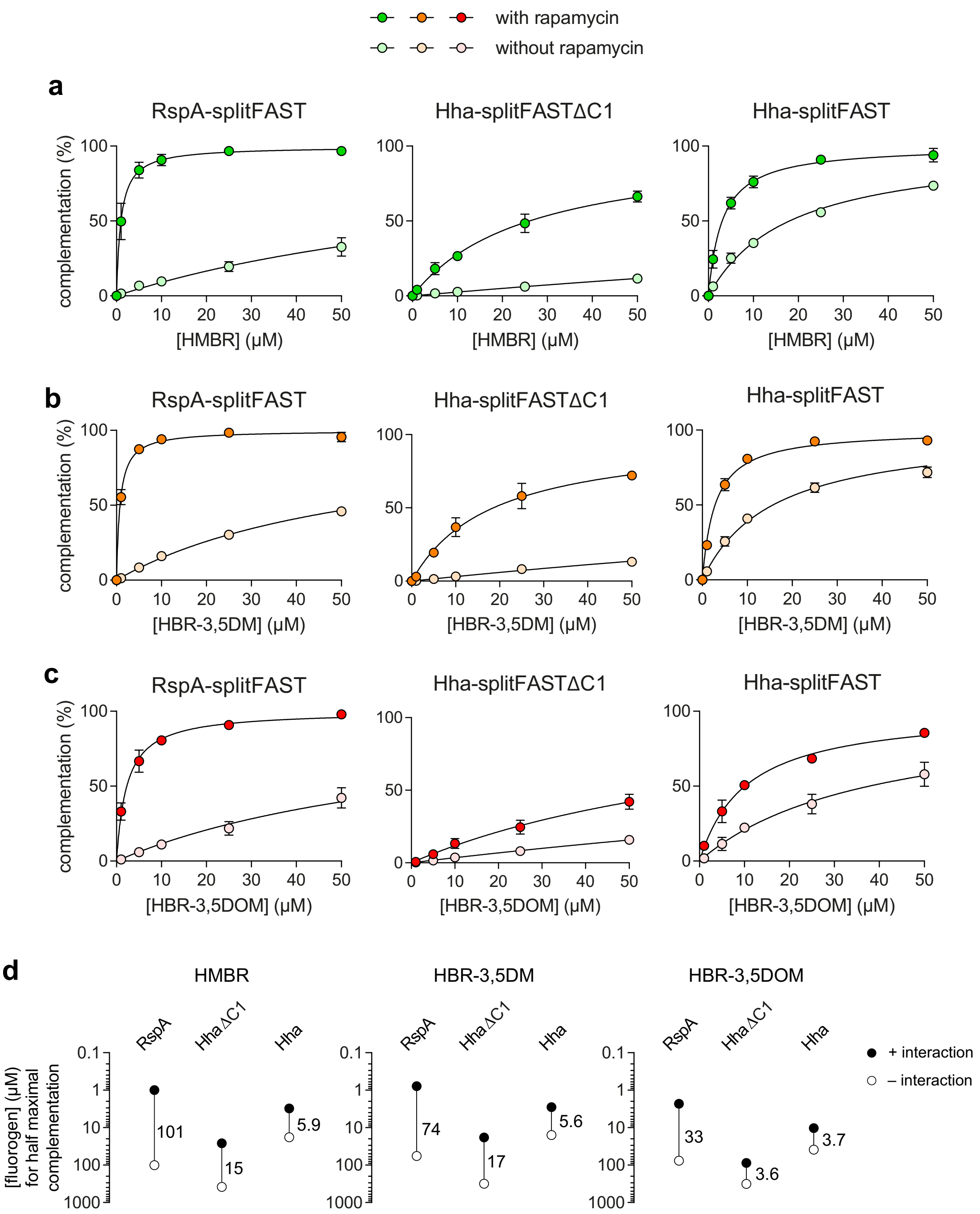


**Figure S3. Association properties of RspA-splitFAST, Hha-splitFAST and Hha-splitFASTΔC1. a-c** Normalized average fluorescence of about 50,000 HEK293T cells co expressing the FK506-binding protein (FKBP) fused to the C-terminal fragment of the split reporters and the FKBP-rapamycin-binding domain of mammalian target of rapamycin (FRB) fused to the N-terminal fragment of the split reporters treated without or with 500 nM of rapamycin, and with 1, 5, 10, 25 or 50 µM of HMBR (**a**), HBR-3,5DM (**b**) or HBR-3,5DOM (**c**) (see **Fig. S2**). Data represent the mean ± standard deviation of three independent experiments. These data allowed to determine for each split reporter the percentage of self-complementation (see **Fig. S4a**), the percentage of interaction-specific complementation (see **Fig. S4b**) and the fluorescence fold increase (dynamic range) (see **Fig. S4c**) at various fluorogen concentrations. **d** Concentration of fluorogen for half maximal complementation in presence ($EC_{50,+interaction}$) or absence ($EC_{50,–interaction}$) of rapamycin (See also **Table S3**). The ratio $EC_{50,–interaction}$ / $EC_{50,+interaction}$ is indicated for each split reporter.

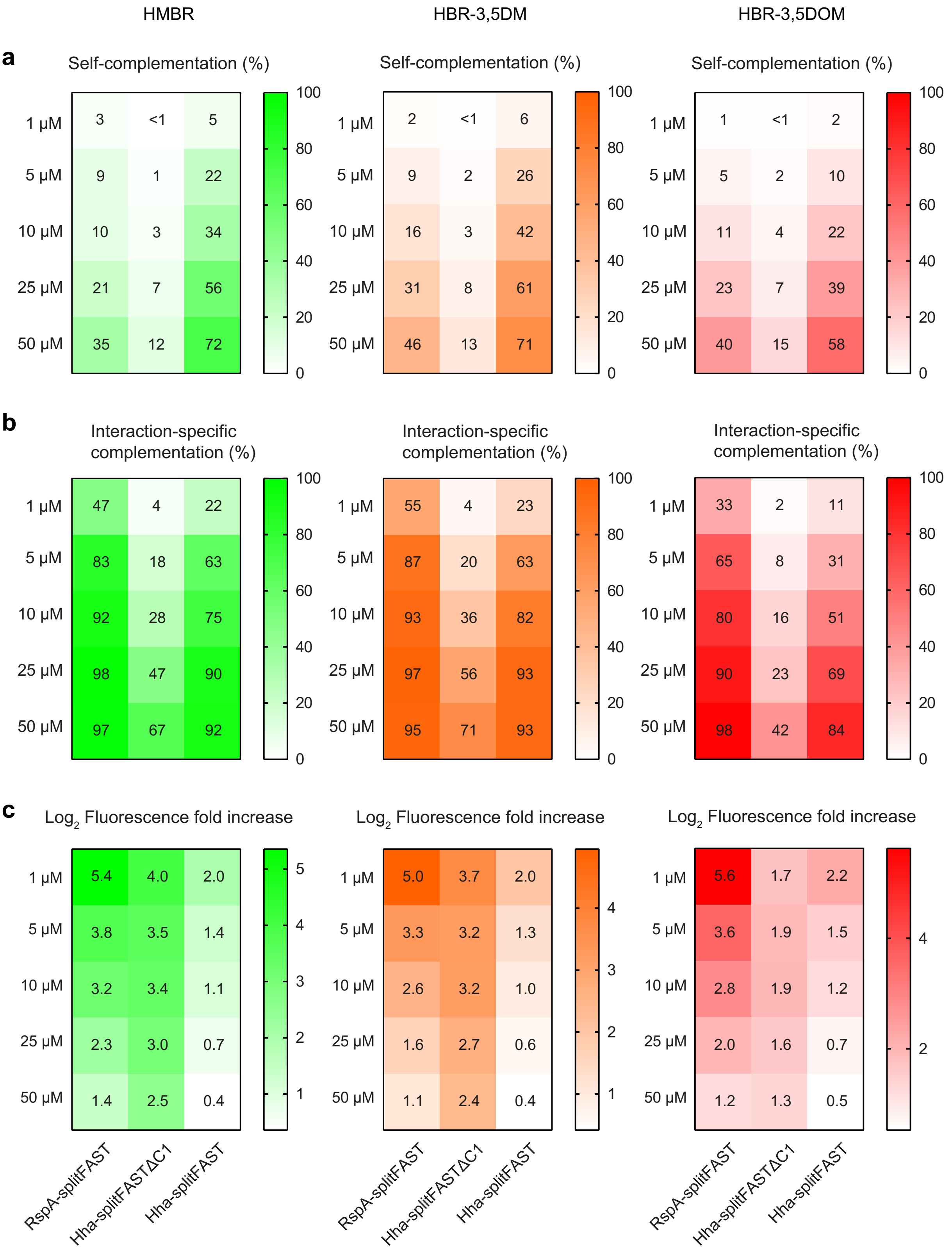


**Figure S4. Association properties of RspA-splitFAST, Hha-splitFAST and Hha-splitFASTΔC1. a** Percentage of self-complementation, **b** percentage of interaction-specific complementation, and **c** fluorescence fold increase (dynamic range) at various fluorogen concentrations (see also **Fig. S3**).

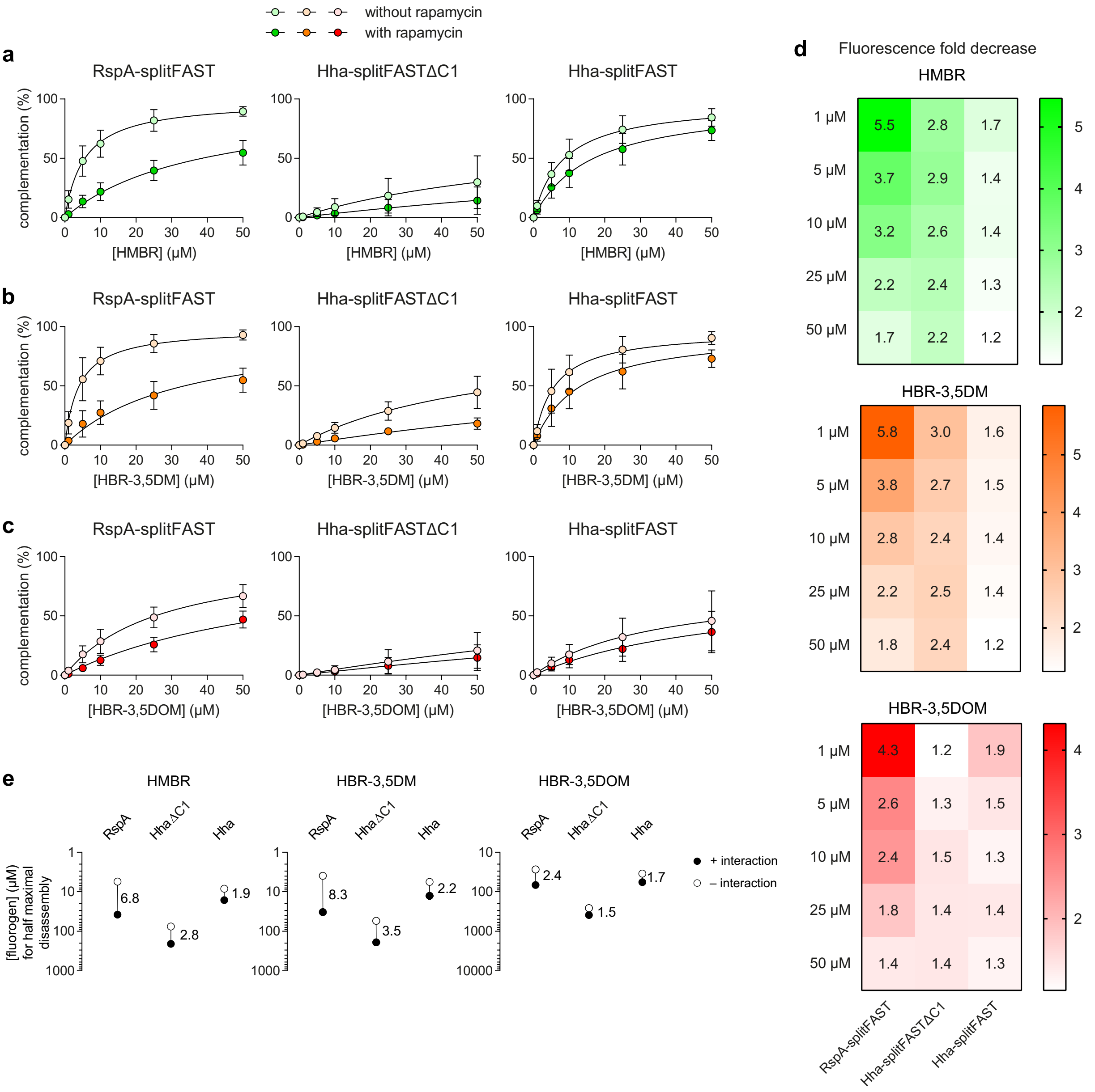


**Figure S5. Dissociation properties of RspA-splitFAST, Hha-splitFAST and Hha-splitFASTΔC1. a-c** Normalized average fluorescence of about 50,000 HEK293T cells co-expressing the homodimerizing FK506-binding protein F36M mutant ($FKBP_{F36M}$) fused to the N-terminal and C-terminal fragments of the split reporters treated without or with 500 nM of rapamycin, and with 1, 5, 10, 25 or 50 µM of HMBR(**a**), HBR-3,5DM (**b**) or HBR-3,5DOM (**c**). Data represent the mean ± standard deviation of three independent experiments. These data allowed to determine for each split reporter **d** the fluorescence fold decrease at various fluorogen concentrations. **e** Concentration of fluorogen for half maximal complementation in presence ($EC_{50,–interaction}$) or absence ($EC_{50,+interaction}$) of rapamycin (See also **Table S5**). The ratio $EC_{50,+interaction}$ / $EC_{50,–interaction}$ is indicated for each split reporter.

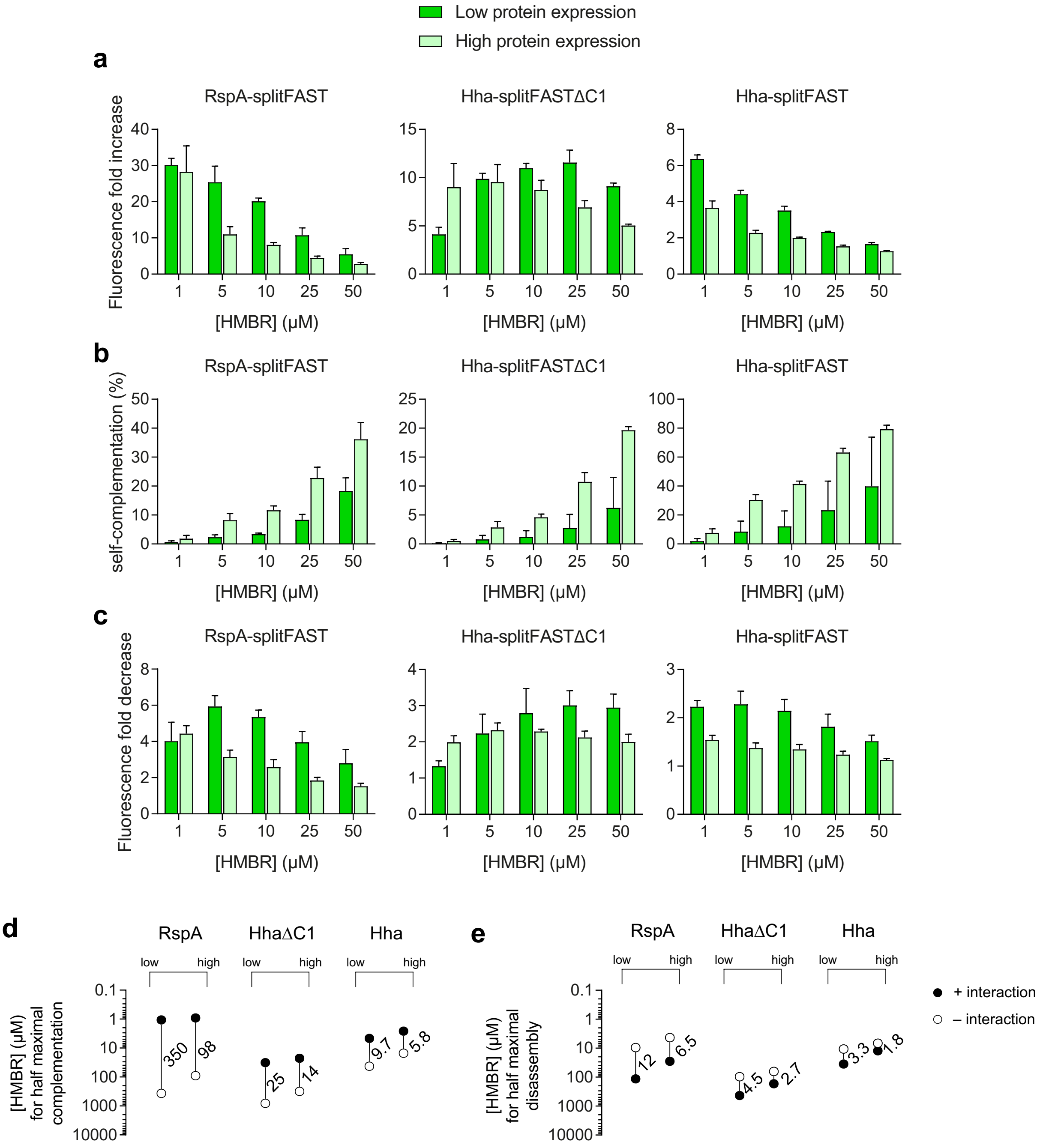


**Figure S6. Influence of the expression level on the complementation. a** Association dynamic range of RspA-splitFAST, Hha-splitFAST ΔC1, Hha-splitFAST at low (dark green) and high (light green) expression levels, and at various HMBR concentrations. **b** Self-complementation of RspA-splitFAST, Hha-splitFAST ΔC1, Hha-splitFAST at low (dark green) and high (light green) expression levels, and at various HMBR concentrations. **c** Dissociation dynamic range of RspA-splitFAST Hha-splitFAST ΔC1, Hha-splitFAST at low (dark green) and high (light green), and at various HMBR concentrations. Data represent the mean ± standard deviation of three independent experiments. mTurquoise2 and IRFP670 transfection reporters were used to define cell subgroups with low and high expression. **d** Concentration of fluorogen for half maximal complementation in presence ($EC_{50,+interaction}$) or absence ($EC_{50,–interaction}$) of rapamycin. The ratio $EC_{50,–interaction}$ / $EC_{50,+interaction}$ is indicated for each split reporter. **e** Concentration of fluorogen for half maximal complementation in presence ($EC_{50,–interaction}$) or absence ($EC_{50,+interaction}$) of rapamycin. The ratio $EC_{50,+interaction}$ / $EC_{50,–interaction}$ is indicated for each split reporter.

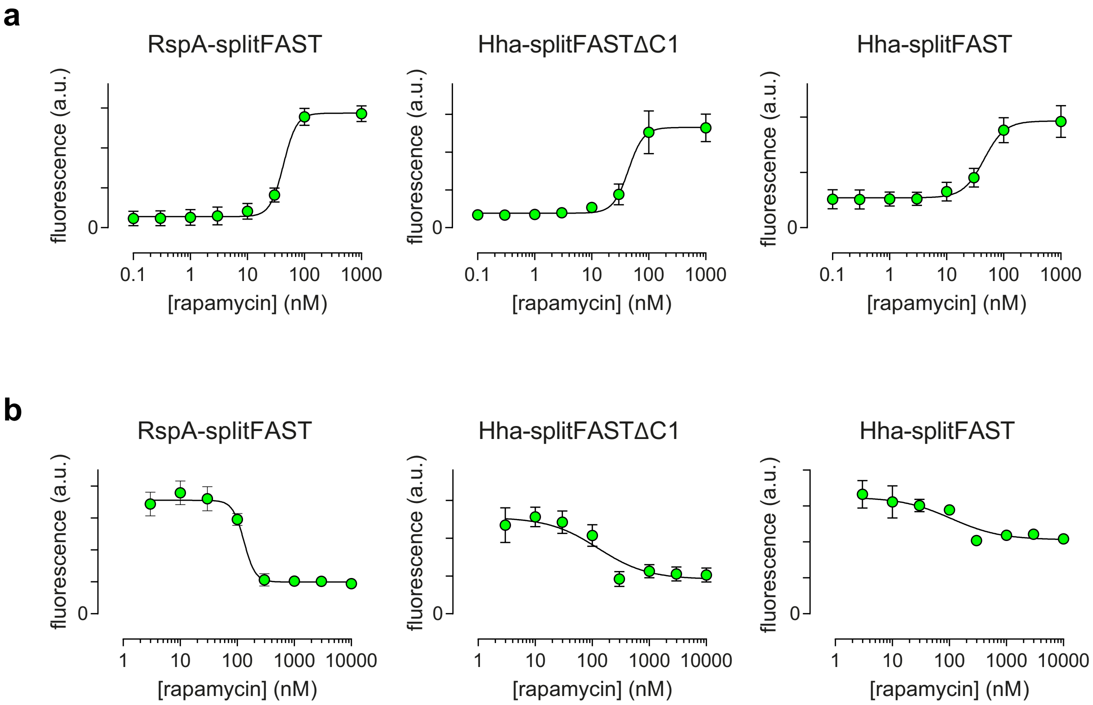


**Figure S7. Characterization of inducers and inhibitors of interactions. (a)** Fluorescence of about 50,000 HEK293T cells co-expressing the FK506-binding protein (FKBP) fused to the C-terminal fragment of the split reporters and the FKBP-rapamycin-binding domain of mammalian target of rapamycin (FRB) fused to the N-terminal fragment of the split reporters treated with 5 µM of HMBR and various concentrations of rapamycin. mTurquoise2 and IRP670 were used as transfection reporters. Data represent the mean ± standard deviation of three independent experiments. Least-square fit (line) gave the half-maximum effective concentration ($EC_{50}$) allowing to promote the FRB-FKBP interaction. **(b)** Fluorescence of about 50,000 HEK293T cells co-expressing the homodimerizing FK506-binding protein F36M mutant ($FKBP_{F36M}$) fused to the N-terminal and C-terminal fragments of the split reporters treated with 5 µM of HMBR and various concentrations of rapamycin. mTurquoise2 and IRP670 were used as transfection reporters. Data represent the mean ± standard deviation of three independent experiments. Least-square fit (line) gave the half-maximum effective concentration ($EC_{50}$) allowing to inhibit the $FKBP_{F36M}$-$FKBP_{F36M}$ interaction.

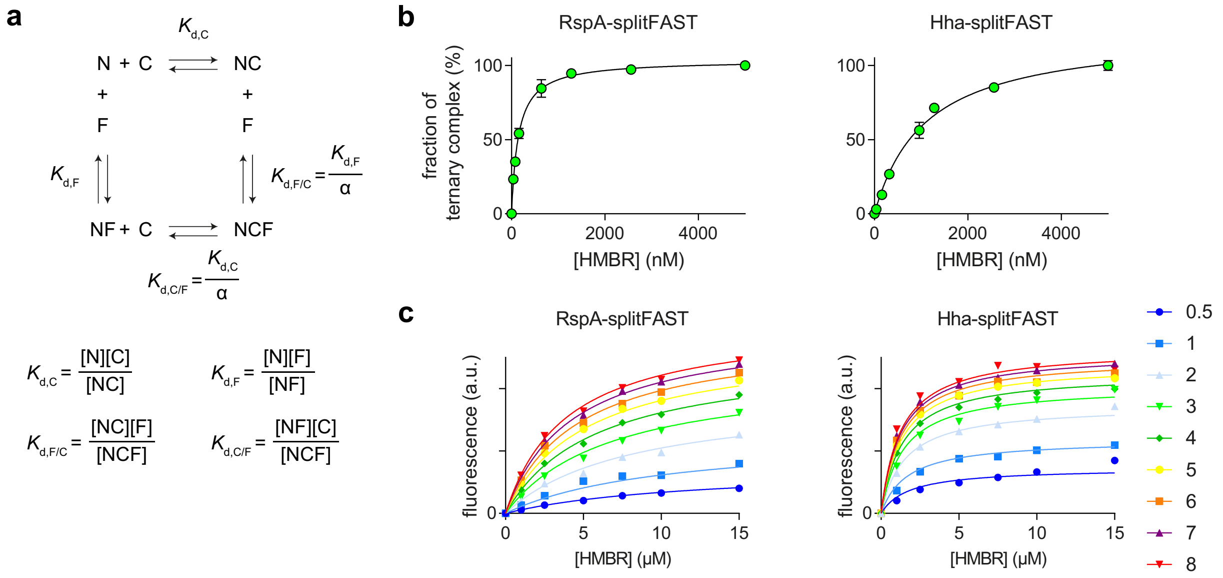


**d**

| | $K_{d,F}$ (μM) | $K_{d,C}$ (μM) | $K_{d,F/C}$ (μM) | $K_{d,C/F}$ (μM) | α |
|---|---|---|---|---|---|
| RspA-splitFAST | 21 ± 2 | 220 ± 10 | 0.15 ± 0.01 | 1.6 | 140 |
| Hha-splitFAST | 2.0 ± 0.2 | 3.1 ± 0.3 | 1.1 ± 0.1 | 1.6 | 2 |

**Figure S8. Thermodynamic characterization of Hha-splitFAST and RspA-splitFAST. a** Thermodynamic model of the self-complementation of the split reporters. **b** Purified recombinant proteins FRB-N and FKBP-C proteins (50 nM) in presence of 500 nM of rapamycin were treated with 0, 40, 80, 160, 320, 480, 640, 960, 1280, 2560 and 5000 nM of HMBR, and the fraction of complex was determined by measuring the fluorescence of the complex. Data represent the mean ± standard deviation of three independent experiments. Least-square fit gave the dissociation constants $K_{d,F/C}$. **c** Recombinant N-terminal fragment (50 nM) was treated with various concentrations of synthetic C-fragment (between 0.5 – 8 μM) and various concentations of HMBR. The evolution of the fluorescence intensity (in a.u.) is shown. Global least-square fit allowed the determination of $K_{d,F}$ and $K_{d,C}$, knowing $K_{d,F/C}$ (see **supplementary text 1** for details). **d** Extracted thermodynamic constants.

**Table S1.** Physico-chemical properties of FAST orthologs. Abbreviations are as follows: $\lambda_{abs}$, wavelength of maximal absorption; $\lambda_{em}$, wavelength of maximal emission; $\phi$, fluorescence quantum yield; $K_D$ thermodynamic dissociation constant.

| | fluorogen | $\lambda_{abs}$ (nm) | $\lambda_{em}$ (nm) | $\phi$ | $K_D$ ($\mu$M) |
|---|---|---|---|---|---|
| FAST | HMBR | 481 | 540 | 0.23 | 0.13 |
| | HBR-3,5DM | 499 | 562 | 0.49 | 0.08 |
| | HBR-3,5DOM | 518 | 600 | 0.31 | 0.97 |
| HboL-FAST | HMBR | 480 | 545 | 0.22 | 0.08 |
| | HBR-3,5DM | 500 | 569 | 0.53 | 0.043 |
| | HBR-3,5DOM | 525 | 598 | 0.25 | 0.33 |
| HspG-FAST | HMBR | 480 | 545 | 0.34 | 0.044 |
| | HBR-3,5DM | 500 | 567 | 0.41 | 0.014 |
| | HBR-3,5DOM | 526 | 604 | 0.33 | 0.18 |
| RspA-FAST | HMBR | 480 | 544 | 0.26 | 0.031 |
| | HBR-3,5DM | 499 | 572 | 0.45 | 0.011 |
| | HBR-3,5DOM | 527 | 601 | 0.42 | 0.074 |
| Ilo-FAST | HMBR | 481 | 546 | 0.35 | 0.23 |
| | HBR-3,5DM | 500 | 567 | 0.54 | 0.26 |
| | HBR-3,5DOM | 517 | 599 | 0.41 | 1.4 |
| TsiA-FAST | HMBR | 480 | 544 | 0.23 | 0.052 |
| | HBR-3,5DM | 500 | 572 | 0.53 | 0.008 |
| | HBR-3,5DOM | 519 | 603 | 0.34 | 0.060 |
| Rsa-FAST | HMBR | 476 | 545 | 0.14 | 0.17 |
| | HBR-3,5DM | 497 | 567 | 0.48 | 0.20 |
| | HBR-3,5DOM | 521 | 601 | 0.26 | 0.65 |

**Table S2. Comparison of the association efficiency of the orthologous split systems in cells.** Concentration of fluorogen for half maximal complementation in presence ($EC_{50,+interaction}$) or absence ($EC_{50,-interaction}$) of rapamycin (in μM). Data obtained by fitting the data of **Fig. 2a** and reported on **Fig 2e**.

| | | HboL-splitFAST | HspG-splitFAST | RspA-splitFAST | Ilo-splitFAST | TsiA-splitFAST | Rsa-splitFAST | Hha-splitFAST |
|---|---|---|---|---|---|---|---|---|
| $EC_{50,+interaction}$ | Value | 3.7 | 2.6 | 1.4 | 28 | 0.73 | 9.8 | 3.2 |
| | Std. Error | 0.2 | 0.2 | 0.1 | 2 | 0.04 | 0.3 | 0.3 |
| $EC_{50,-interaction}$ | Value | 4.2 | 45 | 100 | 190 | 42 | 180 | 19 |
| | Std. Error | 1 | 2 | 7 | 9 | 2 | 6 | 1 |

**Table S3. Comparison of the association efficiency of RspA-splitFAST, Hha-splitFAST and Hha-splitFASTΔC1 in cells.** Concentration of fluorogen for half maximal complementation in presence ($EC_{50,+interaction}$) or absence ($EC_{50,–interaction}$) of rapamycin (in µM). Data obtained by fitting the data of **Fig. S3a-c** and reported on **Fig. S3d**.

| **HMBR** | | RspA-splitFAST | Hha-splitFASTΔC1 | Hha-splitFAST |
|---|---|---|---|---|
| $EC_{50,+interaction}$ | Value | 1.0 | 26 | 3.1 |
| | Std. Error | 0.1 | 3 | 0.2 |
| $EC_{50,-interaction}$ | Value | 100 | 380 | 18 |
| | Std. Error | 8 | 50 | 1 |

| **HBR-3.5DM** | | RspA-splitFAST | Hha-splitFASTΔC1 | Hha-splitFAST |
|---|---|---|---|---|
| $EC_{50,+interaction}$ | Value | 0.78 | 18 | 2.8 |
| | Std. Error | 0.05 | 2 | 0.2 |
| $EC_{50,-interaction}$ | Value | 57 | 310 | 16 |
| | Std. Error | 2 | 40 | 1 |

| **HBR-3.5DOM** | | RspA-splitFAST | Hha-splitFASTΔC1 | Hha-splitFAST |
|---|---|---|---|---|
| $EC_{50,+interaction}$ | Value | 2.3 | 88 | 10 |
| | Std. Error | 0.2 | 29 | 1 |
| $EC_{50,-interaction}$ | Value | 77 | 320 | 39 |
| | Std. Error | 6 | 90 | 4 |

**Table S4. Comparison of the dissociation efficiency of the orthologous split systems in cells.** Concentration of fluorogen for half maximal complementation in absence ($EC_{50,+interaction}$) or presence ($EC_{50,–interaction}$) of rapamycin (in μM). Data obtained by fitting the data of **Fig. 2g** and reported on **Fig 2i**.

| | | HboL-splitFAST | HspG-splitFAST | RspA-splitFAST | Ilo-splitFAST | TsiA-splitFAST | Rsa-splitFAST | Hha-splitFAST |
|---|---|---|---|---|---|---|---|---|
| $EC_{50,+interaction}$ | Value | 21 | 22 | 38 | 120 | 58 | 78 | 16 |
| | Std. Error | 2 | 1 | 5 | 9 | 3 | 36 | 3 |
| $EC_{50,–interaction}$ | Value | 7.7 | 7.0 | 5.5 | 54 | 4.2 | 92 | 8.3 |
| | Std. Error | 0.8 | 0.4 | 0.9 | 4 | 0.3 | 41 | 1.6 |

**Table S5. Comparison of the dissociation efficiency of RspA-splitFAST, Hha-splitFAST and Hha-splitFASTΔC1 in cells.** Concentration of fluorogen for half maximal complementation in absence ($EC_{50,+interaction}$) or presence ($EC_{50,–interaction}$) of rapamycin (in μM). Data obtained by fitting the data of **Fig. S5a-c** and reported on **Fig. S5e**.

| **HMBR** | | RspA-splitFAST | Hha-splitFASTΔC1 | Hha-splitFAST |
|---|---|---|---|---|
| $EC_{50,+interaction}$ | Value | 38 | ~ 200 | 16 |
| | Std. Error | 5 | - | 3 |
| $EC_{50,–interaction}$ | Value | 5.5 | ~ 75 | 8.3 |
| | Std. Error | 0.9 | - | 1.6 |

| **HBR-3.5DM** | | RspA-splitFAST | Hha-splitFASTΔC1 | Hha-splitFAST |
|---|---|---|---|---|
| $EC_{50,+interaction}$ | Value | 33 | 190 | 13 |
| | Std. Error | 5 | 60 | 3 |
| $EC_{50,–interaction}$ | Value | 3.9 | 54 | 5.6 |
| | Std. Error | 0.8 | 21 | 1.3 |

| **HBR-3.5DOM** | | RspA-splitFAST | Hha-splitFASTΔC1 | Hha-splitFAST |
|---|---|---|---|---|
| $EC_{50,+interaction}$ | Value | 67 | ~ 380 | 57 |
| | Std. Error | 13 | - | 31 |
| $EC_{50,–interaction}$ | Value | 27 | ~ 260 | 34 |
| | Std. Error | 6 | - | 21 |

**Table S6. Comparison of the in-cell association efficiency of the chimera resulting from the fragment crossing of Hha-splitFAST and Rspa-splitFAST.** Concentration of fluorogen for half maximal complementation in presence ($EC_{50,+interaction}$) or absence ($EC_{50,-interaction}$) of rapamycin (in μM). Data obtained by fitting the data of **Fig. 4a** and reported on **Fig. 4e**.

| | | RspA-splitFAST | Hha-splitFAST | RspA(N)::Hha(C) | Hha(N)::RspA(C) |
|---|---|---|---|---|---|
| $EC_{50,+interaction}$ | Value | 1.6 | 4.9 | 5.1 | 2.1 |
| | Std. Error | 0.2 | 0.5 | 0.4 | 0.2 |
| $EC_{50,-interaction}$ | Value | 71 | 22 | 240 | 8.4 |
| | Std. Error | 9 | 2 | 22 | 0.7 |

**Table S7. Comparison of the in-cell dissociation efficiency of the chimera resulting from the fragment crossing of Hha-splitFAST and Rspa-splitFAST.** Concentration of fluorogen for half maximal complementation in absence ($EC_{50,+interaction}$) or presence ($EC_{50,–interaction}$) of rapamycin (in μM). Data obtained by fitting the data of **Fig. 4f** and reported on **Fig. 4h**.

| | | RspA-splitFAST | Hha-splitFAST | RspA(N)::Hha(C) | Hha(N)::RspA(C) |
|---|---|---|---|---|---|
| $EC_{50, + interaction}$ | Value | 45 | 39 | 200 | 16 |
| | Std. Error | 7 | 6 | 76 | 1 |
| $EC_{50, – interaction}$ | Value | 7.3 | 20 | 34 | 9.8 |
| | Std. Error | 1.4 | 4 | 6 | 0.7 |

### Supplementary Materials and Methods

### Molecular Biology

The plasmids pAG378-383 encoding each Histag-TEV-ortholog were constructed by Gibson assembly from the plasmid pAG87 encoding Hha-FAST. The sequences encoding each ortholog were ordered as g-blocks from Integrated DNA Technology (IDT). The backbone of pAG87 was amplified using primers ag321/KanF and ag322/KanR.

The plasmid pAG765, encoding myc-HboLFAST-IRES-mTurquoise2, was constructred by Gibson assembly from the plasmid pAG490(Tebo and Gautier, 2019) encoding FRB-Hha(N)-IRES-mTurquoise2. The sequence coding for HboLFAST was amplified by PCR from the plasmid pAG378 encoding HboL-FAST using primers ag1009/ag1010. The backbone of pAG490 was amplified in two fragments using primers ag412/ag313 and ag776/ag314. The three fragments were then assembled by Gibson assembly.

The plasmid pAG766, encoding myc-HspGFAST-IRES-mTurquoise2, was constructred by Gibson assembly from the plasmid pAG490(Tebo and Gautier, 2019) encoding FRB-Hha(N)-IRES-mTurquoise2. The sequence coding for HspGFAST was amplified by PCR from the plasmid pAG379 encoding HspG-FAST using primers ag1009/ag1011. The backbone of pAG490 was amplified in two fragments using primers ag412/ag313 and ag778/ag314. The three fragments were then assembled by Gibson assembly.

The plasmid pAG767, encoding myc-RspAFAST-IRES-mTurquoise2, was constructred by Gibson assembly from the plasmid pAG490(Tebo and Gautier, 2019) encoding FRB-Hha(N)-IRES-mTurquoise2. The sequence coding for RspAFAST was amplified by PCR from the plasmid pAG380 encoding RspA-FAST using primers ag1009/ag1012. The backbone of pAG490 was amplified in two fragments using primers ag412/ag313 and ag780/ag314. The three fragments were then assembled by Gibson assembly.

The plasmid pAG768, encoding myc-IloFAST-IRES-mTurquoise2, was constructred by Gibson assembly from the plasmid pAG490(Tebo and Gautier, 2019) encoding FRB-Hha(N)-IRES-mTurquoise2. The sequence coding for Ilo-FAST was amplified by PCR from the plasmid pAG381 encoding Ilo-FAST using primers ag1013/ag1014. The backbone of pAG490 was amplified in two fragments using primers ag412/ag313 and ag782/ag314. The three fragments were then assembled by Gibson assembly.

The plasmid pAG769, encoding myc-TsiAFAST-IRES-mTurquoise2, was constructred by Gibson assembly from the plasmid pAG490(Tebo and Gautier, 2019) encoding FRB-Hha(N)-IRES-mTurquoise2. The sequence coding for TsiA-FAST was amplified by PCR from the plasmid pAG382 encoding TsiA-FAST using primers ag1015/ag1016. The backbone of pAG490 was amplified in two fragments using primers ag412/ag313 and ag784/ag314. The three fragments were then assembled by Gibson assembly.

The plasmid pAG770, encoding myc-RsaFAST-IRES-mTurquoise2, was constructred by Gibson assembly from the plasmid pAG490(Tebo and Gautier, 2019) encoding

FRB-Hha(N)-IRES-mTurquoise2. The sequence coding for Rsa-FAST was amplified by PCR from the plasmid pAG383 encoding Rsa-FAST using primers ag1017/ag1018. The backbone of pAG490 was amplified in two fragments using primers ag412/ag313 and ag786/ag314. The three fragments were then assembled by Gibson assembly.

The plasmid pAG571, encoding FRB-HboL(N)-IRES-mTurquoise2, was constructed by Gibson assembly from the plasmid pAG490 (Tebo and Gautier, 2019) encoding FRB-Hha(N)-IRES-mTurquoise2. The sequence coding for HboL(N) was amplified by PCR from the plasmid pAG378 encoding HboL-FAST using primers ag766/ag767. The backbone of pAG490 was amplified in two fragments using primers ag374/ag313 and ag694/ag314. The three fragments were then assembled by Gibson assembly.

The plasmid pAG572, encoding FRB-HspG(N)-IRES-mTurquoise2, was constructed by Gibson assembly from the plasmid pAG490 (Tebo and Gautier, 2019) encoding FRB-Hha(N)-IRES-mTurquoise2. The sequence coding for HspG(N) was amplified by PCR from the plasmid pAG379 encoding HspG-FAST using primers ag766/ag768. The backbone of pAG490 was amplified in two fragments using primers ag374/ag313 and ag694/ag314. The three fragments were then assembled by Gibson assembly.

The plasmid pAG573, encoding FRB-RspA(N)-IRES-mTurquoise2, was constructed by Gibson assembly from the plasmid pAG490 (Tebo and Gautier, 2019) encoding FRB-Hha(N)-IRES-mTurquoise2. The sequence coding for RspA(N) was amplified by PCR from the plasmid pAG380 encoding RspA-FAST using primers ag766/ag769. The backbone of pAG490 was amplified in two fragments using primers ag374/ag313 and ag694/ag314. The three fragments were then assembled by Gibson assembly.

The plasmid pAG574, encoding FRB-Ilo(N)-IRES-mTurquoise2, was constructed by Gibson assembly from the plasmid pAG490 (Tebo and Gautier, 2019) encoding FRB-Hha(N)-IRES-mTurquoise2. The sequence coding for Ilo(N) was amplified by PCR from the plasmid pAG381 encoding Ilo-FAST using primers ag770/ag771. The backbone of pAG490 was amplified in two fragments using primers ag374/ag313 and ag694/ag314. The three fragments were then assembled by Gibson assembly.

The plasmid pAG575, encoding FRB-TsiA(N)-IRES-mTurquoise2, was constructed by Gibson assembly from the plasmid pAG490 (Tebo and Gautier, 2019) encoding FRB-Hha(N)-IRES-mTurquoise2. The sequence coding for TsiA(N) was amplified by PCR from the plasmid pAG382 encoding TsiA-FAST using primers ag772/ag773. The backbone of pAG490 was amplified in two fragments using primers ag374/ag313 and ag694/ag314. The three fragments were then assembled by Gibson assembly.

The plasmid pAG576, encoding FRB-Rsa(N)-IRES-mTurquoise2, was constructed by Gibson assembly from the plasmid pAG490(Tebo and Gautier, 2019) encoding FRB-Hha(N)-IRES-mTurquoise2. The sequence coding for Rsa(N) was amplified by PCR from the plasmid pAG383 encoding Rsa-FAST using primers ag774/ag775. The backbone of pAG490 was amplified in two fragments using primers ag374/ag313 and ag694/ag314. The three fragments were then assembled by Gibson assembly.

The plasmid pAG577 encoding FKBP-Hha(C)-IRES-iRFP670, was constructed by Gibson assembly from the plasmid pAG153(Tebo and Gautier, 2019) encoding CMV-cmyc-FKBP-linker-Hha(C). The sequence coding for Hha(C)-IRES was amplified by PCR from the plasmid pAG490 using primers ag694/ag695. The sequence coding for iRFP670 was amplified by PCR using primers ag763/ag764. The backbone of pAG153 was amplified using primers ag777/ag313. The backbone of pAG104 encoding for CMV-FAST540-human-cmyc was amplified using primers ag347/ag314. The four fragments were then assembled by Gibson assembly.

The plasmid pAG578, encoding FKBP-HboL(C)-IRES-iRFP670, was constructed by Gibson assembly from the plasmid pAG153 (Tebo and Gautier, 2019) encoding CMV-cmyc-FKBP-linker-Hha(C). The sequence coding for HboL(C)-IRES was amplified by PCR from the plasmid pAG490 using primers ag776/ag695. The sequence coding for iRFP670 was amplified by PCR using primers ag763/ag764. The backbone of pAG153 was amplified using primers ag777/ag313. The backbone of pAG104 encoding for CMV-FAST540-human-cmyc was amplified using primers ag347/ag314. The four fragments were then assembled by Gibson assembly.

The plasmid pAG579, encoding FKBP-HspG(C)-IRES-iRFP670, was constructed by Gibson assembly from the plasmid pAG153 (Tebo and Gautier, 2019) encoding CMV-cmyc-FKBP-linker-Hha(C). The sequence coding for HspG(C)-IRES was amplified by PCR from the plasmid pAG490 using primers ag778/ag695. The sequence coding for iRFP670 was amplified by PCR using primers ag763/ag764. The backbone of pAG153 was amplified using primers ag779/ag313. The backbone of pAG104 encoding for CMV-FAST540-human-cmyc was amplified using primers ag347/ag314. The four fragments were then assembled by Gibson assembly.

The plasmid pAG580, encoding FKBP-RspA(C)-IRES-iRFP670, was constructed by Gibson assembly from the plasmid pAG153 (Tebo and Gautier, 2019) encoding CMV-cmyc-FKBP-linker-Hha(C). The sequence coding for RspA(C)-IRES was amplified by PCR from the plasmid pAG490 using primers ag780/ag695. The sequence coding for iRFP670 was amplified by PCR using primers ag763/ag764. The backbone of pAG153 was amplified using primers ag781/ag313. The backbone of pAG104 encoding for CMV-FAST540-human-cmyc was amplified using primers ag347/ag314. The four fragments were then assembled by Gibson assembly.

The plasmid pAG581, encoding FKBP-Ilo(C)-IRES-iRFP670, was constructed by Gibson assembly from the plasmid pAG153 (Tebo and Gautier, 2019) encoding CMV-cmyc-FKBP-linker-(C). The sequence coding for Ilo(C)-IRES was amplified by PCR from the plasmid pAG490 using primers ag782/ag695. The sequence coding for iRFP670 was amplified by PCR using primers ag763/ag764. The backbone of pAG153 was amplified using primers ag783/ag313. The backbone of pAG104 encoding for CMV-FAST540-human-cmyc was amplified using primers ag347/ag314. The four fragments were then assembled by Gibson assembly.

The plasmid pAG582, encoding FKBP-TsiA(C)-IRES-iRFP670, was constructed by Gibson assembly from the plasmid pAG153 (Tebo and Gautier, 2019) encoding CMV-cmyc-FKBP-linker-Hha(C). The sequence coding for TsiA(C)-IRES was amplified by PCR from the plasmid pAG490 using primers ag784/ag695. The sequence coding for iRFP670 was amplified by PCR using primers ag763/ag764. The backbone of pAG153 was amplified using primers ag785/ag313. The backbone of pAG104 encoding for CMV-FAST540-human-cmyc was amplified using primers ag347/ag314. The four fragments were then assembled by Gibson assembly.

The plasmid pAG583, encoding FKBP-Rsa(C)-IRES-iRFP670, was constructed by Gibson assembly from the plasmid pAG153(Tebo and Gautier, 2019) encoding CMV-cmyc-FKBP-linker-Hha(C). The sequence coding for Rsa(C)-IRES was amplified by PCR from the plasmid pAG490 using primers ag786/ag695. The sequence coding for iRFP670 was amplified by PCR using primers ag763/ag764. The backbone of pAG153 was amplified using primers ag787/ag313. The backbone of pAG104 encoding for CMV-FAST540-human-cmyc was amplified using primers ag347/ag314. The four fragments were then assembled by Gibson assembly.

The plasmid pAG815, encoding $FKBP_{F36M}$-Ilo(C)-IRES-iRFP670, was constructed by Gibson assembly from the plasmid pAG581 encoding FKBP-Ilo(C)-IRES-iRFP670. The sequence coding for $FKBP_{F36M}$ was amplified by PCR using primers ag1039/ag1040. The backbone of pAG581 was amplified in two fragments using primers ag1041/ag313 and ag1042/ag314. The three fragments were then assembled by Gibson assembly.

The plasmid pAG804, encoding $FKBP_{F36M}$-HboL(N)-IRES-mTurquoise2, was constructed by subcloning the sequence coding for $FKBP_{F36M}$ (from pAG815) into the plasmid pAG571 using BglII and BspEI restriction sites.

The plasmid pAG805, encoding $FKBP_{F36M}$-HspG(N)-IRES-mTurquoise2, was constructed by subcloning the sequence coding for $FKBP_{F36M}$ (from pAG815) into the plasmid pAG572 using BglII and BspEI restriction sites.

The plasmid pAG806, encoding $FKBP_{F36M}$-RspA(N)-IRES-mTurquoise2, was constructed by subcloning the sequence coding for $FKBP_{F36M}$ (from pAG815) into the plasmid pAG572 using BglII and BspEI restriction sites.

The plasmid pAG807, encoding $FKBP_{F36M}$-Ilo(N)-IRES-mTurquoise2, was constructed by subcloning the sequence coding for $FKBP_{F36M}$ (from pAG815) into the plasmid pAG572 using BglII and BspEI restriction sites.

The plasmid pAG808, encoding $FKBP_{F36M}$-TsiA(N)-IRES-mTurquoise2, was constructed by subcloning the sequence coding for $FKBP_{F36M}$ (from pAG815) into the plasmid pAG572 using BglII and BspEI restriction sites.

The plasmid pAG809, encoding $FKBP_{F36M}$-Rsa(N)-IRES-mTurquoise2, was constructed by subcloning the sequence coding for $FKBP_{F36M}$ (from pAG815) into the plasmid pAG576 using BglII and BspEI restriction sites.

The plasmid pAG810, encoding $FKBP_{F36M}$-Hha(C)-IRES-iRFP670, was constructed by subcloning the sequence coding for $FKBP_{F36M}$ (from pAG815) into the plasmid pAG577 using BglII and BspEI restriction sites.

The plasmid pAG811, encoding $FKBP_{F36M}$-HhaΔC1(C)-IRES-iRFP670, was constructed by subcloning the sequence coding for $FKBP_{F36M}$ (from pAG815) into the plasmid pAG819 using BglII and BspEI restriction sites.

The plasmid pAG812, encoding $FKBP_{F36M}$-HboL(C)-IRES-iRFP670, was constructed by subcloning the sequence coding for $FKBP_{F36M}$ (from pAG815) into the plasmid pAG578 using BglII and BspEI restriction sites.

The plasmid pAG813, encoding $FKBP_{F36M}$-HspG(C)-IRES-iRFP670, was constructed by subcloning the sequence coding for $FKBP_{F36M}$ (from pAG815) into the plasmid pAG579 using BglII and BspEI restriction sites.

The plasmid pAG814, encoding $FKBP_{F36M}$-RspA(C)-IRES-iRFP670, was constructed by subcloning the sequence coding for $FKBP_{F36M}$ (from pAG815) into the plasmid pAG580 using BglII and BspEI restriction sites.

The plasmid pAG816, encoding $FKBP_{F36M}$-TsiA(C)-IRES-iRFP670, was constructed by subcloning the sequence coding for $FKBP_{F36M}$ (from pAG815) into the plasmid pAG582 using BglII and BspEI restriction sites.

The plasmid pAG817, encoding $FKBP_{F36M}$-Rsa(C)-IRES-iRFP670, was constructed by subcloning the sequence coding for $FKBP_{F36M}$ (from pAG815) into the plasmid pAG583 using BglII and BspEI restriction sites.

The plasmid pAG818, encoding $FKBP_{F36M}$-Hha(N)-IRES-mTurquoise2, was constructed by subcloning $FKBP_{F36M}$ (from pAG815) into the plasmid pAG490 using BglII and BspEI restriction sites.

The plasmid pAG819 encoding FKBP-HhaΔC1(C)-IRES-iRFP670, was constructed by Gibson assembly from the plasmid pAG577 encoding FKBP-Hha(C)-IRES-iRFP670. The sequence coding for HhaΔC1(C) was amplified by PCR using primers ag694/ag1019. The backbone of pAG577 was amplified using primers ag314/ag313. The three fragments were then assembled by Gibson assembly.

The plasmid pAG209 encoding for His-TEV-Hha(N) was reported previously (Tebo and Gautier, 2019).

The plasmid pAG820, His-TEV-RspA(N), was constructed by Gibson assembly from the plasmid pAG87(Plamont et al., 2016) encoding His-TEV-FAST. The backbone of pAG87 was amplified in two fragments using primers kanF/ag321 and ag322/kanR. The sequence coding for RspA(N) was amplified by PCR from the plasmid pAG573 using primers ag1133/ag1133. The three fragments were then assembled by Gibson assembly.

The plasmid pAG1015, His-TEV-FKBP-RspA(C), was constructed by Gibson assembly from the plasmid pAG820 encoding His-TEV-RspA(N). The backbone of pAG820 was amplified in two fragments using primers kanF/ag321 and ag322/kanR. The sequence coding for FKBP-RspA(C) was amplified by PCR from the plasmid pAG580 using primers ag1576/ag1577. The three fragments were then assembled by Gibson assembly.

The plasmid pAG1016, His-TEV-FRB-RspA(N), was constructed by Gibson assembly from the plasmid pAG820 encoding His-TEV-RspA(N). The backbone of pAG820 was amplified in two fragments using primers kanF/ag321 and ag322/kanR. The sequence coding for FRB-RspA(N) was amplified by PCR from the plasmid pAG573 using primers ag1576/ag1580. The three fragments were then assembled by Gibson assembly.

The plasmid pAG1017, His-TEV-FKBP-Hha(C), was constructed by Gibson assembly from the plasmid pAG820 encoding His-TEV-RspA(N). The backbone of pAG820 was amplified in two fragments using primers kanF/ag321 and ag322/kanR. The sequence coding for FKBP-Hha(C) was amplified by PCR from the plasmid pAG577 using primers ag1576/ag1580. The three fragments were then assembled by Gibson assembly.

The plasmid pAG1118, His-TEV-FRB-Hha(N), was constructed by Gibson assembly from the plasmid pAG820 encoding His-TEV-RspA(N). The backbone of pAG820 was amplified in two fragments using primers kanF/ag321 and ag322/kanR. The sequence coding for FRB-Hha(N) was amplified by PCR from the plasmid pAG577 using primers ag1578/ag1581. The three fragments were then assembled by Gibson assembly.
The plasmids used in these studies allowed the expression of proteins in mammalian cells. Expression is under the control of a CMV promoter.

The plasmid pAG892, encoding RspA(N)-H3-IRES-mTurquoise2, was constructed by Gibson assembly from the plasmid pAG891 encoding RspA(C)-H3-IRES-mTurquoise2. The backbone of pAG891 was amplified in two fragments using primers ag1353/ag313 and ag1063/ag314. The sequence coding for RspA(N) was amplified by PCR from the plasmid pAG573 encoding using primers ag1354/ag1355. The three fragments were then assembled by Gibson assembly.

The plasmid pAG895, encoding HP1-RspA(C)-IRES-iRFP670, was constructed by Gibson assembly from the plasmid pAG573 and pAG580 encoding FRB-RspA(N)-IRES-mTurquoise2 and FKBP-RspA(C)-IRES-iRFP670, respectively. The backbone of pAG573 was amplified using primers ag1353/ag313 and pAG580 was amplified using primers ag314/ag1357. The two fragments were then assembled with a synthetic gBlock encoding for HP1-RspA(C) by Gibson assembly.

The plasmid pAG1049, encoding HP1W45A-RspA(C)-IRES-iRFP670, was constructed by Gibson assembly from the plasmid pAG895 encoding HP1-RspA(C)-IRES-iRFP670. The backbone of pAG895 was amplified in two fragments using primers ag1610/ag313 and ag1609/ag314. The two fragments were then assembled by Gibson assembly.

The plasmid pAG1143, encoding RspA(N)-MEK1, was constructed by Gibson assembly from the plasmid pAG298(Tebo and Gautier, 2019) encoding Hha(N)-MEK1. The backbone of pAG298 was amplified in two fragments using primers ag1776/ag313 and ag1777/ag314. The sequence coding for RspA(N) was amplified by PCR from the plasmid pAG573 encoding FRB-RspA(N)-IRES-mTurquoise2 using primers ag1746/ag1146. The three fragments were then assembled by Gibson assembly.

The plasmid pAG1144, encoding lyn11-FRB-RspA(N), was constructed by Gibson assembly from the plasmid pAG336(Tebo and Gautier, 2019) encoding lyn11-FRB-Hha(N). The backbone of pAG336 was amplified in two fragments using primers ag1778/ag313 and ag347/ag314. The sequence coding for RspA(N) was amplified by PCR from the plasmid pAG573 encoding FRB-RspA(N)-IRES-mTurquoise2 using primers ag1779/ag1780. The three fragments were then assembled by Gibson assembly.

The plasmid pAG1176, encoding mCherry-ERK2-RspA(C), was constructed by Gibson assembly from the plasmid pAG297(Tebo and Gautier, 2019) encoding mCherry-ERK2-Hha(C). The backbone of pAG297 was amplified in two fragments using primers ag1829/ag313 and ag1830/ag314. The two fragments were then assembled by Gibson assembly.

The plasmid pAG1177, encoding FKBP-RspA(C), was constructed by Gibson assembly from the plasmid pAG153(Tebo and Gautier, 2019) encoding FKBP-Hha(C). The backbone of pAG153 was amplified in two fragments using primers ag1829/ag313 and ag1830/ag314. The two fragments were then assembled by Gibson assembly.

**Table S8.** Sequences of the primers used in this study

| Primer | Sequence |
|---|---|
| kanF | gcatcaaccaaaccgttattcattcgtg |
| kanR | cacgaatgaataacggtttggttgatgc |
| ag313 | ctcaccttgctcctgccgagaaagtatcca |
| ag314 | tggatactttctcggcaggagcaaggtgag |
| ag321 | gccctgaaaatacaggttttcgctagc |
| ag322 | taatagctcgagcaccaccaccac |
| ag347 | taataggcggccgcgactctag |
| ag374 | ggatccccctccgccgctgccgcctcctccgga |
| ag412 | caagtcctcttcagaaataagcttttgttc |
| ag694 | taactcgaggactacaaggacgacg |
| ag695 | ctcctcgcccttgctcaccatgaattcagcgtaatctggaacatcgtatg |
| ag763 | catacgatgttccagattacgctgaattcatggcgcgtaaggtcgatctcac |
| ag764 | ctagagtcgcggccgcctattagcgttggtggtgggcgg |
| ag766 | gcagcggcggagggggatccatggagaccgtgagattcggcg |
| ag767 | cgtcgtccttgtagtcctcgagttagctgatggccttcttcatgtgcac |
| ag768 | cgtcgtccttgtagtcctcgagttagctcagggccttcttcatgtgc |
| ag769 | cgtcgtccttgtagtcctcgagttaggtcatggccttcttcatgtgcac |
| ag770 | gcagcggcggagggggatccatggagatcgtgcagttcggc |
| ag771 | cgtcgtccttgtagtcctcgagttaggtcagggccttcttcatgtgc |
| ag772 | gcagcggcggagggggatccatggagctgctgagcttcgg |
| ag773 | cgtcgtccttgtagtcctcgagttacaccagggccttcttcatgtgc |
| ag774 | gcagcggcggagggggatccatggagatgatcaagttcggccag |
| ag775 | cgtcgtccttgtagtcctcgagttaggtgatggctctcttcatgtgcac |
| ag776 | cacctactggatcttcgtgaagagactgtaactcgaggactacaaggacgacg |
| ag777 | ctcttcacgaagatccagtaggtgtcgccggatccccctccgccgc |
| ag778 | caccttctggatcttcgtgaagagactgtaactcgaggactacaaggacgacg |
| ag779 | ctcttcacgaagatccagaaggtgtcgccggatccccctccgccgc |
| ag780 | cagcttctggatcttcgtgaagagactgtaactcgaggactacaaggacgacg |
| ag781 | cttcacgaagatccagaagctgtcgccggatccccctccgccgc |
| ag782 | cacctactgggtgttcgtgaagagactgtaactcgaggactacaaggacgacg |
| ag783 | cttcacgaacacccagtaggtgtcgccggatccccctccgccgc |
| ag784 | cacctactgggtgttcgtgaagagagtgtaactcgaggactacaaggacgacg |
| ag785 | cttcacgaacacccagtaggtgtcgtcggatccccctccgccgc |
| ag786 | cagctactggatcttcgtgaagagagtgtaactcgaggactacaaggacgacg |
| ag787 | cttcacgaagatccagtagctgtcgccggatccccctccgccgc |
| ag1009 | caaaagcttatttctgaagaggacttggaattcgagaccgtgagattcggcgg |
| ag1010 | cagtctcttcacgaagatccagtaggtg |
| ag1011 | cagtctcttcacgaagatccagaaggtg |
| ag1012 | cagtctcttcacgaagatccagaagc |
| ag1013 | caaaagcttatttctgaagaggacttggaattcgagatcgtgcagttcggcagc |
| ag1014 | cagtctcttcacgaacacccagtagg |
| ag1015 | caaaagcttatttctgaagaggacttggaattcgagctgctgagcttcggcg |
| ag1016 | cactctcttcacgaacacccagtagg |
| ag1017 | caaaagcttatttctgaagaggacttggaattcgagatgatcaagttcggccaggacg |
| ag1018 | cactctcttcacgaagatccagtagctg |
| ag1019 | cgtcgtccttgtagtcctcgagttaccgtttcacaaagacccaatagc |
| ag1039 | ctgaagaggacttggaattcggagtgcaggtggaaaccatc |
| ag1040 | ccgctgccgcctcctccggattcttccagttttagaagctcc |
| ag1041 | gatggtttccacctgcactccgaattccaagtcctcttcag |
| ag1042 | gagcttctaaaactggaagaatccggaggaggcggcag |
| ag1063 | tccggaggaggcggcag |
| ag1132 | gaaaacctgtattttcagggcatggagaccgtgagattcgg |
| ag1133 | gtggtgctcgagctattaggtcatggccttcttcatgtgc |
| ag1146 | ggtcatggccttcttcatg |
| ag1353 | catggtggcagatctgagtccg |
| ag1354 | ccggactcagatctgccaccatggagaccgtgagattcggcgg |

| | |
|---|---|
| ag1355 | ctgccgcctcctccggaggtcatggccttcttcatgtgc |
| ag1576 | gaaaacctgtattttcagggcggagtgcaggtggaaaccatc |
| ag1577 | gtggtgctcgagctattacagtctcttcacgaagatccag |
| ag1578 | gaaaacctgtattttcagggcgagatgtggcatgaaggcctg |
| ag1580 | gtggtgctcgagctattacacccgtttcacaaagaccc |
| ag1581 | gtggtgctcgagctattagacctgctttgagattcgtcgg |
| ag1609 | gagtactatctgaaagccaagggctatcccg |
| ag1610 | cgggatagcccttggctttcagatagtactc |
| ag1746 | atggagaccgtgagattcggcg |
| ag1776 | cgaatctcacggtctccatggtggcagatctgagtccggtag |
| ag1777 | cacatgaagaaggccatgacctccggaggaggcggcagcg |
| ag1778 | cctccggagacctgctttgagattc |
| ag1779 | ctcaaagcaggtctccggaggaggcggcagcggcggagggggatccatggagac |
| ag1780 | agagtcgcggccgcctattaggtcatggccttcttcatgtgc |
| ag1829 | cacgaagatccagaagctgtcgccggatccccctccgccgc |
| ag1830 | gatcttcgtgaagagactgtaataggcggccgcgactc |

**Table S9.** Plasmids developed in this study

| Plasmid | Encoded proteins | References |
|---|---|---|
| pAG209 | His-Hha(N) | (Tebo and Gautier, 2019) |
| pAG490 | FRB-Hha(N)-IRES-mTurquoise2 | (Tebo and Gautier, 2019) |
| pAG378 | His-HboLFAST | |
| pAG379 | His-HspGFAST | |
| pAG380 | His-RspAFAST | |
| pAG381 | His-IloFAST | |
| pAG382 | His-TsiAFAST | |
| pAG383 | His-RsaFAST | |
| pAG571 | FRB-HboL(N)-IRES-mTurquoise2 | |
| pAG572 | FRB-HspG(N)-IRES-mTurquoise2 | |
| pAG573 | FRB-RspA(N)-IRES-mTurquoise2 | |
| pAG574 | FRB-Ilo(N)-IRES-mTurquoise2 | |
| pAG575 | FRB-TsiA(N)-IRES-mTurquoise2 | |
| pAG576 | FRB-Rsa(N)-IRES-mTurquoise2 | |
| pAG577 | FKBP-Hha(C)-IRES-iRFP670 | |
| pAG578 | FKBP-HboL(C)-IRES-iRFP670 | |
| pAG579 | FKBP- HspG(C)-IRES-iRFP670 | |
| pAG580 | FKBP-RspA(C)-IRES-iRFP670 | |
| pAG581 | FKBP-Ilo(C)-IRES-iRFP670 | |
| pAG582 | FKBP-TsiA(C)-IRES-iRFP670 | |
| pAG583 | FKBP-Rsa(C)-IRES-iRFP670 | |
| pAG765 | Myc-HboLFAST-IRES-mTurquoise2 | |
| pAG766 | Myc-HspGFAST-IRES-mTurquoise2 | |
| pAG767 | Myc-RspAFAST-IRES-mTurquoise2 | |
| pAG768 | Myc-IloFAST-IRES-mTurquoise2 | |
| pAG769 | Myc-TsiAFAST-IRES-mTurquoise2 | |
| pAG770 | Myc-RsaFAST-IRES-mTurquoise2 | |
| pAG804 | $FKBP_{F36M}$-HboL(N)-IRES-mTurquoise2 | |
| pAG805 | $FKBP_{F36M}$-HspG(N)-IRES-mTurquoise2 | |
| pAG806 | $FKBP_{F36M}$-RspA(N)-IRES-mTurquoise2 | |
| pAG807 | $FKBP_{F36M}$-Ilo(N)-IRES-mTurquoise2 | |
| pAG808 | $FKBP_{F36M}$-TsiA(N)-IRES-mTurquoise2 | |
| pAG809 | $FKBP_{F36M}$-Rsa(N)-IRES-mTurquoise2 | |
| pAG810 | $FKBP_{F36M}$-Hha(C)-IRES-iRFP670 | |
| pAG811 | $FKBP_{F36M}$- HhaΔC1(C)-IRES-iRFP670 | |
| pAG812 | $FKBP_{F36M}$-HboL(C)-IRES-iRFP670 | |
| pAG813 | $FKBP_{F36M}$-HspG(C)-IRES-iRFP670 | |

| | |
|---|---|
| pAG814 | $FKBP_{F36M}$-RspA(C)-IRES-iRFP670 |
| pAG815 | $FKBP_{F36M}$-Ilo-C-IRES-iRFP670 |
| pAG816 | $FKBP_{F36M}$-TsiA(C)-IRES-iRFP670 |
| pAG817 | $FKBP_{F36M}$-Rsa(C)-IRES-iRFP670 |
| pAG818 | $FKBP_{F36M}$-Hha(N)-IRES-mTurquoise2 |
| pAG819 | FKBP-HhaΔC1(C)-IRES-iRFP670 |
| pAG820 | His-TEV-RspA(N) |
| pAG892 | RspA(N)-H3-IRES-mTurquoise2 |
| pAG895 | HP1-RspA(C)-IRES-iRFP670 |
| pAG1015 | His-TEV-FKBP-RspA(C) |
| pAG1016 | His-TEV-FKBP-RspA(N) |
| pAG1017 | His-TEV-FKBP-Hha(C) |
| pAG1049 | HP1W45A-RspA(C)-IRES-iRFP670 |
| pAG1118 | His-TEV-FRB-Hha(N) |
| pAG1143 | RspA(N)-MEK1 |
| pAG1144 | lyn11-FRB-RspA(N) |
| pAG1176 | mCherry-ERK2-RspA(C) |
| pAG1177 | FKBP-RspA(C) |

**Table S10.** Sequences

| Vector | ORF | ORF sequence |
| --- | --- | --- |
| | Hha(N) | atggagcatgttgcctttggcagtgaggacatcgagaacactctggccaaaatggacgacg<br>gacaactggatgggttggcctttggcgcaattcagctcgatggtgacgggaatatcctgcagt<br>acaatgctgctgaaggagacatcacaggcagagatcccaaacaggtgattgggaagaac<br>ttcttcaaggatgttgcacctggaacggattctcccgagttttacggcaaattcaaggaaggcg<br>tagcgtcagggaatctgaacaccatgttcgaatggatgataccgacaagcaggggaccaa<br>ccaaggtcaaggtgcacatgaagaaagccctttcc |
| | HboL(N) | atggagaccgtgagattcggcggcgacgacatcgagaacagcctggccaagatggacga<br>caagaagctggacgagctggccttcggcgccatccagctggacgccaacggcaagatcat<br>ccagtacaacgccgccgagggcggcatcaccggcagagaccccaagagcgtgatcggc<br>aagaacttcttcaccgaggtggcccccggcacccagagcaaggagttccagggcagattc<br>aaggagggcgtgagcagcggcgagctgaacaccatgttcgagtggatgatccccaccag<br>cagaggccccaccaaggtgaaggtgcacatgaagaaggccatcagc |
| | HspG(N) | atggagaccgtgagattcggcggcgacgacatcgagaacgccctggccaacatggacga<br>caagaagctggacaccctggccttcggcgccatccagctggacgccaacggcaagatcat<br>ccagtacaacgccgccgagggcggcatcaccggcagagaccccaagagcgtgatcggc<br>aagaacttcttcaccgacgtggcccccggcacccagagcaaggagttccagggcagattc<br>aaggagggcgtgaagaacggcgacctgaacaccatgttcgagtggatgatccccaccag<br>cagaggccccaccaaggtgaaggtgcacatgaagaaggccctgagc |
| | RspA(N) | atggagaccgtgagattcggcggcgacgacatcgagaacagcctggccaagatggacga<br>caaggccctggacaagctggccttcggcgccatccagctggacggcaacggcaagatcat<br>ccactacaacgccgccgagggcaccatcaccggcagagaccccaagaccgtgatcggc<br>aagaacttcttcaccgacgtggcccccggcacccagagcaaggagttccagggcagattc<br>aaggagggcgtgcagaagggcgacctgaacaccatgttcgagtggatgatccccaccag<br>cagaggccccaccaaggtgaaggtgcacatgaagaaggccatgacc |
| | Ilo(N) | atggagatcgtgcagttcggcagcgacgacatcgagaacaccctgagcaagatgagcga<br>cgacaagctgaacgacatcgccttcggcgccatccagctggacgccagcggcaagatcat<br>ccagtacaacgccgccgagggcgacatcaccggcagagaccccggcgccgtggtgggc<br>aagaacttcttcaacgaggtggcccccggcaccaacagccccgagttcaagggcagattc<br>gacgagggcgtgaagaacggcaacctgaacaccatgttcgagtggatgatccccaccag<br>cagaggccccaccaaggtgaaggtgcacatgaagaaggccctgacc |
| | TsiA(N) | atggagctgctgagcttcggcgccgacaacatcgagaacagcctggccaagatgagcaa<br>gggcgacctgaacaagctggccttcggcgccatccagctgaacgcccagggcaagatcct<br>gcagtacaacgccgccgagggcgacatcaccggcagaaagcccaccgaggtgatcggc<br>aagaacttcttcctggaggtggcccccggcaccaacagaaccgagttcaagggcagattc<br>gaccagggcatcaagagcggcaacctgaacaccatgttcgagtggatgatccccaccagc<br>agaggccccaccaaggtgaaggtgcacatgaagaaggccctggtg |
| | Rsa(N) | atggagatgatcaagttcggccaggacgacatcgagaacgccatggccgacatgggcga<br>cgcccagatcgacgacctggccttcggcgccatccagctggacgagaccggcaccatcct<br>ggcctacaacgccgccgagggcgagctgaccggcagaagcccccaggacgtgatcggc<br>aagaacttcttcaaggacatcgcccccggcaccgacaccgaggagttcggcggcagattc<br>agagagggcgtggccaacggcgacctgaacgccatgttcgagtggatgatccccaccag<br>cagaggccccaccaaggtgaaggtgcacatgaagagagccatcacc |
| | Hha(C) | ggtgacagctattgggtctttgtgaaacgggtg |
| | HhaΔC1(C) | ggtgacagctattgggtctttgtgaaacgg |
| | HboL(C) | ggcgacacctactggatcttcgtgaagagactg |

| | | |
|---|---|---|
| | HspG(C) | ggcgacaccttctggatcttcgtgaagagactg |
| | RspA(C) | ggcgacagcttctggatcttcgtgaagagactg |
| | Ilo(C) | ggcgacacctactgggtgttcgtgaagagactg |
| | TsiA(C) | gacgacacctactgggtgttcgtgaagagagtg |
| | Rsa(C) | ggcgacagctactggatcttcgtgaagagagtg |
| pAG209 | His-tag-thrombin-Hha(N) | atgggcagcagccatcatcatcatcatcacagcagcggcctggtgccgcgcggcagccat<br>atggctagcatggagcatgttgcctttggcagtgaggacatcgagaacactctggccaaaat<br>ggacgacggacaactggatgggttggcctttggcgcaattcagctcgatggtgacgggaat<br>atcctgcagtacaatgctgctgaaggagacatcacaggcagagatcccaaacaggtgattg<br>ggaagaacttcttcaaggatgttgcacctggaacggattctcccgagttttacggcaaattcaa<br>ggaaggcgtagcgtcagggaatctgaacaccatgttcgaatggatgataccgacaagcag<br>gggaccaaccaaggtcaaggtgcacatgaagaaagccctttcctaa |
| pAG490 | C-myc-FRB-Hha(N) | atggaacaaaagcttatttctgaagaggacttggaattcgagatgtggcatgaaggcctgga<br>agaggcatctcgtttgtactttggggaaaggaacgtgaaaggcatgtttgaggtgctggagcc<br>cttgcatgctatgatggaacggggcccccagactctgaaggaaacatcctttaatcaggcct<br>atggtcgagatttaatggaggcccaagagtggtgcaggaagtacatgaaatcagggaatgt<br>caaggacctcacccaagcctgggacctctattatcatgtgttccgacgaatctcaaagcaggt<br>ctccggaggaggcggcagcggcggagggggatccatggagcatgttgcctttggcagtga<br>ggacatcgagaacactctggccaaaatggacgacggacaactggatgggttggcctttggc<br>gcaattcagctcgatggtgacgggaatatcctgcagtacaatgctgctgaaggagacatcac<br>aggcagagatcccaaacaggtgattgggaagaacttcttcaaggatgttgcacctggaacg<br>gattctcccgagttttacggcaaattcaaggaaggcgtagcgtcagggaatctgaacaccat<br>gttcgaatggatgataccgacaagcaggggaccaaccaaggtcaaggtgcacatgaaga<br>aagccctttcctaa |
| pAG571 | C-myc-FRB-HboL(N) | atggaacaaaagcttatttctgaagaggacttggaattcgagatgtggcatgaaggcctgga<br>agaggcatctcgtttgtactttggggaaaggaacgtgaaaggcatgtttgaggtgctggagcc<br>cttgcatgctatgatggaacggggcccccagactctgaaggaaacatcctttaatcaggcct<br>atggtcgagatttaatggaggcccaagagtggtgcaggaagtacatgaaatcagggaatgt<br>caaggacctcacccaagcctgggacctctattatcatgtgttccgacgaatctcaaagcaggt<br>ctccggaggaggcggcagcggcggagggggatccatggagaccgtgagattcggcggc<br>gacgacatcgagaacagcctggccaagatggacgacaagaagctggacgagctggcctt<br>cggcgccatccagctggacgccaacggcaagatcatccagtacaacgccgccgagggc<br>ggcatcaccggcagagaccccaagagcgtgatcggcaagaacttcttcaccgaggtggcc<br>cccggcacccagagcaaggagttccagggcagattcaaggagggcgtgagcagcggcg<br>agctgaacaccatgttcgagtggatgatccccaccagcagaggccccaccaaggtgaag<br>gtgcacatgaagaaggccatcagctaa |

| | | |
|---|---|---|
| pAG572 | C-myc-FRB-HspG(N) | atggaacaaaagcttatttctgaagaggacttggaattcgagatgtggcatgaaggcctggaagaggcatctcgtttgtactttggggaaaggaacgtgaaaggcatgtttgaggtgctggagcccttgcatgctatgatggaacggggcccccagactctgaaggaaacatcctttaatcaggcctatggtcgagatttaatggaggcccaagagtggtgcaggaagtacatgaaatcagggaatgtcaaggacctcacccaagcctgggacctctattatcatgtgttccgacgaatctcaaagcaggtctccggaggaggcggcagcggcggagggggatccatggagaccgtgagattcggcggcgacgacatcgagaacgccctggccaacatggacgacaagaagctggacaccctggccttcggcgccatccagctggacgccaacggcaagatcatccagtacaacgccgccgagggcggcatcaccggcagagaccccaagagcgtgatcggcaagaacttcttcaccgacgtggccccggcacccagagcaaggagttccagggcagattcaaggagggcgtgaagaacggcgacctgaacaccatgttcgagtggatgatccccaccagcagaggccccaccaaggtgaaggtgcacatgaagaaggccctgagctaa |
| pAG573 | C-myc-FRB-RspA(N) | atggaacaaaagcttatttctgaagaggacttggaattcgagatgtggcatgaaggcctggaagaggcatctcgtttgtactttggggaaaggaacgtgaaaggcatgtttgaggtgctggagcccttgcatgctatgatggaacggggcccccagactctgaaggaaacatcctttaatcaggcctatggtcgagatttaatggaggcccaagagtggtgcaggaagtacatgaaatcagggaatgtcaaggacctcacccaagcctgggacctctattatcatgtgttccgacgaatctcaaagcaggtctccggaggaggcggcagcggcggagggggatccatggagaccgtgagattcggcggcgacgacatcgagaacagcctggccaagatggacgacaaggccctggacaagctggccttcggcgccatccagctggacggcaacggcaagatcatccactacaacgccgccgagggcaccatcaccggcagagaccccaagaccgtgatcggcaagaacttcttcaccgacgtggccccggcacccagagcaaggagttccagggcagattcaaggagggcgtgcagaagggcgacctgaacaccatgttcgagtggatgatccccaccagcagaggccccaccaaggtgaaggtgcacatgaagaaggccatgacctaa |
| pAG574 | C-myc-FRB-Ilo(N) | atggaacaaaagcttatttctgaagaggacttggaattcgagatgtggcatgaaggcctggaagaggcatctcgtttgtactttggggaaaggaacgtgaaaggcatgtttgaggtgctggagcccttgcatgctatgatggaacggggcccccagactctgaaggaaacatcctttaatcaggcctatggtcgagatttaatggaggcccaagagtggtgcaggaagtacatgaaatcagggaatgtcaaggacctcacccaagcctgggacctctattatcatgtgttccgacgaatctcaaagcaggtctccggaggaggcggcagcggcggagggggatccatggagatcgtgcagttcggcagcgacgacatcgagaacaccctgagcaagatgagcgacgacaagctgaacgacatcgccttcggcgccatccagctggacgccagcggcaagatcatccagtacaacgccgccgagggcgacatcaccggcagagaccccggcgccgtggtgggcaagaacttcttcaacgaggtggcccccggcaccaacagccccgagttcaagggcagattcgacgagggcgtgaagaacggcaacctgaacaccatgttcgagtggatgatccccaccagcagaggccccaccaaggtgaaggtgcacatgaagaaggccctgacctaa |
| pAG575 | C-myc-FRB-TsiA(N) | atggaacaaaagcttatttctgaagaggacttggaattcgagatgtggcatgaaggcctggaagaggcatctcgtttgtactttggggaaaggaacgtgaaaggcatgtttgaggtgctggagcccttgcatgctatgatggaacggggcccccagactctgaaggaaacatcctttaatcaggcctatggtcgagatttaatggaggcccaagagtggtgcaggaagtacatgaaatcagggaatgtcaaggacctcacccaagcctgggacctctattatcatgtgttccgacgaatctcaaagcaggtctccggaggaggcggcagcggcggagggggatccatggagctgctgagcttcggcgccgacaacatcgagaacagcctggccaagatgagcaagggcgacctgaacaagctggccttcggcgccatccagctgaacgcccagggcaagatcctgcagtacaacgccgccgagggcgacatcaccggcagaaagcccaccgaggtgatcggcaagaacttcttcctggaggtggcccccggcaccaacagaaccgagttcaagggcagattcgaccagggcatcaagagcggcaacctgaacaccatgttcgagtggatgatccccaccagcagaggccccaccaaggtgaaggtgcacatgaagaaggccctggtgtaa |
| pAG576 | C-myc-FRB-Rsa(N) | atggaacaaaagcttatttctgaagaggacttggaattcgagatgtggcatgaaggcctggaagaggcatctcgtttgtactttggggaaaggaacgtgaaaggcatgtttgaggtgctggagcccttgcatgctatgatggaacggggcccccagactctgaaggaaacatcctttaatcaggcctatggtcgagatttaatggaggcccaagagtggtgcaggaagtacatgaaatcagggaatgtcaaggacctcacccaagcctgggacctctattatcatgtgttccgacgaatctcaaagcaggtctccggaggaggcggcagcggcggagggggatccatggagatgatcaagttcggccaggacgacatcgagaacgccatggccgacatgggcgacgcccagatcgacgacctggccttcggcgccatccagctggacgagaccggcaccatcctggcctacaacgccgccgagggcgagctgaccggcagaagcccccaggacgtgatcggcaagaacttcttcaaggacatcgccccggcaccgacaccgaggagttcggcggcagattcagagagggcgtggccaacggcgacctgaacgccatgttcgagtggatgatccccaccagcagaggccccaccaaggtgaaggtgcacatgaagagagccatcacctaa |

| | | |
|---|---|---|
| pAG577 | C-myc-FKBP-Hha(C) | atggaacaaaagcttatttctgaagaggacttggaattcggagtgcaggtggaaaccatctccccaggagacgggcgcaccttccccaagcgcggccagacctgcgtggtgcactacaccgggatgcttgaagatggaaagaaatttgattcctcccgggacagaaacaagccctttaagtttatgctaggcaagcaggaggtgatccgaggctgggaagaaggggttgcccagatgagtgtgggtcagagagccaaactgactatatctccagattatgcctatggtgccactgggcacccaggcatcatcccaccacatgccactctcgtcttcgatgtggagcttctaaaactggaagaatccggaggaggcggcagcggcggagggggatccggtgacagctattgggtctttgtgaaacgggtgtaa |
| pAG578 | C-myc-FKBP-HboL(C) | atggaacaaaagcttatttctgaagaggacttggaattcggagtgcaggtggaaaccatctccccaggagacgggcgcaccttccccaagcgcggccagacctgcgtggtgcactacaccgggatgcttgaagatggaaagaaatttgattcctcccgggacagaaacaagccctttaagtttatgctaggcaagcaggaggtgatccgaggctgggaagaaggggttgcccagatgagtgtgggtcagagagccaaactgactatatctccagattatgcctatggtgccactgggcacccaggcatcatcccaccacatgccactctcgtcttcgatgtggagcttctaaaactggaagaatccggaggaggcggcagcggcggagggggatccggcgacacctactggatcttcgtgaagagactgtaa |
| pAG579 | C-myc-FKBP-HspG(C) | atggaacaaaagcttatttctgaagaggacttggaattcggagtgcaggtggaaaccatctccccaggagacgggcgcaccttccccaagcgcggccagacctgcgtggtgcactacaccgggatgcttgaagatggaaagaaatttgattcctcccgggacagaaacaagccctttaagtttatgctaggcaagcaggaggtgatccgaggctgggaagaaggggttgcccagatgagtgtgggtcagagagccaaactgactatatctccagattatgcctatggtgccactgggcacccaggcatcatcccaccacatgccactctcgtcttcgatgtggagcttctaaaactggaagaatccggaggaggcggcagcggcggagggggatccggcgacaccttctggatcttcgtgaagagactgtaa |
| pAG580 | C-myc-FKBP-RspA(C) | atggaacaaaagcttatttctgaagaggacttggaattcggagtgcaggtggaaaccatctccccaggagacgggcgcaccttccccaagcgcggccagacctgcgtggtgcactacaccgggatgcttgaagatggaaagaaatttgattcctcccgggacagaaacaagccctttaagtttatgctaggcaagcaggaggtgatccgaggctgggaagaaggggttgcccagatgagtgtgggtcagagagccaaactgactatatctccagattatgcctatggtgccactgggcacccaggcatcatcccaccacatgccactctcgtcttcgatgtggagcttctaaaactggaagaatccggaggaggcggcagcggcggagggggatccggcgacagcttctggatcttcgtgaagagactgtaa |
| pAG581 | C-myc-FKBP-Ilo(C) | atggaacaaaagcttatttctgaagaggacttggaattcggagtgcaggtggaaaccatctccccaggagacgggcgcaccttccccaagcgcggccagacctgcgtggtgcactacaccgggatgcttgaagatggaaagaaatttgattcctcccgggacagaaacaagccctttaagtttatgctaggcaagcaggaggtgatccgaggctgggaagaaggggttgcccagatgagtgtgggtcagagagccaaactgactatatctccagattatgcctatggtgccactgggcacccaggcatcatcccaccacatgccactctcgtcttcgatgtggagcttctaaaactggaagaatccggaggaggcggcagcggcggagggggatccggcgacacctactgggtgttcgtgaagagactgtaa |
| pAG582 | C-myc-FKBP-TsiA(C) | atggaacaaaagcttatttctgaagaggacttggaattcggagtgcaggtggaaaccatctccccaggagacgggcgcaccttccccaagcgcggccagacctgcgtggtgcactacaccgggatgcttgaagatggaaagaaatttgattcctcccgggacagaaacaagccctttaagtttatgctaggcaagcaggaggtgatccgaggctgggaagaaggggttgcccagatgagtgtgggtcagagagccaaactgactatatctccagattatgcctatggtgccactgggcacccaggcatcatcccaccacatgccactctcgtcttcgatgtggagcttctaaaactggaagaatccggaggaggcggcagcggcggagggggatccgacgacacctactgggtgttcgtgaagagagtgtaa |
| pAG583 | C-myc-FKBP-Rsa(C) | atggaacaaaagcttatttctgaagaggacttggaattcggagtgcaggtggaaaccatctccccaggagacgggcgcaccttccccaagcgcggccagacctgcgtggtgcactacaccgggatgcttgaagatggaaagaaatttgattcctcccgggacagaaacaagccctttaagtttatgctaggcaagcaggaggtgatccgaggctgggaagaaggggttgcccagatgagtgtgggtcagagagccaaactgactatatctccagattatgcctatggtgccactgggcacccaggcatcatcccaccacatgccactctcgtcttcgatgtggagcttctaaaactggaagaatccggaggaggcggcagcggcggagggggatccggcgacagctactggatcttcgtgaagagagtgtaa |

| | | |
|---|---|---|
| pAG804 | C-myc-FKBP$_{F36M}$-HboL(N) | atggaacaaaagcttatttctgaagaggacttggaattcggagtgcaggtggaaaccatctc cccaggagacgggcgcaccttccccaagcgcggccagacctgcgtggtgcactacaccg ggatgcttgaagatggaaagaaaatggattcctcccgggacagaaacaagccctttaagttt atgctaggcaagcaggaggtgatccgaggctgggaagaaggggttgcccagatgagtgt gggtcagagagccaaactgactatatctccagattatgcctatggtgccactgggcacccag gcatcatcccaccacatgccactctcgtcttcgatgtggagcttctaaaactggaagaatccg gaggaggcggcagcggcggagggggatccatggagaccgtgagattcggcggcgacg acatcgagaacagcctggccaagatggacgacaagaagctggacgagctggccttcggc gccatccagctggacgccaacggcaagatcatccagtacaacgccgccgagggcggcat caccggcagagaccccaagagcgtgatcggcaagaacttcttcaccgaggtggccccg gcacccagagcaaggagttccagggcagattcaaggagggcgtgagcagcggcgagct gaacaccatgttcgagtggatgatccccaccagcagaggccccaccaaggtgaaggtgc acatgaagaaggccatcagctaa |
| pAG805 | C-myc-FKBP$_{F36M}$-HspG(N) | atggaacaaaagcttatttctgaagaggacttggaattcggagtgcaggtggaaaccatctc cccaggagacgggcgcaccttccccaagcgcggccagacctgcgtggtgcactacaccg ggatgcttgaagatggaaagaaaatggattcctcccgggacagaaacaagccctttaagttt atgctaggcaagcaggaggtgatccgaggctgggaagaaggggttgcccagatgagtgt gggtcagagagccaaactgactatatctccagattatgcctatggtgccactgggcacccag gcatcatcccaccacatgccactctcgtcttcgatgtggagcttctaaaactggaagaatccg gaggaggcggcagcggcggagggggatccatggagaccgtgagattcggcggcgacg acatcgagaacgccctggccaacatggacgacaagaagctggacaccctggccttcggc gccatccagctggacgccaacggcaagatcatccagtacaacgccgccgagggcggcat caccggcagagaccccaagagcgtgatcggcaagaacttcttcaccgacgtggccccg gcacccagagcaaggagttccagggcagattcaaggagggcgtgaagaacggcgacct gaacaccatgttcgagtggatgatccccaccagcagaggccccaccaaggtgaaggtgc acatgaagaaggccctgagctaa |
| pAG806 | C-myc-FKBP$_{F36M}$-RspA(N) | atggaacaaaagcttatttctgaagaggacttggaattcggagtgcaggtggaaaccatctc cccaggagacgggcgcaccttccccaagcgcggccagacctgcgtggtgcactacaccg ggatgcttgaagatggaaagaaaatggattcctcccgggacagaaacaagccctttaagttt atgctaggcaagcaggaggtgatccgaggctgggaagaaggggttgcccagatgagtgt gggtcagagagccaaactgactatatctccagattatgcctatggtgccactgggcacccag gcatcatcccaccacatgccactctcgtcttcgatgtggagcttctaaaactggaagaatccg gaggaggcggcagcggcggagggggatccatggagaccgtgagattcggcggcgacg acatcgagaacagcctggccaagatggacgacaaggccctggacaagctggccttcggc gccatccagctggacggcaacggcaagatcatccactacaacgccgccgagggcaccat caccggcagagaccccaagaccgtgatcggcaagaacttcttcaccgacgtggccccg gcacccagagcaaggagttccagggcagattcaaggagggcgtgcagaagggcgacct gaacaccatgttcgagtggatgatccccaccagcagaggccccaccaaggtgaaggtgc acatgaagaaggccatgacctaa |
| pAG807 | C-myc-FKBP$_{F36M}$-Ilo(N) | atggaacaaaagcttatttctgaagaggacttggaattcggagtgcaggtggaaaccatctc cccaggagacgggcgcaccttccccaagcgcggccagacctgcgtggtgcactacaccg ggatgcttgaagatggaaagaaaatggattcctcccgggacagaaacaagccctttaagttt atgctaggcaagcaggaggtgatccgaggctgggaagaaggggttgcccagatgagtgt gggtcagagagccaaactgactatatctccagattatgcctatggtgccactgggcacccag gcatcatcccaccacatgccactctcgtcttcgatgtggagcttctaaaactggaagaatccg gaggaggcggcagcggcggagggggatccatggagatcgtgcagttcggcagcgacga catcgagaacaccctgagcaagatgagcgacgacaagctgaacgacatcgccttcggcg ccatccagctggacgccagcggcaagatcatccagtacaacgccgccgagggcgacatc accggcagagaccccggcgccgtggtgggcaagaacttcttcaacgaggtggcccccgg caccaacagccccgagttcaagggcagattcgacgagggcgtgaagaacggcaacctg aacaccatgttcgagtggatgatccccaccagcagaggccccaccaaggtgaaggtgcac atgaagaaggccctgacctaa |

| | | |
|---|---|---|
| pAG808 | C-myc-<br>FKBP$_{F36M}$-<br>TsiA(N) | atggaacaaaagcttatttctgaagaggacttggaattcggagtgcaggtggaaaccatctc<br>cccaggagacgggcgcaccttccccaagcgcggccagacctgcgtggtgcactacaccg<br>ggatgcttgaagatggaaagaaaatggattcctcccgggacagaaacaagccctttaagttt<br>atgctaggcaagcaggaggtgatccgaggctgggaagaaggggttgcccagatgagtgt<br>gggtcagagagccaaactgactatatctccagattatgcctatggtgccactgggcacccag<br>gcatcatcccaccacatgccactctcgtcttcgatgtggagcttctaaaactggaagaatccg<br>gaggaggcggcagcggcggagggggatccatggagctgctgagcttcggcgccgacaa<br>catcgagaacagcctggccaagatgagcaagggcgacctgaacaagctggccttcggcg<br>ccatccagctgaacgcccagggcaagatcctgcagtacaacgccgccgagggcgacatc<br>accggcagaaagcccaccgaggtgatcggcaagaacttcttcctggaggtggcccccggc<br>accaacagaaccgagttcaagggcagattcgaccagggcatcaagagcggcaacctga<br>acaccatgttcgagtggatgatccccaccagcagaggccccaccaaggtgaaggtgcaca<br>tgaagaaggccctggtgtaa |
| pAG809 | C-myc-<br>FKBP$_{F36M}$-<br>Rsa(N) | atggaacaaaagcttatttctgaagaggacttggaattcggagtgcaggtggaaaccatctc<br>cccaggagacgggcgcaccttccccaagcgcggccagacctgcgtggtgcactacaccg<br>ggatgcttgaagatggaaagaaaatggattcctcccgggacagaaacaagccctttaagttt<br>atgctaggcaagcaggaggtgatccgaggctgggaagaaggggttgcccagatgagtgt<br>gggtcagagagccaaactgactatatctccagattatgcctatggtgccactgggcacccag<br>gcatcatcccaccacatgccactctcgtcttcgatgtggagcttctaaaactggaagaatccg<br>gaggaggcggcagcggcggagggggatccatggagatgatcaagttcggccaggacga<br>catcgagaacgccatggccgacatgggcgacgcccagatcgacgacctggccttcggcg<br>ccatccagctggacgagaccggcaccatcctggcctacaacgccgccgagggcgagctg<br>accggcagaagcccccaggacgtgatcggcaagaacttcttcaaggacatcgcccccgg<br>caccgacaccgaggagttcggcggcagattcagagagggcgtggccaacggcgacctg<br>aacgccatgttcgagtggatgatccccaccagcagaggccccaccaaggtgaaggtgcac<br>atgaagagagccatcacctaa |
| pAG810 | C-myc-<br>FKBP$_{F36M}$-<br>Hha(C) | atggaacaaaagcttatttctgaagaggacttggaattcggagtgcaggtggaaaccatctc<br>cccaggagacgggcgcaccttccccaagcgcggccagacctgcgtggtgcactacaccg<br>ggatgcttgaagatggaaagaaaatggattcctcccgggacagaaacaagccctttaagttt<br>atgctaggcaagcaggaggtgatccgaggctgggaagaaggggttgcccagatgagtgt<br>gggtcagagagccaaactgactatatctccagattatgcctatggtgccactgggcacccag<br>gcatcatcccaccacatgccactctcgtcttcgatgtggagcttctaaaactggaagaatccg<br>gaggaggcggcagcggcggagggggatccggtgacagctattgggtctttgtgaaacggg<br>tgtaa |
| pAG811 | C-myc-<br>FKBP$_{F36M}$-<br>HhaΔC1(C) | atggaacaaaagcttatttctgaagaggacttggaattcggagtgcaggtggaaaccatctc<br>cccaggagacgggcgcaccttccccaagcgcggccagacctgcgtggtgcactacaccg<br>ggatgcttgaagatggaaagaaaatggattcctcccgggacagaaacaagccctttaagttt<br>atgctaggcaagcaggaggtgatccgaggctgggaagaaggggttgcccagatgagtgt<br>gggtcagagagccaaactgactatatctccagattatgcctatggtgccactgggcacccag<br>gcatcatcccaccacatgccactctcgtcttcgatgtggagcttctaaaactggaagaatccg<br>gaggaggcggcagcggcggagggggatccggtgacagctattgggtctttgtgaaacggt<br>aa |
| pAG812 | C-myc-<br>FKBP$_{F36M}$-<br>HboL(C) | atggaacaaaagcttatttctgaagaggacttggaattcggagtgcaggtggaaaccatctc<br>cccaggagacgggcgcaccttccccaagcgcggccagacctgcgtggtgcactacaccg<br>ggatgcttgaagatggaaagaaaatggattcctcccgggacagaaacaagccctttaagttt<br>atgctaggcaagcaggaggtgatccgaggctgggaagaaggggttgcccagatgagtgt<br>gggtcagagagccaaactgactatatctccagattatgcctatggtgccactgggcacccag<br>gcatcatcccaccacatgccactctcgtcttcgatgtggagcttctaaaactggaagaatccg<br>gaggaggcggcagcggcggagggggatccggcgacacctactggatcttcgtgaagag<br>actgtaa |
| pAG813 | C-myc-<br>FKBP$_{F36M}$-<br>HspG(C) | atggaacaaaagcttatttctgaagaggacttggaattcggagtgcaggtggaaaccatctc<br>cccaggagacgggcgcaccttccccaagcgcggccagacctgcgtggtgcactacaccg<br>ggatgcttgaagatggaaagaaaatggattcctcccgggacagaaacaagccctttaagttt<br>atgctaggcaagcaggaggtgatccgaggctgggaagaaggggttgcccagatgagtgt<br>gggtcagagagccaaactgactatatctccagattatgcctatggtgccactgggcacccag<br>gcatcatcccaccacatgccactctcgtcttcgatgtggagcttctaaaactggaagaatccg<br>gaggaggcggcagcggcggagggggatccggcgacaccttctggatcttcgtgaagaga<br>ctgtaa |

| | | |
|---|---|---|
| pAG814 | C-myc-FKBP$_{F36M}$-RspA(C) | atggaacaaaagcttatttctgaagaggacttggaattcggagtgcaggtggaaaccatctccccaggagacgggcgcaccttccccaagcgcggccagacctgcgtggtgcactacaccgggatgcttgaagatggaaagaaaatggattcctcccgggacagaaacaagccctttaagtttatgctaggcaagcaggaggtgatccgaggctgggaagaaggggttgcccagatgagtgtgggtcagagagccaaactgactatatctccagattatgcctatggtgccactgggcacccaggcatcatcccaccacatgccactctcgtcttcgatgtggagcttctaaaactggaagaatccggaggaggcggcagcggcggaggggggatccggcgacagcttctggatcttcgtgaagagactgtaa |
| pAG815 | C-myc-FKBP$_{F36M}$-Ilo(C) | atggaacaaaagcttatttctgaagaggacttggaattcggagtgcaggtggaaaccatctccccaggagacgggcgcaccttccccaagcgcggccagacctgcgtggtgcactacaccgggatgcttgaagatggaaagaaaatggattcctcccgggacagaaacaagccctttaagtttatgctaggcaagcaggaggtgatccgaggctgggaagaaggggttgcccagatgagtgtgggtcagagagccaaactgactatatctccagattatgcctatggtgccactgggcacccaggcatcatcccaccacatgccactctcgtcttcgatgtggagcttctaaaactggaagaatccggaggaggcggcagcggcggaggggggatccggcgacacctactgggtgttcgtgaagagactgtaa |
| pAG816 | C-myc-FKBP$_{F36M}$-TsiA(C) | atggaacaaaagcttatttctgaagaggacttggaattcggagtgcaggtggaaaccatctccccaggagacgggcgcaccttccccaagcgcggccagacctgcgtggtgcactacaccgggatgcttgaagatggaaagaaaatggattcctcccgggacagaaacaagccctttaagtttatgctaggcaagcaggaggtgatccgaggctgggaagaaggggttgcccagatgagtgtgggtcagagagccaaactgactatatctccagattatgcctatggtgccactgggcacccaggcatcatcccaccacatgccactctcgtcttcgatgtggagcttctaaaactggaagaatccggaggaggcggcagcggcggaggggggatccgacgacacctactgggtgttcgtgaagagagtgtaa |
| pAG817 | C-myc-FKBP$_{F36M}$-Rsa(C) | atggaacaaaagcttatttctgaagaggacttggaattcggagtgcaggtggaaaccatctccccaggagacgggcgcaccttccccaagcgcggccagacctgcgtggtgcactacaccgggatgcttgaagatggaaagaaaatggattcctcccgggacagaaacaagccctttaagtttatgctaggcaagcaggaggtgatccgaggctgggaagaaggggttgcccagatgagtgtgggtcagagagccaaactgactatatctccagattatgcctatggtgccactgggcacccaggcatcatcccaccacatgccactctcgtcttcgatgtggagcttctaaaactggaagaatccggaggaggcggcagcggcggaggggggatccggcgacagctactggatcttcgtgaagagagtgtaa |
| pAG818 | C-myc-FKBP$_{F36M}$-Hha(N) | atggaacaaaagcttatttctgaagaggacttggaattcggagtgcaggtggaaaccatctccccaggagacgggcgcaccttccccaagcgcggccagacctgcgtggtgcactacaccgggatgcttgaagatggaaagaaaatggattcctcccgggacagaaacaagccctttaagtttatgctaggcaagcaggaggtgatccgaggctgggaagaaggggttgcccagatgagtgtgggtcagagagccaaactgactatatctccagattatgcctatggtgccactgggcacccaggcatcatcccaccacatgccactctcgtcttcgatgtggagcttctaaaactggaagaatccggaggaggcggcagcggcggaggggggatccatggagcatgttgcctttggcagtgaggacatcgagaacactctggccaaaatggacgacggacaactggatgggttggcctttggcgcaattcagctcgatggtgacgggaatatcctgcagtacaatgctgctgaaggagacatcacaggcagagatcccaaacaggtgattgggaagaacttcttcaaggatgttgcacctggaacggattctcccgagttttacggcaaattcaaggaaggcgtagcgtcagggaatctgaacaccatgttcgaatggatgataccgacaagcaggggaccaaccaaggtcaaggtgcacatgaagaaagcccttttcctaa |
| pAG819 | C-myc-FKBP-HhaΔC1(C) | atggaacaaaagcttatttctgaagaggacttggaattcggagtgcaggtggaaaccatctccccaggagacgggcgcaccttccccaagcgcggccagacctgcgtggtgcactacaccgggatgcttgaagatggaaagaaatttgattcctcccgggacagaaacaagccctttaagtttatgctaggcaagcaggaggtgatccgaggctgggaagaaggggttgcccagatgagtgtgggtcagagagccaaactgactatatctccagattatgcctatggtgccactgggcacccaggcatcatcccaccacatgccactctcgtcttcgatgtggagcttctaaaactggaagaatccggaggaggcggcagcggcggaggggggatccggtgacagctattgggtctttgtgaaacggtaa |
| pAG820 | His-Tag-thrombin-TEV-RspA(N) | atgggcagcagccatcatcatcatcatcacagcagcggcctggtgccgcgcggcagccatatggctagcgaaaacctgtattttcagggcatggagaccgtgagattcggcggcgacgacatcgagaacagcctggccaagatggacgacaaggccctggacaagctggccttcggcgccatccagctggacggcaacggcaagatcatccactacaacgccgccgagggcaccatcaccggcagagaccccaagaccgtgatcggcaagaacttcttcaccgacgtggcccccggcacccagagcaaggagttccagggcagattcaaggagggcgtgcagaagggcgacctgaacaccatgttcgagtggatgatccccaccagcagaggccccaccaaggtgaaggtgcacatgaagaaggccatgacctaa |

| | | |
|---|---|---|
| pAG892 | RspA(N)-H3 | atggagaccgtgagattcggcggcgacgacatcgagaacagcctggccaagatggacgacaaggccctggacaagctggccttcggcgccatccagctggacggcaacggcaagatcatccactacaacgccgccgagggcaccatcaccggcagagaccccaagaccgtgatcggcaagaacttcttcaccgacgtggcccccggcacccagagcaaggagttccagggcagattcaaggagggcgtgcagaagggcgacctgaacaccatgttcgagtggatgatccccaccagcagaggccccaccaaggtgaaggtgcacatgaagaaggccatgacctccggaggaggcggcagcggcggaggggggatccatggcacgaaccaagcagactgcccgcaaatccaccggcggtaaggcaccacggaagcagctggctaccaaagctgctagaaagagtgctcctgcaacaggggggagtgaaaaagcctcaccgatatcgccctgggactgtggcccttagggagatcagaagataccaaaaaagcacagagttgctgatccggaagctgccatttcaacgactggtgcgggagatagcccaggactttaagaccgatcttaggttccaatcctcagctgttatggcactccaggaggccagtgaagcttacctcgttgggttgtttgaagacaccaacctgtgcgccatccacgccaaaagggtgacaatcatgccgaaggatattcagctcgccagaagaatcagaggggagagggcttaa |
| pAG895 | HP1-RspA(C) | atggaggaggagtacgccgtggaaaagatcatcgacaggcgggtgcgcaagggaaaggtggagtactatctgaaatggaagggctatcccgaaactgagaacacgtgggagccggagaacaatctcgactgccaggatcttatccagcagtacgaggcgagccgcaaggattccggaggaggcggcagcggcggaggggggatccggcgacagcttctggatcttcgtgaagagactgtaa |
| pAG1015 | His-Tag-thrombin-TEV-FKBP-RspA(C) | atgggcagcagccatcatcatcatcatcacagcagcggcctggtgccgcgcggcagccatatggctagcgaaaacctgtattttcagggcggagtgcaggtggaaaccatctccccaggagacgggcgcaccttccccaagcgcggccagacctgcgtggtgcactacaccgggatgcttgaagatggaaagaaatttgattcctcccgggacagaaacaagccctttaagtttatgctaggcaagcaggaggtgatccgaggctgggaagaaggggttgcccagatgagtgtgggtcagagagccaaactgactatatctccagattatgcctatggtgccactgggcacccaggcatcatcccaccacatgccactctcgtcttcgatgtggagcttctaaaactggaagaatccggaggaggcggcagcggcggaggggggatccggcgacagcttctggatcttcgtgaagagactgtaa |
| pAG1016 | His-Tag-thrombin-TEV-FRB -RspA(N) | atgggcagcagccatcatcatcatcatcacagcagcggcctggtgccgcgcggcagccatatggctagcgaaaacctgtattttcagggcgagatgtggcatgaaggcctggaagaggcatctcgtttgtactttggggaaaggaacgtgaaaggcatgtttgaggtgctggagcccttgcatgctatgatggaacggggcccccagactctgaaggaaacatcctttaatcaggcctatggtcgagatttaatggaggcccaagagtggtgcaggaagtacatgaaatcagggaatgtcaaggacctcacccaagcctgggacctctattatcatgtgttccgacgaatctcaaagcaggtctccggaggaggcggcagcggcggaggggggatccatggagaccgtgagattcggcggcgacgacatcgagaacagcctggccaagatggacgacaaggccctggacaagctggccttcggcgccatccagctggacggcaacggcaagatcatccactacaacgccgccgagggcaccatcaccggcagagaccccaagaccgtgatcggcaagaacttcttcaccgacgtggcccccggcacccagagcaaggagttccagggcagattcaaggagggcgtgcagaagggcgacctgaacaccatgttcgagtggatgatccccaccagcagaggccccaccaaggtgaaggtgcacatgaagaaggccatgacctaa |
| pAG1017 | His-Tag-thrombin-TEV-FKBP-Hha(C) | atgggcagcagccatcatcatcatcatcacagcagcggcctggtgccgcgcggcagccatatggctagcgaaaacctgtattttcagggcggagtgcaggtggaaaccatctccccaggagacgggcgcaccttccccaagcgcggccagacctgcgtggtgcactacaccgggatgcttgaagatggaaagaaatttgattcctcccgggacagaaacaagccctttaagtttatgctaggcaagcaggaggtgatccgaggctgggaagaaggggttgcccagatgagtgtgggtcagagagccaaactgactatatctccagattatgcctatggtgccactgggcacccaggcatcatcccaccacatgccactctcgtcttcgatgtggagcttctaaaactggaagaatccggaggaggcggcagcggcggaggggggatccggtgacagctattgggtctttgtgaaacgggtgtaa |
| pAG1049 | HP1W45A-RspA(C) | atggaggaggagtacgccgtggaaaagatcatcgacaggcgggtgcgcaagggaaaggtggagtactatctgaaagccaagggctatcccgaaactgagaacacgtgggagccggagaacaatctcgactgccaggatcttatccagcagtacgaggcgagccgcaaggattccggaggaggcggcagcggcggaggggggatccggcgacagcttctggatcttcgtgaagagactgtaa |
| pAG1082 | Hha(C)-H3 | atgggtgacagctattgggtctttgtgaaacgggtgtccggaggaggcggcagcggcggaggggggatccatggcacgaaccaagcagactgcccgcaaatccaccggcggtaaggcaccacggaagcagctggctaccaaagctgctagaaagagtgctcctgcaacaggggggagtgaaaaagcctcaccgatatcgccctgggactgtggcccttagggagatcagaagataccaaaaaagcacagagttgctgatccggaagctgccatttcaacgactggtgcgggagatagcccaggactttaagaccgatcttaggttccaatcctcagctgttatggcactccaggaggccagtgaagcttacctcgttgggttgtttgaagacaccaacctgtgcgccatccacgccaaaagggtgacaatcatgccgaaggatattcagctcgccagaagaatcagaggggagagggcttaa |

| | | |
|---|---|---|
| pAG1118 | His-Tag-thrombin-TEV-FRB -Hha(N) | atgggcagcagccatcatcatcatcatcacagcagcggcctggtgccgcgcggcagccatatggctagcgaaaacctgtattttcagggcgagatgtggcatgaaggcctggaagaggcatctcgtttgtactttggggaaaggaacgtgaaaggcatgtttgaggtgctggagcccttgcatgctatgatggaacggggcccccagactctgaaggaaacatcctttaatcaggcctatggtcgagatttaatggaggcccaagagtggtgcaggaagtacatgaaatcagggaatgtcaaggacctcacccaagcctgggacctctattatcatgtgttccgacgaatctcaaagcaggtctccggaggaggcggcagcggcggagggggatccatggagcatgttgcctttggcagtgaggacatcgagaacactctggccaaaatggacgacggacaactggatgggttggcctttggcgcaattcagctcgatggtgacgggaatatcctgcagtacaatgctgctgaaggagacatcacaggcagagatcccaaacaggtgattgggaagaacttcttcaaggatgttgcacctggaacggattctccgagttttacggcaaattcaaggaaggcgtagcgtcagggaatctgaacaccatgttcgaatggatgataccgacaagcaggggaccaaccaaggtcaaggtgcacatgaagaaagcccttttcctaa |
| pAG1143 | RspA(N)-MEK1- C-myc | atggagaccgtgagattcggcggcgacgacatcgagaacagcctggccaagatggacgacaaggccctggacaagctggccttcggcgccatccagctggacggcaacggcaagatcatccactacaacgccgccgagggcaccatcaccggcagagaccccaagaccgtgatcggcaagaacttcttcaccgacgtggcccccggcacccagagcaaggagttccagggcagattcaaggagggcgtgcagaagggcgacctgaacaccatgttcgagtggatgatccccaccagcagaggccccaccaaggtgaaggtgcacatgaagaaggccatgacctccggaggaggcggcagcggcggagggggatccatgcccaagaagaagccgacgcccatccagctgaacccggcccccgacggctctgcagttaacgggaccagctctgcggagaccaacttggaggccttgcagaagaagctggaggagctagagcttgatgagcagcagcgaaagcgccttgaggcctttcttacccagaagcagaaggtgggagaactgaaggatgacgactttgagaagatcagtgagctgggggctggcaatggcggtgtggtgttcaaggtctcccacaagccttctggcctggtcatggccagaaagctaattcatctggagatcaaacccgcaatccggaaccagatcataagggagctgcaggttctgcatgagtgcaactctccgtacatcgtgggcttctatggtgcgttctacagcgatggcgagatcagtatctgcatggagcacatggatggaggttctctggatcaagtcctgaagaaagctggaagaattcctgaacaaattttaggaaaagttagcattgctgtaataaaaggcctgacatatctgagggagaagcacaagatcatgcacagagatgtcaagccctccaacatcctagtcaactcccgtggggagatcaagctctgtgactttggggtcagcgggcagctcatcgactccatggccaactccttcgtgggcacaaggtcctacatgtcgccagaaagactccaggggactcattactctgtgcagtcagacatctggagcatgggactgtctctggtagagatggcggttgggaggtatcccatccctcctccagatgccaaggagctggagctgatgtttgggtgccaggtggaaggagatgcggctgagaccccacccaggccaaggacccccgggaggccccttagctcatacggaatggacagccgacctcccatggcaatttttgagttgttggattacatagtcaacgagcctcctccaaaactgcccagtggagtgttcagtctggaatttcaagattttgtgaataaatgcttaataaaaaaccccgcagagagagcagatttgaagcaactcatggttcatgcttttatcaagagatctgatgctgaggaagtggattttgcaggttggctctgctccaccatcggccttaaccagcccagcacaccaacccatgctgctggcgtcgaacaaaagcttatttctgaagaggacttgtaa |
| pAG1144 | Lyn11- FRB-RspA(N) | atgggctgcatcaagtccaagggcaaggactccgccgaacaaaagcttatttctgaagaggacttggaattcgagatgtggcatgaaggcctggaagaggcatctcgtttgtactttggggaaaggaacgtgaaaggcatgtttgaggtgctggagcccttgcatgctatgatggaacggggcccccagactctgaaggaaacatcctttaatcaggcctatggtcgagatttaatggaggcccaagagtggtgcaggaagtacatgaaatcagggaatgtcaaggacctcacccaagcctgggacctctattatcatgtgttccgacgaatctcaaagcaggtctccggaggaggcggcagcggcggagggggatccatggagaccgtgagattcggcggcgacgacatcgagaacagcctggccaagatggacgacaaggccctggacaagctggccttcggcgccatccagctggacggcaacggcaagatcatccactacaacgccgccgagggcaccatcaccggcagagaccccaagaccgtgatcggcaagaacttcttcaccgacgtggcccccggcacccagagcaaggagttccagggcagattcaaggagggcgtgcagaagggcgacctgaacaccatgttcgagtggatgatccccaccagcagaggccccaccaaggtgaaggtgcacatgaagaaggccatgacctaa |

| | | |
|---|---|---|
| pAG1175 | H2B-mCherry-Hha(C) | atgcccgaacctgcgaagtcagcgcccgctcccaaaaaaggctctaaaaagctgtcgcc<br>aagacccagaagaaggggataagaaaaggcgtaagaccaggaaagagagttacgc<br>catttacgtgtacaaagtactaaaacaagtccacccggacactggcatctcctcaaaggcg<br>atgggcattatgaactcatttgtaaacgacatcttcgagcgcatcgccggagaagcgtcgcg<br>cctggcgcattacaacaagcgctccactatcacatcccgggagatccagacggccgtgcg<br>cctgctcctgcccggagaactggccaaacacgctgtgtctgagggcacaaaggccgtgac<br>caagtacaccagctccaagggcggaggctccggaggcggatccatgggtgagcaagggc<br>gaggaggataacatggccatcatcaaggagttcatgcgcttcaaggtgcacatggagggct<br>ccgtgaacggccacgagttcgagatcgagggcgagggcgagggccgcccctacgaggg<br>cacccagaccgccaagctgaaggtgaccaagggtggcccctgcccttcgcctgggacat<br>cctgtcccctcagttcatgtacggctccaaggcctacgtgaagcaccccgccgacatcccg<br>actacttgaagctgtccttccccgagggcttcaagtgggagcgcgtgatgaacttcgaggac<br>ggcggcgtggtgaccgtgacccaggactcctccctgcaggacggcgagttcatctacaagg<br>tgaagctgcgcggcaccaacttcccctccgacggccccgtaatgcagaagaagaccatgg<br>gctgggaggcctcctccgagcggatgtaccccgaggacggcgccctgaagggcgagatc<br>aagcagaggctgaagctgaaggacggcggccactacgacgctgaggtcaagaccacct<br>acaaggccaagaagcccgtgcagctgcccggcgcctacaacgtcaacatcaagttggac<br>atcacctcccacaacgaggactacaccatcgtggaacagtacgaacgcgccgagggccg<br>ccactccaccggcggcatggacgagctgtacaagaggaggttctacggccccgtgggtga<br>cagctattgggtctttgtgaaacgggtgtaa |
| pAG1176 | C-myc-mCherry-ERK2-RspA(C) | atggaacaaaagcttatttctgaagaggacttggtgagcaagggcgaggaggataacatg<br>gccatcatcaaggagttcatgcgcttcaaggtgcacatggagggctccgtgaacggccacg<br>agttcgagatcgagggcgagggcgagggccgcccctacgagggcacccagaccgccaa<br>gctgaaggtgaccaagggtggcccctgcccttcgcctgggacatcctgtcccctcagttcat<br>gtacggctccaaggcctacgtgaagcaccccgccgacatccccgactacttgaagctgtcct<br>tccccgagggcttcaagtgggagcgcgtgatgaacttcgaggacggcggcgtggtgaccgt<br>gacccaggactcctccctgcaggacggcgagttcatctacaaggtgaagctgcgcggcac<br>caacttcccctccgacggccccgtaatgcagaagaagaccatgggctgggaggcctcctc<br>cgagcggatgtaccccgaggacggcgccctgaagggcgagatcaagcagaggctgaag<br>ctgaaggacggcggccactacgacgctgaggtcaagaccacctacaaggccaagaagc<br>ccgtgcagctgcccggcgcctacaacgtcaacatcaagttggacatcacctcccacaacga<br>ggactacaccatcgtggaacagtacgaacgcgccgagggccgccactccaccggcggc<br>atggacgagctgtacaagtccggactcagatctcgagctcaagcttcgaattctgcagtcatg<br>gcggcggcggcggcggcgggcgcgggcccggagatggtccgcgggcaggtgttcgacg<br>tggggccgcgctacaccaacctctcgtacatcggcgagggcgcctacggcatggtgtgctct<br>gcttatgataatgtcaacaaagttcgagtagctatcaagaaaatcagcccctttgagcaccag<br>acctactgccagagaaccctgagggagataaaaatcttactgcgcttcagacatgagaaca<br>tcattggaatcaatgacattattcgagcaccaaccatcgagcaaatgaaagatgtatatatag<br>tacaggacctcatggaaacagatctttacaagctcttgaagacacaacacctcagcaatga<br>ccatatctgctatttctctaccagatcctcagagggttaaaatatatccattcagctaacgttctg<br>caccgtgacctcaagccttccaacctgctgctcaacaccacctgtgatctcaagatctgtgact<br>ttggcctggcccgtgttgcagatccagaccatgatcacacagggttcctgacagaatatgtgg<br>ccacacgttggtacagggctccagaaattatgttgaattccaagggctacaccaagtccattg<br>atatttggtctgtaggctgcattctggcagaaatgctttctaacaggcccatctttccagggaag<br>cattatcttgaccagctgaaccacattttgggtattcttggatccccatcacaagaagacctga<br>attgtataataaatttaaaagctaggaactatttgctttctcttccacacaaaaataaggtgccat<br>ggaacaggctgttcccaaatgctgactccaaagctctggacttattggacaaaatgttgacatt<br>caacccacacaagaggattgaagtagaacaggctctggcccacccatatctggagcagta<br>ttacgacccgagtgacgagcccatcgccgaagcaccattcaagttcgacatggaattggat<br>gacttgcctaaggaaaagctcaaagaactaatttttgaagagactgctagattccagccagg<br>atacagatcttccggaggaggcggcagcggcggagggggatccggcgacagcttctggat<br>cttcgtgaagagactgtaa |
| pAG1177 | C-myc-FKBP-RspA(C) | atggaacaaaagcttatttctgaagaggacttggaattcggagtgcaggtggaaaccatctc<br>cccaggagacgggcgcaccttccccaagcgcggccagacctgcgtggtgcactacaccg<br>ggatgcttgaagatggaaagaaatttgattcctcccgggacagaaacaagccctttaagttta<br>tgctaggcaagcaggaggtgatccgaggctgggaagaaggggttgcccagatgagtgtg<br>ggtcagagagccaaactgactatatctccagattatgcctatggtgccactgggcacccagg<br>catcatcccaccacatgccactctcgtcttcgatgtggagcttctaaaactggaagaatccgg<br>aggaggcggcagcggcggagggggatccggcgacagcttctggatcttcgtgaagagac<br>tgtaa |
